# Inhibition of DHODH Activates Pyroptosis and cGAS-STING Pathways to Enhance NK cell Infiltration Mediated Anti-tumor Immunity in Melanoma

**DOI:** 10.1101/2025.03.20.644471

**Authors:** Yongrui Hai, Ruizhuo Lin, Weike Liao, Shuo Fu, Renming Fan, Guiquan Ding, Junyan Zhuang, Bingjie Zhang, Yi Liu, Junke Song, Gaofei Wei

**Author notes:** **Corresponding authors:** (Yi Liu); (Junke Song); (Gaofei Wei). These authors contributed equally to this work (Yongrui Hai, Ruizhuo Lin, Weike Liao).

## Abstract

Cancer cells are heavily reliant on *de novo* pyrimidine synthesis. Suppression of pyrimidine metabolism directly inhibits tumor growth and fosters immune activation within the tumor microenvironment. Dihydroorotate dehydrogenase (DHODH), a crucial enzyme governing *de novo* pyrimidine synthesis, is a critical player in this context. Inhibition of DHODH not only reverses immunosuppression but also instigates a mild innate immune response. However, the impact of DHODH inhibition on natural killer (NK) cells remains unexplored. In this study, we found that inhibition of DHODH efficiently promotes NK cells infiltration in tumors. Suppression of DHODH led to increased oxidative stress in mitochondria, the release of mtDNA, and activation of caspase 3, which in turn activated the cGAS-STING pathway and pyroptosis in cancer cells, respectively, contributing to NK cells induced antitumor immune responses in melanoma. Additionally, we developed EA6, a novel DHODH inhibitor with higher efficacy in promoting NK cells infiltration. In summary, this study underscores that modulation of pyrimidine metabolism can effectively trigger antitumor immune responses, with a specific emphasis on NK cells. This finding opens new avenues for enhancing the efficacy of targeted nucleotide metabolism in cancer therapy.

**GRAPHICAL ABSTRACT:** 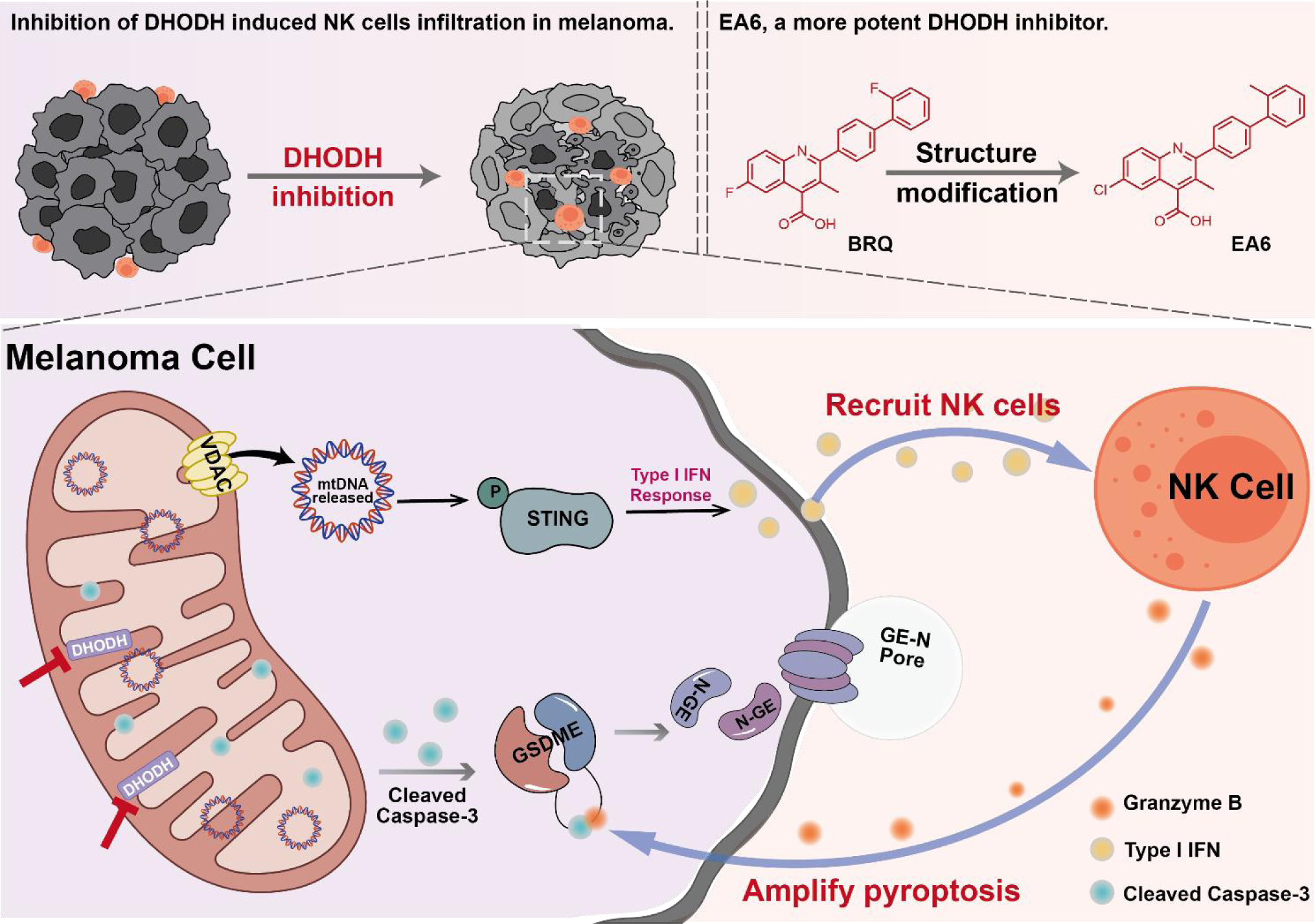

**The anti-tumor mechanisms of DHODH inhibition.** Inhibition of DHODH activates cGAS-STING pathways to enhance NK cell infiltration. And the tumor-infiltrating NK cells facilitate melanoma cells pyroptosis which providing a positive feedback mechanism for DHODH-mediated anti-tumor immunity.

## INTRODUCTION

Abnormally proliferating cancer cells exhibit a heightened demand for energy and biomacromolecules [1]. Nucleotides, the fundamental components of RNA and DNA, play a crucial role in the accelerated nucleotide metabolism that drives tumor cell proliferation, metastasis, and immune evasion [2]. Consequently, targeting nucleotide synthesis remains a cornerstone of contemporary cancer therapy. Traditional nucleotide synthesis inhibitors, such as 5-fluorouracil (5-FU), methotrexate, and cytarabine, are analogs of tumor nucleotide metabolites. Recent research has revealed that nucleotide antimetabolite drugs (NMDs) not only restrain tumor cell proliferation but also hinder immune evasion, providing novel insights into therapeutic approaches aimed at nucleotide metabolism [3].

Pyrimidine nucleotides, a key component among nucleotides, are synthesized through two distinct pathways: the salvage pathway and the *de novo* pathway. Terminally differentiated or resting cells primarily rely on the salvage pathway to maintain nucleotide homeostasis, whereas rapidly proliferating cells, particularly cancer cells, depend heavily on *de novo* pyrimidine synthesis [4]. Thus, effectively suppressing *de novo* pyrimidine synthesis has proven instrumental in impeding tumor progression. DHODH, a critical rate-limiting enzyme in the *de novo* pyrimidine synthesis pathway, catalyzes the conversion of dihydroorotate to orotate. Unique among pyrimidine synthetases, DHODH is located on the outer surface of the inner mitochondrial membrane and is linked to the mitochondrial electron transport chain [5]. Inhibition of DHODH has demonstrated potent efficacy in suppressing the growth of melanoma [6], glioma [7], small cell lung cancer [8], and various solid tumors, while simultaneously overcoming differentiation blockade in acute myeloid leukemia [9]. These findings emphasize the potential of DHODH as a promising pharmaceutical target in cancer treatment.

Notably, pyrimidine metabolism significantly influences the immune response within the tumor microenvironment. Disruptions in purine and pyrimidine pools can lead to increased genomic instability, enhancing immunogenicity [3]. Hans-Georg Sprenger et al. have suggested that imbalanced pyrimidine levels in cells can trigger innate immune responses [10]. Conversely, pyrimidine metabolism is closely associated with anti-tumor effects of various immune cells. DHODH inhibition restrains cell proliferation without affecting T cell functionality [11]. It also promotes the differentiation of myeloid-derived suppressor cells (MDSC), which can reverse immune-suppressive microenvironments [12]. Additionally, pyrimidines released from macrophages limit gemcitabine therapy in pancreatic cancer [13], indicating that blocking pyrimidine uptake by cancer cells can enhance the therapeutic efficiency of chemotherapeutic agents. Thus, the perturbation of pyrimidine balance holds the potential to augment antitumor immunity.

NK cells are pivotal roles in innate immunosurveillance of cancer by identifying and attacking stressed cells, particularly those with suppressed major histocompatibility complex class I (MHC-I) expression [14, 15]. As MHC-unrestricted NK cells complement T cell-mediated tumor immunity, exploring the impact of pyrimidine imbalance on NK cells activation is of significant interest.

In this study, we observed a substantial increase in tumor-infiltrating NK cells following DHODH inhibition. The suppression of DHODH led to cancer cells pyroptosis and activated the cGAS-STING pathway, which serves as the foundation for NK cells-induced anti-tumor immune responses. Additionally, we designed and synthesized a series of DHODH inhibitors, with EA6 exhibiting the highest activity. EA6 emerges as a markedly more efficient DHODH inhibitor, boasting approximately a 4-fold improvement in activity compared to the classical DHODH inhibitor brequniar (BRQ) and trigger NK infiltration more efficiently. In summary, this study underscores that modulation of pyrimidine metabolism can effectively trigger antitumor immune responses, with a specific emphasis on NK cells. This finding opens new avenues for enhancing the efficacy of targeted nucleotide metabolism in cancer therapy.

## RESULTS

### Overexpression of DHODH relates to immunosuppressive in melanoma

To elucidate the role of DHODH in tumors, we investigated DHODH expression in various cancers using clinical data from TCGA. Our analysis revealed that DHODH is highly expressed in multiple cancers, including skin cutaneous melanoma (SKCM) (Fig. 1A). In the SKCM samples, DHODH expression displayed a significant increase at all stages (I-IV; n = 77, 140, 171, 24), compared to normal skin tissue (n = 1809), although no significant differences were observed in DHODH levels among different tumor grades (Fig. 1B).

**Fig. 1.**
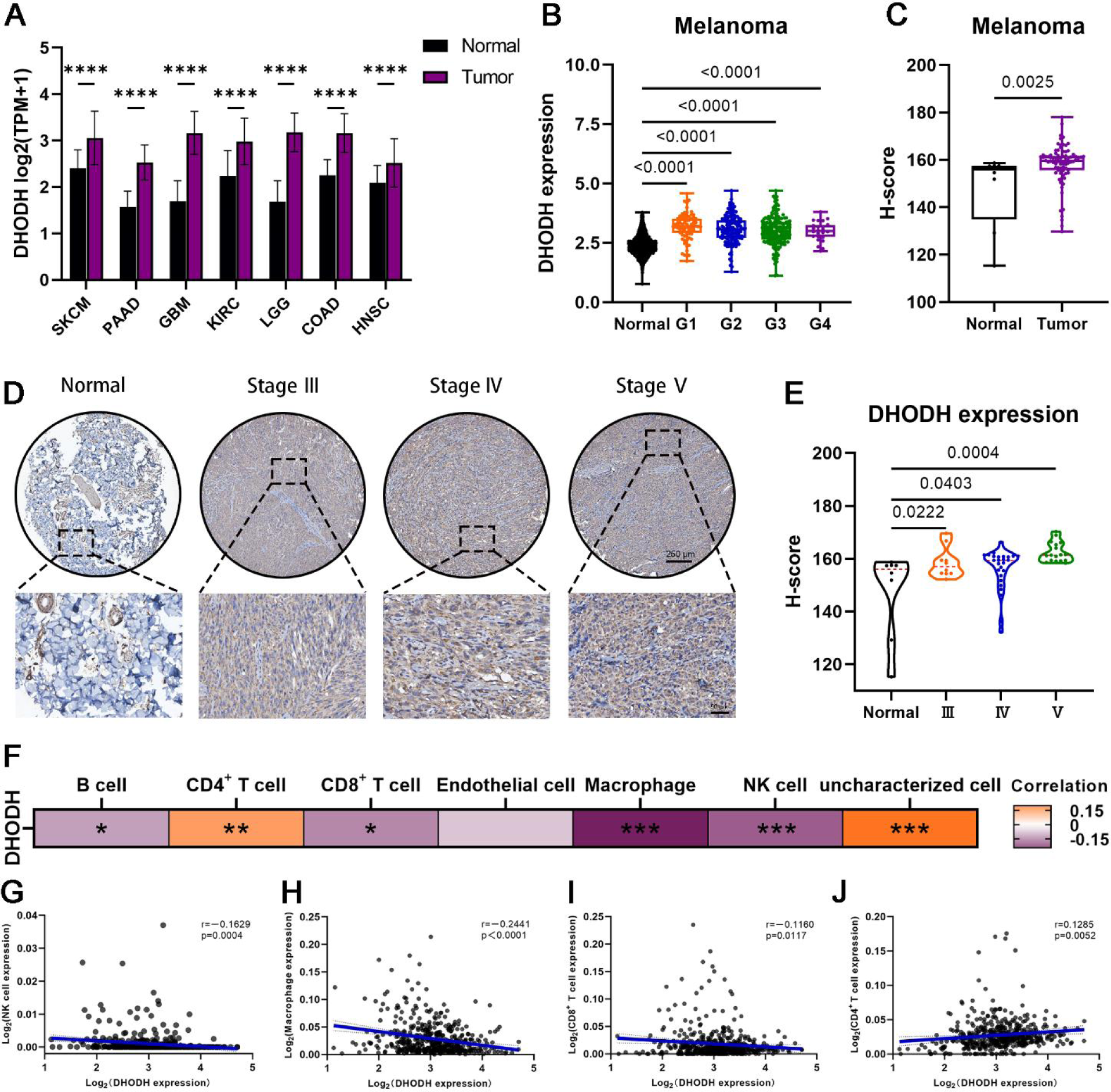
Over expressed DHODH relates to immunosuppressive in melanoma. (A) DHODH expression levels in different tumors; two-way ANOVA. (B) DHODH levels among different grades in melanoma; one way ANOVA. Dates were obtained from TCGA. (C) Histochemistry score(H-score) of DHODH in normal tissues (n = 8) and melanoma (n = 85); two-tailed Student’s t-test. (D) Representative images of immunohistochemical staining of DHODH. Melanoma staging by Clark classification. (E) Immunohistochemistry score of DHODH in normal tissues (n = 8) and different grades tissue samples of melanoma (n = 10; 26; 17); one way ANOVA. (F) Correlation between DHODH expression and immune cell infiltration. Dates were obtained from TCGA. (G-J) Plots of NK cells, macrophage, CD8^+^ T cells and CD4^+^ T cells versus DHODH expression in melanoma.

Immunohistochemistry was performed on melanoma tissue microarrays to evaluate DHODH expression, with a full view of the melanoma tissue microarray showed in Fig. S1A. Consistently, DHODH was upregulated in melanoma tissues relative to normal skin tissues (Fig. 1C) and elevated DHODH expression was observed across various melanoma stages (Fig. 1D-E). Notably, inhibition of DHODH resulted in a significant regression of melanoma growth [16]. These compelling findings underscore the potential of DHODH as a promising therapeutic target for melanoma treatment.

Furthermore, prior research has demonstrated that cellular pyrimidine imbalance has been shown to effectively trigger innate immunity [10] which prompted us to explore a potential correlation between DHODH expression and the immune response. To investigate this relationship, the computational method EPIC was employed to estimate the proportions of immune and cancer cells within the tumor microenvironment. As anticipated, DHODH expression levels showed a negative correlation with the presence of CD8^+^ T cells, NK cells, B cells, and macrophages (Fig. 1F-J). These findings collectively suggest that high levels of DHODH are associated with immunosuppression in melanoma.

These results provide compelling evidence that DHODH inhibition could serve as an effective therapeutic strategy against melanoma by promoting immune activation. To further investigate the effects of DHODH inhibition, we conducted experiments using A375 and B16F10 cell lines, which are known for their high DHODH expression levels (Fig. S1B-C).

### BRQ induced NK cells infiltration in tumor

BRQ is a well-known DHODH inhibitor that effectively impedes *de novo* pyrimidines synthesis and has been widely utilized in tumor treatment. To assess the antitumor efficacy of BRQ, we conducted experiments using the B16F10 melanoma model in C57BL/6 mice, as outlined in Fig. 2A. BRQ significantly suppressed tumor growth, as illustrated in Fig. 2B-D.

**Fig. 2.**
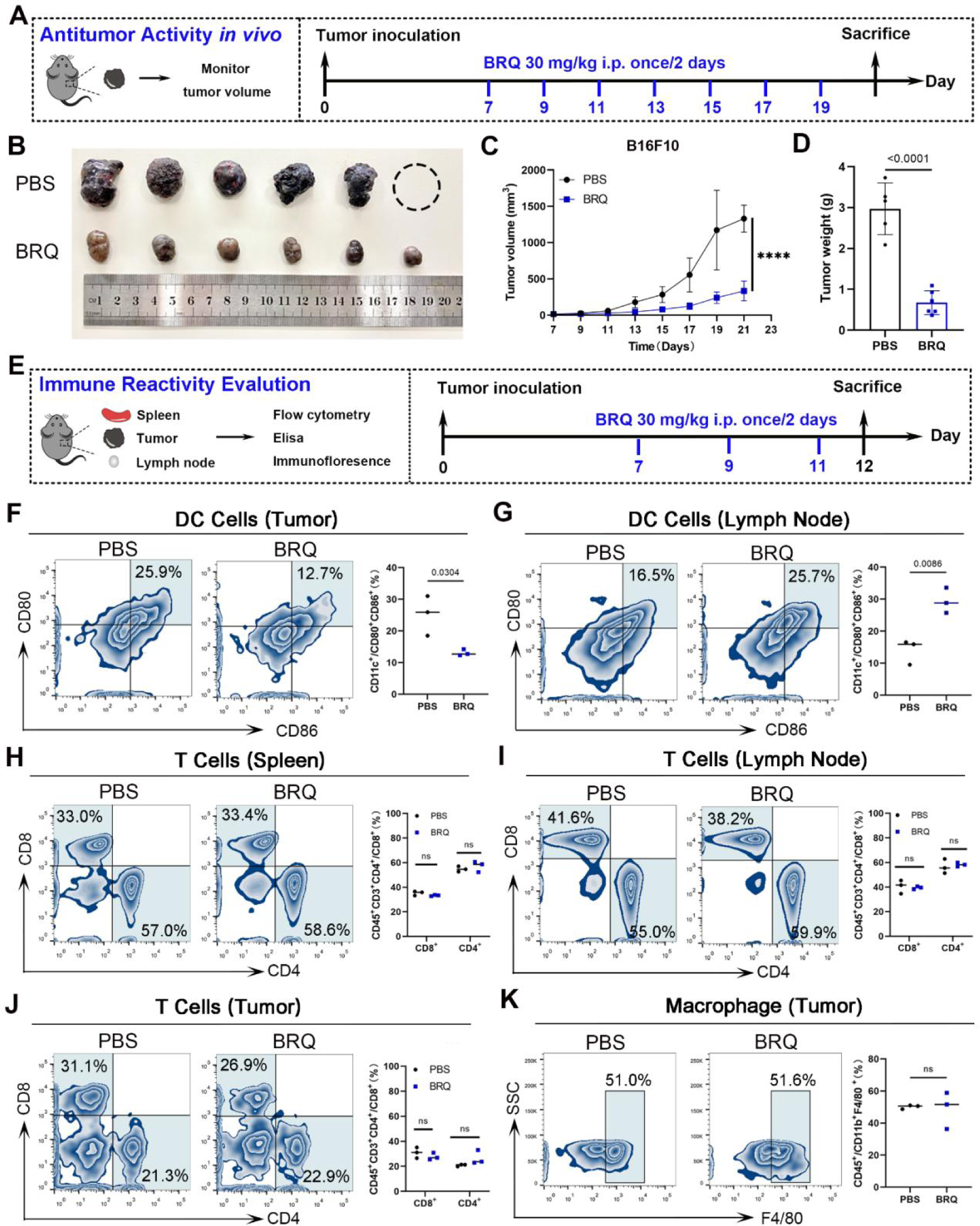
BRQ restrains melanoma growth. (A) Experiment scheme for BRQ therapy of the B16F10 tumor bearing C57BL/6 female mouse model. (B) Images of isolated tumors at day 21 for each group. (C) Tumor growth in the indicted groups of mice (n = 6; Mean ± SD). (D) Average tumor weight at Day 21 of each group. two-tailed Student’s t-test. (E) Immune experiment scheme of the B16F10 tumor bearing C57BL/6 female mouse model. (F-G) Representative flow cytometry images and quantitative graph of DC cells in (F) tumor and (G) lymph node. (H-J) Representative flow cytometry images and quantitative graph of T cells in (H) spleen, (I) lymph nodes and (J) tumor. (K) Representative flow cytometry images and quantitative graph of macrophage in tumor.

To explore the relationship between BRQ-mediated antitumor activity and immune activation, we performed comprehensive analyses of immune cell infiltration. Following three doses of BRQ treatment, tumors, spleens, and lymph nodes were subjected to flow cytometry, immunofluorescence, and immunohistochemistry analyses (Fig. 2E). The results demonstrated a decrease in mature dendritics cells (DCs) (CD11c^+^ CD80^+^ CD86^+^) within the tumor (Fig. 2F), while a notable increase was observed in the lymph nodes in response to BRQ treatment (Fig. 2G). The migration of mature DCs from the tumor to the lymph nodes can activate T cells[17]. However, the proportion of CD8^+^ T and CD4^+^ T cells in the spleens and lymph nodes remained unaltered following BRQ treatment (Fig. 2H-I), and there was no significant difference in tumor-infiltrating CD8^+^ T or CD4^+^ T cells between the PBS and BRQ therapy groups (Fig. 2J). Similarly, Macrophages are also an important component of immune system, tumor-infiltrating macrophages affect tumor cells proliferation, invasion and metastasis[18]. Nevertheless, the populations of macrophages showed no significant difference within tumors following BRQ treatment (Fig. 2K).

NK cells are critical for broad-spectrum antitumor effects, as they directly recognize and attack tumor cells without MHC restriction. This makes them particularly effective against tumors with downregulated or lost MHC-I expression, such as malignant melanomas [15, 19]. To determine the role of NK cells in BRQ-mediated tumor suppression, we analyzed NK cells infiltration using flow cytometry and Immunofluorescence. We found a significant increase in NK cells within tumors after BRQ treatment (Fig. 3A-B). Immunofluorescence analysis revealed that NK cells were less abundant and primarily located at the periphery of tumors in the PBS-treated group, whereas BRQ treatment facilitated deeper infiltration of NK cells into the tumor (Fig. 3C). Additionally, BRQ treatment upregulated the expression of pro-inflammatory cytokines, such as IFN-γ and IL-1β (Fig. S2A-B), while downregulating TGF-β, an immune suppressor known to inhibit NK cell activity (Fig. S2C) [20]. These results highlight that BRQ treatment enhances NK cells infiltration into the tumor microenvironment (Fig. 3D).

**Fig. 3.**
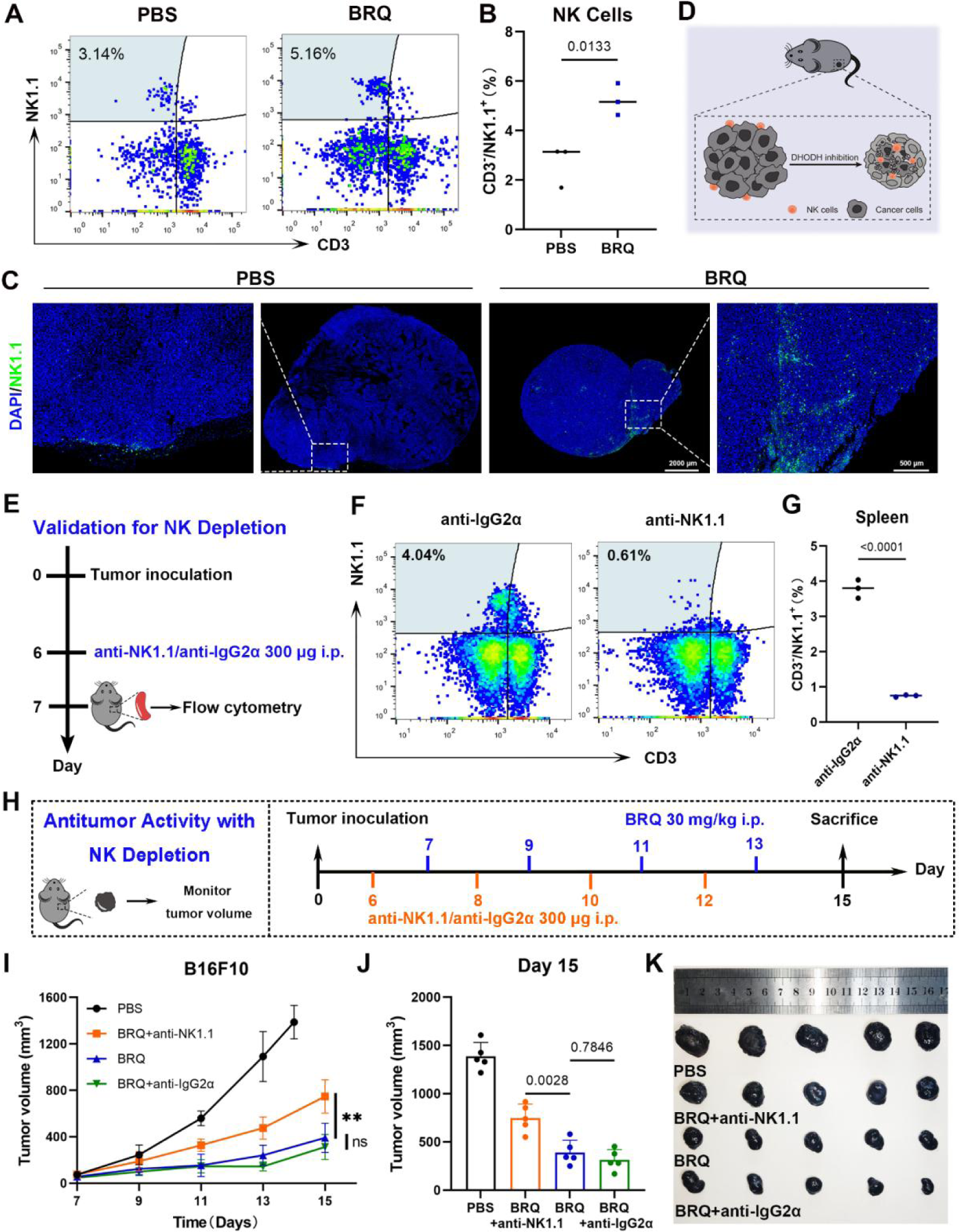
BRQ triggers NK cells infiltration in tumor. (A-B) Representative flow cytometry images and quantitative graph of NK cells in tumor. (C) Representative immunofluorescence images of tumor-infiltrating NK cells. (D) Schematic representation of BRQ induced antitumor immune response. (E) Schematic representation of NK depletion experiment. (F-G) Representative flow cytometry images and quantification of NK cells in spleen after treatment with anti-IgG2α or anti-NK1.1 antibody. (H) Experiment scheme for BRQ therapy with NK depletion. (I) Tumor growth curves in the various groups of mice (n = 5; mean ± SD). (J)Average tumor volume at Day 15 of each group. Two-tailed Student’s t-test. (K) Images of isolated tumors at the end of experiment in NK depletion experiment.

To assess the critical role of NK cells in BRQ-mediated antitumor effects, we used an anti-NK1.1 antibody to deplete NK cells in C57BL/6 mice, with an anti-IgG2α antibody serving as a control. NK cell depletion was confirmed through flow cytometry analysis of spleens 24 hours after the initial anti-NK1.1 antibody injection (Fig. 3E-G). We then evaluated the impact of NK cells depletion on BRQ-mediated antitumor activity, as outlined in Fig. 3H. Remarkably, the antitumor effects of BRQ were partially attenuated in the absence of NK cells, while the anti-IgG2α antibody had no effect on BRQ efficacy (Fig. 3I-K).

These findings underscore the essential role of NK cells in the efficacy of BRQ treatment. Collectively, our results demonstrate that BRQ effectively inhibits melanoma growth, activates the immune system, and relies on NK cells as key effectors.

### RNA-Seq analysis of B16F10 cells treated with BRQ

To elucidate the mechanistic basis underlying the enhanced infiltration of tumor-associated NK cells following BRQ treatment, we performed RNA-seq gene expression analysis. The results indicated high reproducibility among samples within the same group (Fig. S3A), and principal component analysis (PCA) revealed a significant distinction between the BRQ-treated and control groups (Fig. S3B). Notably, BRQ treatment led to the up-regulation of 2,460 genes and down-regulation of 3,518 genes (Fig. S3C-D).

Differentially expressed genes (DEGs) were analyzed using Kyoto Encyclopedia of Genes and Genomes (KEGG) and Gene Ontology (GO) enrichment analyses. KEGG analysis revealed significant enrichment of DEGs in pathways related to pyrimidine and purine metabolism, reflecting BRQ’s inhibitory effect on *de novo* pyrimidine synthesis. Additionally, DEGs were enriched in pathways associated with immune responses, including the Rap1 signaling pathway, NOD-like receptor signaling pathway, JAK-STAT signaling pathway, and PD-L1 expression and PD-1 checkpoint pathway. DEGs also showed enrichment in cytokine-related pathways, such as the TNF signaling pathway, IL-17 signaling pathway, and TGF-β signaling pathway. Notably, many DEGs were linked to NK cell-mediated cytotoxicity (Fig. 4A).

**Fig. 4.**
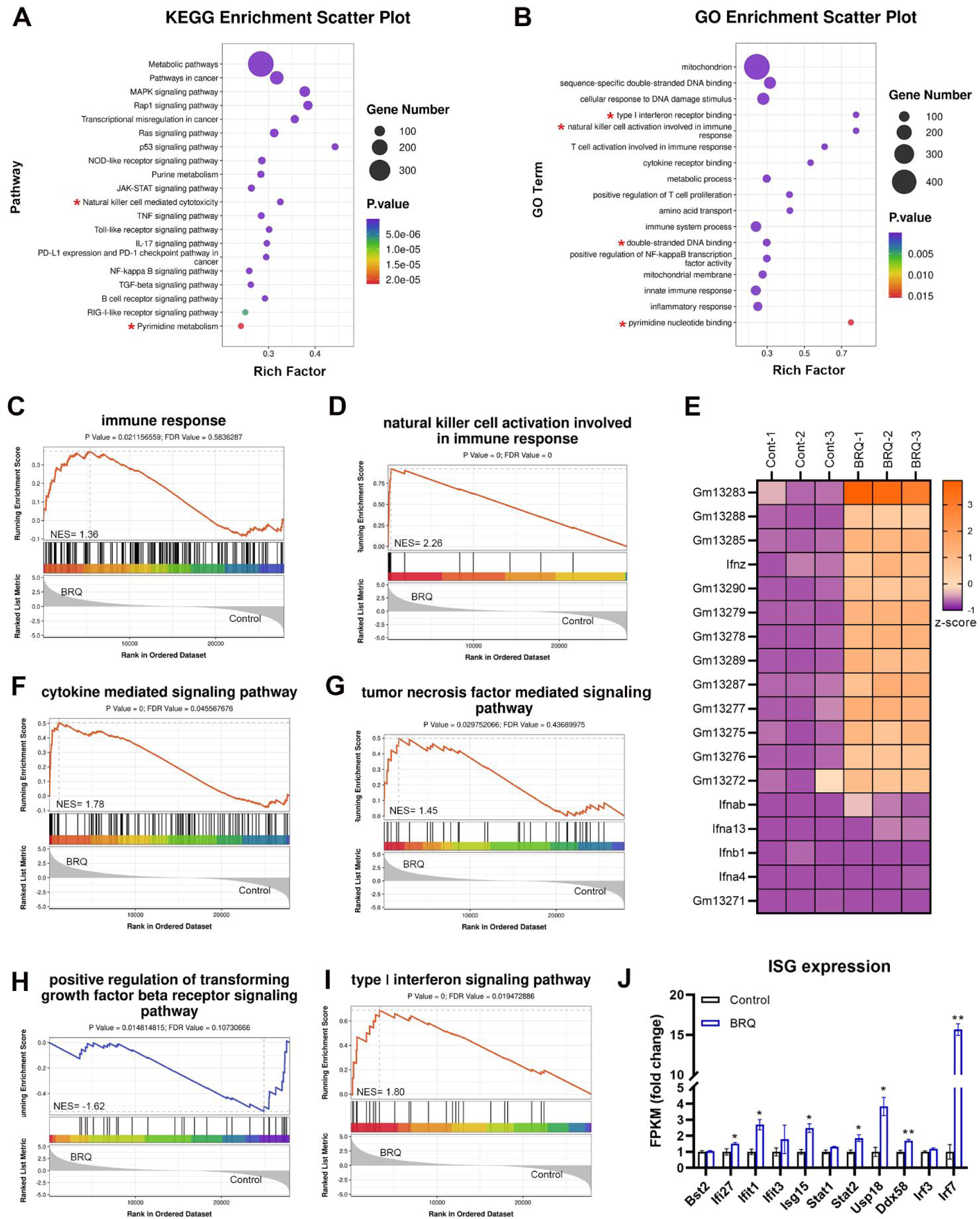
RNA-Seq analysis of B16F10 cells treated with BRQ. (A) KEGG enrichment analysis of DEGs after BRQ treatment. (B) GO enrichment analysis of DEGs after BRQ treatment. (C-D) GSEA analysis of (C) immune response and (D) NK cells activation. (E) Specific genes related to NK cells activation. (F-I) GSEA analysis of (F) cytokine, (G) TNF, (H) TGF-β and (I) IFN-Ⅰ mediated signaling pathway. (J) ISG expression between the BRQ group and the control group.

GO analysis further highlighted the impact of BRQ treatment on pyrimidine metabolism and immune responses. Specifically, BRQ treatment was found to affect mitochondrial function and double-stranded DNA binding, which could contribute to immune activation (Fig. 4B).

Gene Set Enrichment Analysis (GSEA) revealed that genes upregulated by BRQ were significantly enriched in pathways related to the immune system and NK cell activation (Fig. 4C-D), consistent with our in vivo findings. Fig. 4E shows a marked increase in the expression of genes associated with NK cell activation following BRQ treatment. Additionally, BRQ treatment activated the cytokine-mediated signaling pathway (Fig. 4F). Importantly, BRQ upregulated the TNF-α signaling pathway (Fig. 4G) and downregulated the TGF-β signaling pathway (Fig. 4H), corroborating its antitumor effects. The activation of the type I interferon (IFN-Ⅰ) signaling pathway was also observed (Fig. 4I), supported by significant up-regulation of interferon-stimulated genes (ISGs) such as *IFIT1*, *ISG15*, *STAT2*, and *IRF7* [21, 22] (Fig. 4J). This suggests that BRQ may activate immune responses through the IFN-Ⅰ signaling pathway.

Considering the fundamental role of mitochondria in innate immunity [22] and the potential of cellular pyrimidine imbalance to trigger mitochondrial DNA-dependent innate immunity [10], our RNA-seq results support the hypothesis that BRQ-induced pyrimidine imbalance may contribute to NK cells infiltration into tumors by activating the STING signaling pathway.

### BRQ activates cGAS-STING pathway to enhance the antitumor immunity of NK cells

The STING signaling pathway serves as a pivotal link bridging innate and adaptive immunity [23, 24]. Stimulation of the STING pathway can activate NK cells, promising effectors for antitumor responses [25, 26]. STING can be activated in response to DNA damage, with the recognition of intracellular free DNA by cGAS being the initial step. Notably, inhibition of *de novo* pyrimidine synthesis efficiently induces DNA damage and disrupts the pyrimidine balance [4]. Consequently, we explored the effect of BRQ on activating the STING pathway.

Initially, the A375 and B16F10 cell lines were treated with BRQ, followed by the assessment of DNA and mitochondria co-localization through immunofluorescence staining. Our observations revealed a dose-dependent release of mitochondrial DNA (mtDNA) from mitochondria following BRQ treatment in both B16F10 and A375 cells (Fig. 5A). Subsequently, we examined the expression levels of key proteins involved in the cGAS-STING signaling pathway. BRQ led to significant up-regulation of cGAS, p-STING and p-TBK1 in both A375 and B16F10 cells (Fig. 5B). Moreover, immunofluorescent staining showed a notable increase in the expression of p-STING following BRQ treatment (Fig. S4A). The activation of cGAS-STING signaling pathway leads to the induction of type I interferons (IFN-I), which paly critical roles in NK cell biology, including maturation, homeostasis, and activation [27]. To evaluate this, we assessed IFN-β expression in the cell supernatant. As expected, BRQ treatment resulted in a dose-dependent increase in IFN-β expression (Fig. 5C).

**Fig. 5.**
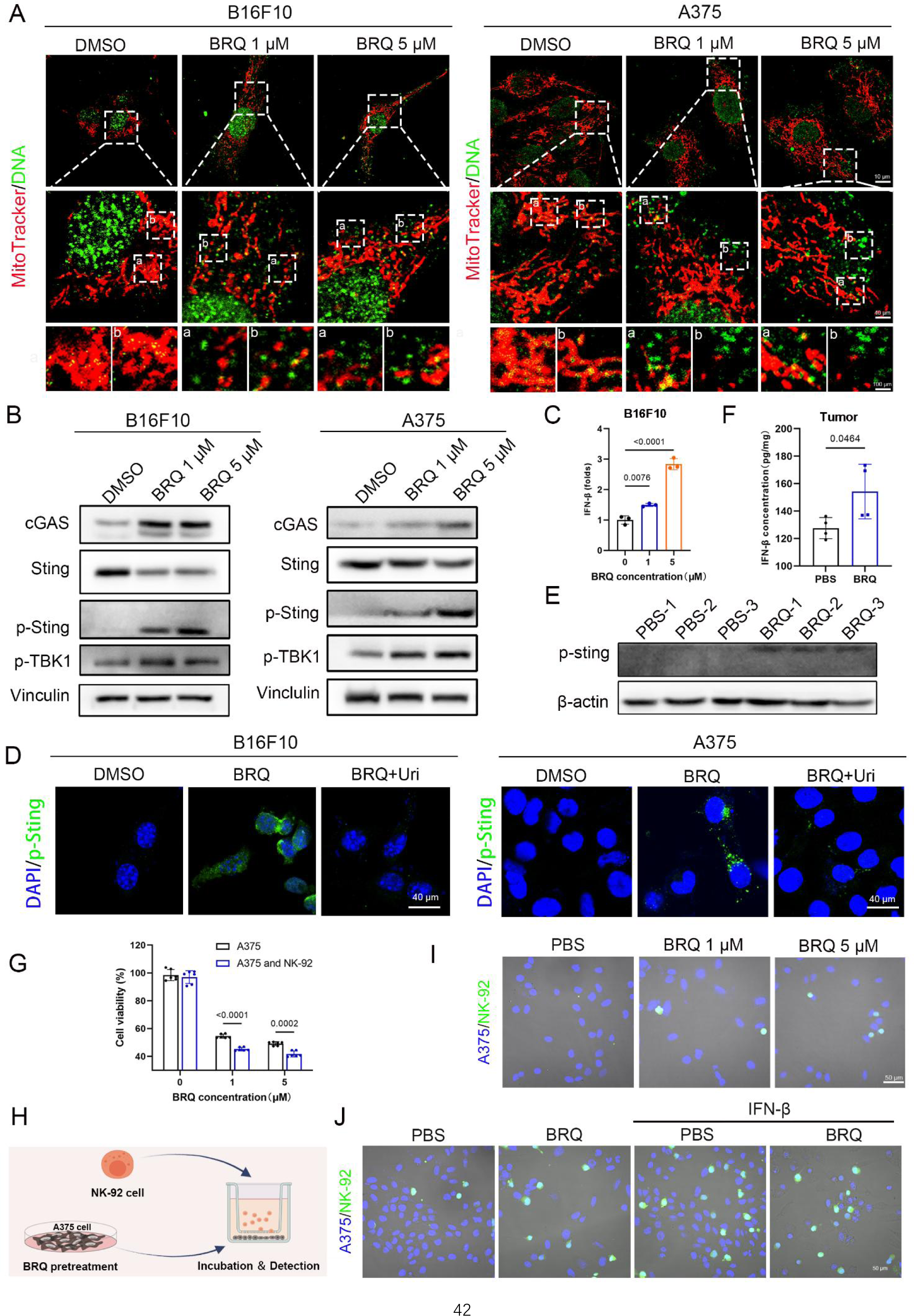
BRQ activates cGAS-STING pathway to enhance the antitumor immunity of NK cells. (A) CLSM images of mtDNA released from mitochondrial in B16F10 and A375 (Red: mitochondrial; green: DNA). (B) Western blot analysis of protein involved in cGAS-STING pathway. (C) ELISA analysis of INF-β in culture medium of untreated, BRQ treated B16F10 cells. (D) p-STING expression in B16F10 and A375 cells with DMSO, BRQ (5 μM) or BRQ (5 μM) plus uridine (100 μM). (E) Western blot analysis of p-STING in tumor tissues (n = 3 for each group). (F) ELISA analysis of INF-β in tumors of untreated or BRQ treated mice. (G) Cell viability of A375 cells cocultured or not with NK-92 cells after pretreated with BRQ. The A375 alone or co-cultured with NK-92 cells without drug treatment were used as control, respectively. (H) *In vitro* experiments for NK cells infiltration after treatment: A375 cells were pretreated with BRQ for 24 h and then co-cultured with NK-92 cells in a transwell system. (I) Images of NK-92 contact with pretreated A375 cells as shown in (H) (green: NK-92 label with CFSE, blue: A375 label with DAPI). (J) Images of NK-92 contact with A375 cells. A375 cells were pretreated with BRQ and then treated with or without IFN-β as shown in (H) (green: NK-92 label with CFSE, blue: A375 label with DAPI).

Uridine serves as the precursor of pyrimidine nucleotides, and its exogenous supplementation has been used to counterbalance the depletion of endogenous uridine resulting from DHODH inhibition. To investigate whether the activation of the cGAS-STING pathway arises from a deficiency in pyrimidine synthesis, we concurrently treated cells with uridine and BRQ. The results indicated that exogenous uridine supplementation attenuated the activation of p-STING in both B16F10 and A375 cells (Fig. 5D). These findings strongly indicate that BRQ effectively activates the cGAS-STING pathway *in vitro*, and this activation can be mitigated by exogenous uridine supplementation.

To confirm that BRQ activates the cGAS-STING signaling pathway *in vivo*, we analyzed tumor tissues from three mice per group to assess p-STING protein expression. BRQ treatment led to the up-regulation of p-STING expression (Fig. 5E), and this finding was further validated through immunohistochemistry (Fig. S4B). Futhermore, both ELISA and western blot results indicated that IFN-β levels were upregulated following BRQ treatment in tumor tissues (Fig. 5F and Fig. S4C). These results indicate that BRQ activates the cGAS-STING signaling pathway and induces the expression of IFN-Ⅰ, which are important for NK cells induced antitumor immunity.

To investigate whether BRQ enhances NK cells cytotoxicity against cancer cells *in vitro*, A375 cells were treated with BRQ at indicated concentrations, followed by coculturing with NK-92 cells. Tumor cells viability assays demonstrated that BRQ-treated A375 cells became more susceptible to NK cell-mediated cytotoxicity (Fig. 5G). Additionally, we implemented a transwell co-culture system with NK-92 cells in the upper chamber and A375 cells in the bottom chamber to investigate whether BRQ facilitate NK cells infiltration (Fig. 5H). Microscopic examination showed that BRQ-pretreated A375 cells were more readily recognized by NK-92 cells (Fig. 5I). Considering the pivotal role of IFN-β in regulating of NK cells numbers, activation, and antitumor activity [27], we supplemented exogenous IFN-β into the bottom chamber to investigate NK cells infiltration. Representative images showed that IFN-β facilitated NK cells infiltration, similar to the effect observed with BRQ treatment, which indicated that BRQ-induced IFN-β release is critical for NK cells infiltration (Fig. 5J).

In summary, these compelling results clearly demonstrate that BRQ exerts potent activation of the STING signaling pathway both *in vitro* and *in vivo*, consequently mediating an effective antitumor immune response which rely on NK cells.

### BRQ induces pyroptosis in melanoma cells and synergizes with NK cells

Notably, the formation of pores on the plasma membrane was observed following stimulation with BRQ in both B16F10 and A375 (Fig. 6A-B), consistent with the hallmark features of pyroptosis. Pyroptosis, a significant innate immune response [28], is closely related to NK cells. Granzyme A from NK cells and cytotoxic T lymphocytes [29], as well as killer-cells granzyme B, have been reported to initiate pyroptosis in target cells by cleaving GSDMB and GSDME, respectively. Importantly, the absence of GSDME in tumors results in reduced NK cells presence in the tumor microenvironment [30]. Thus, we explored the precise mechanism by which BRQ induces pyroptosis and the role of NK cells in activating tumor cells pyroptosis.

**Fig. 6.**
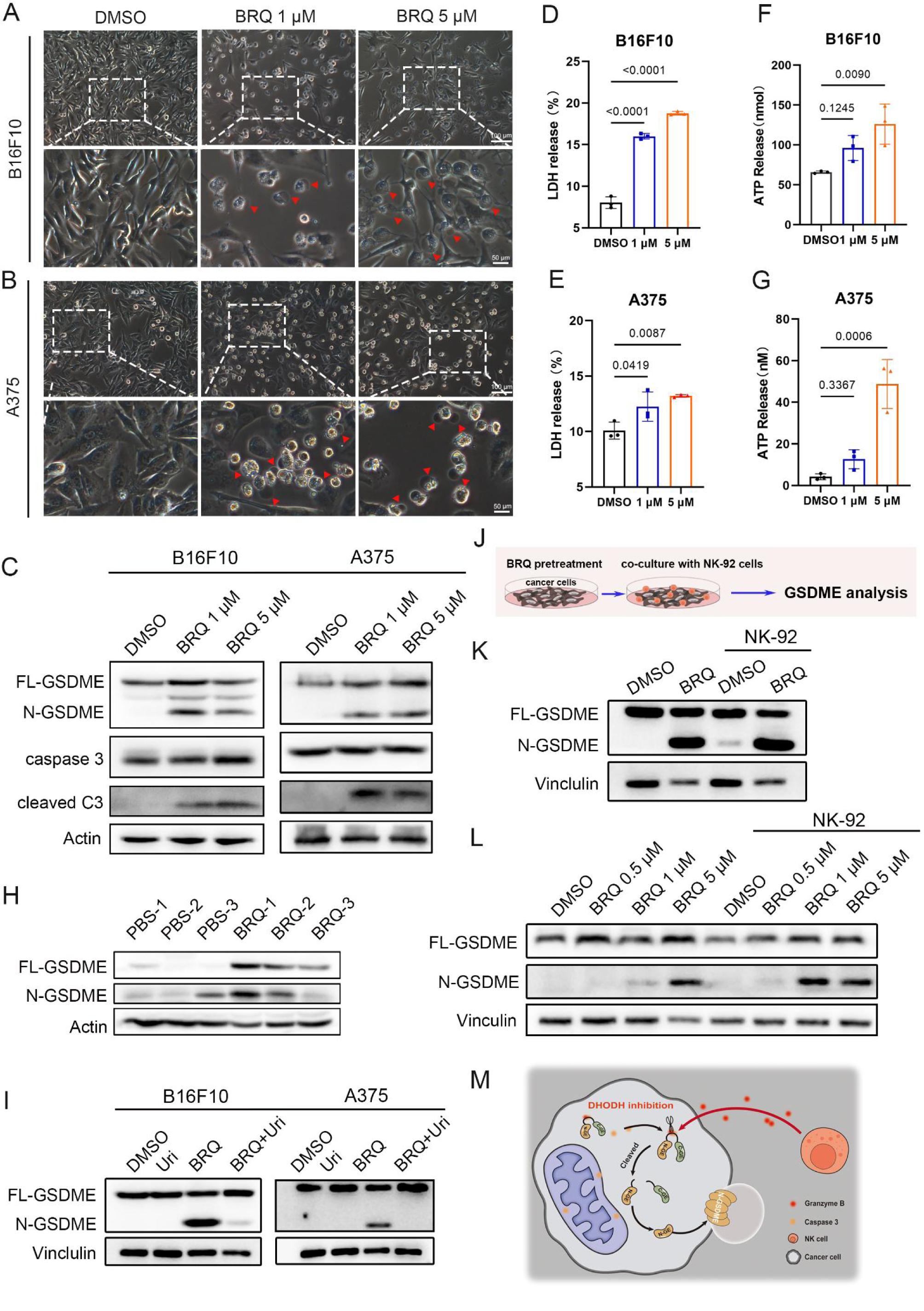
BRQ induces pyroptosis in melanoma cells and synergizes with NK cells. (A-B) Representative images of morphological alterations after BRQ treatment in B16F10 and A375 cells. Arrow indicated cell swelling and rupture. (C) Western blot analysis of GSDME and caspase 3 in B16F10 and A375 cells. (D-E) LDH released in culture supernatants after BRQ treatment in B16F10 and A375 cells. (F-G) ATP released in culture supernatants after BRQ treatment in B16F10 and A375 cells. (H) Western blot analysis of GSDME in tumor tissues (n = 3 for each group). (I) Western blot analysis of GSDME in untreated, uridine treated, BRQ treated or BRQ and uridine treated tumor cells. (J) *In vitro* experiments for NK cells enhancing tumor cells pyroptosis: A375 cells were treated with BRQ for 24 h and then co-cultured with NK-92 cells for 2 h to analysis the expression of GSDME in A375 cells. (K-L) Western blot analysis of GSDME in indicated groups as shown in (J). (M) Schematic representation of NK cells aggravation of cancer cells pyroptosis.

The DEGs were enriched in the NOD-like receptor signaling pathway (Fig. 4A). Among these, NLRP3 holds particular significance as a crucial member of the NOD-like receptor family. The activation of inflammasomes orchestrated by NLRP3 can lead to pyroptosis, a process reliant on GSDMD. To investigate whether BRQ induces pyroptosis through GSDMD, the protein expression of GSDMD was assessed. However, it was observed that BRQ did not induce the production of N-GSDMD in either A375 or B16F10 cell lines (Fig. S5A-B). Nevertheless, pyroptosis can also be elicited through caspase 3 cleavage of GSDME [31]. Interestingly, both cleaved caspase 3 and N-GSDME demonstrated a significant increase in both A375 and B16F10 cells following BRQ incubation, while the full-length GSDME (FL-GSDME) and caspase 3 remained unchanged (Fig. 6C). These results indicate that BRQ activates caspase 3 and triggers pyroptosis by cleaving GSDME.

Another characteristic of pyroptosis is the release of cellular contents. The results showed that lactate dehydrogenase (LDH) release increased in a dose-dependent manner after BRQ treatment, demonstrating plasma membrane rupture and leakage (Fig. 6D-E). Additionally, adenosine triphosphate (ATP), a type of damage-associated molecular pattern (DAMP), which can be released from pyroptotic cells, was significantly increased in the cells supernatant (Fig. 6F-G). Furthermore, flow cytometry analysis of propidium iodide (PI) and annexin Ⅴ staining revealed that BRQ markedly induced cell death (Fig. S5C-D). Thus, it is evident that BRQ triggers pyroptosis through the cleavage of GSDME *in vitro*.

To further validate this conclusion *in vivo*, we examined tumors from three mice per group to detect GSDME protein expression. Remarkably, N-GSDME were noticeably upregulated (Fig. 6H), which concurred with the findings from immunohistochemistry (Fig. S5E). Additionally, in response to BRQ treatment, the expression of high-mobility group box 1 (HMGB1) in tumors increased (Fig. S5F), which contribute to the activation of DC cells (Fig. S2A-B). Moreover, exogenous supplementation with uridine inhibited the production of N-GSDME in B16F10 and A375 cells, indicated that BRQ induced pyroptosis is related to pyrimidine metabolism (Fig. 6I). Therefore, BRQ effectively triggers melanoma cells pyroptosis.

To investigate whether tumor infiltrating NK cells enhanced BRQ induced pyroptosis, A375 cells were pretreated with BRQ and then cocultured with NK-92 cells. Subsequently, the A375 cells were collected to detect the N-GSDME protein expression (Fig. 6J). Significantly, NK-92 cells can induce A375 cells pyroptosis without BRQ treatment (Fig. 6K) and the expression of N-GSDME in A375 cells treated with BRQ were upregulated in a higher level after co-culture with NK-92 cells (Fig. 6K-L). These findings demonstrate that tumor cell pyroptosis promotes NK cell infiltration, and reciprocally, NK cells enhance tumor cell pyroptosis, playing a crucial role in BRQ-induced antitumor immunological effects (Fig. 6M).

### Mitochondrial oxidative stress as the origin of BRQ induced antitumor immunity

To explore the origin of BRQ induced antitumor immunity, we focus on the unique properties of DHODH. DHODH, situated on the external surface of the mitochondrial inner membrane, establishes a connection with the respiratory chain through the coenzyme Q pool [5]. The deficiency in DHODH activity impedes the proper functioning of the respiratory chain, consequently leading to the initiation of oxidative damage [32]. Given that mitochondria serve as pivotal center for immune responses [33], we postulate that BRQ-induced antitumor immunity originates from mitochondrial oxidative stress.

To assess mitochondrial impairment, we initially detected reactive oxygen species (ROS) production subsequent to BRQ stimulation. Notably, BRQ elicited a concentration and time-dependent escalation in ROS levels in both A375 and B16F10 cell lines (Fig. 7A and Fig. S6A). Furthermore, analysis of the JC-1 staining demonstrated an augmentation in the green fluorescence of JC-1 monomers following BRQ treatment, indicating a reduction in mitochondrial membrane potential (Fig. 7B and Fig. S6B).

**Fig. 7.**
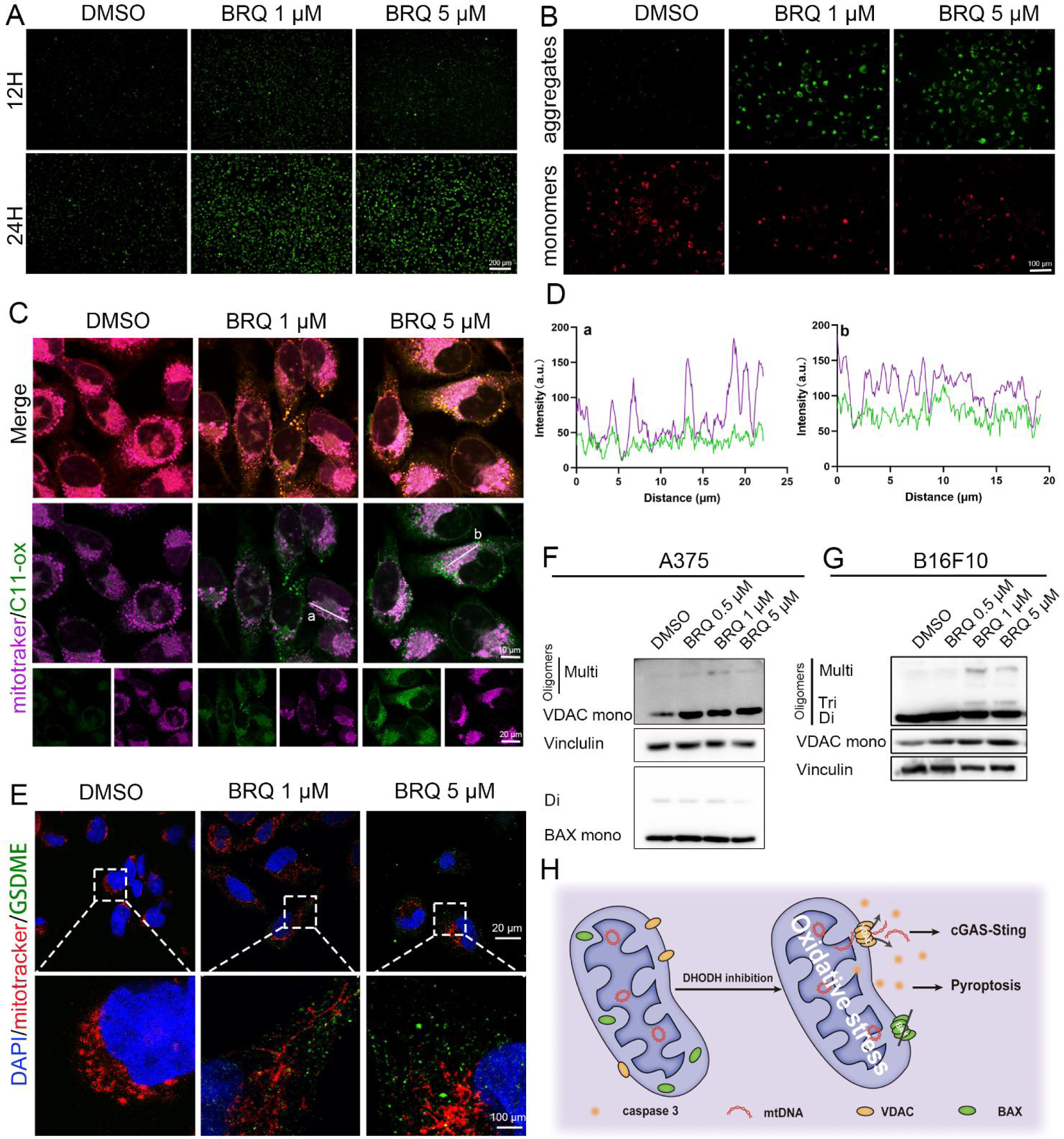
BRQ induces mitochondrial oxidative stress and mtDNA released via VDAC. (A) ROS was detected by DCFH-DA in A375 cells. (B) JC-1 analysis in A375 cells. (C) Representative images of A375 co-stained BODIPY C11(Oxidized BODIPY-green/non oxidized BODIPY-red) and mitochondrial (purple). (D) Colocalization analysis of mitochondrial and oxidized BODIPY in (C). (E) Representative images of A375 co-stained mitochondrial(red) and GSDME (green). DAPI was used to stain nuclei. (F) Western blot analysis of VDAC and BAX in A375 cells. (G) Western blot analysis of VDAC in A375 cells. (H) Schematic showing the release of mtDNA from mitochondria.

Considering the susceptibility of mitochondrial membranes to oxidative injury and the protective role of DHODH against lipid peroxidation, particularly in GPX4 low expression cells [34], we subsequently measured mitochondrial lipid peroxidation. In comparison to the 786-O cell line (GPX4 high expression cells) [34], A375 and B16F10 cells exhibited much lower GPX4 expression, rendering them more sensitive to BRQ mediated DHODH inhibition (Fig. S6C). Confocal laser scanning microscopy (CLSM) results provided compelling evidence that BRQ prominently augmented mitochondrial lipid peroxidation (Fig. 7C-D and Fig. S6D-E). Collectively, the comprehensive findings underscore the pronounced induction of oxidative stress within the mitochondrial by BRQ treatment.

Given the release of mtDNA as observed in Fig. 5A, we delved into the underlying mechanism governing mtDNA release from damaged mitochondria. Mitochondrial outer membrane permeabilization (MOMP) emerged as a key contributor to mtDNA release. Following MOMP, the gradual expansion of outer membrane pores triggers extrusion and rupture of the inner mitochondrial membrane (IMM) [35]. Although mitochondrial membrane permeabilization can be facilitated by GSDME-N [36], it is noteworthy that despite BRQ significantly elevating GSDME expression, no discernible co-localization between mitochondria and GSDME was detected via CLSM (Fig. 7E). These observations negate the involvement of GSDME-N in mtDNA release.

Moreover, BAX/BAK oligomerization, known to form pores in the IMM that facilitates mtDNA release [37], was not observed in A375 cells after BRQ treatment (Fig. 7F). Voltage-dependent anion channel (VDAC) oligomers have been recognized as inducers of mtDNA release [38]. Indeed, VDAC oligomers were detected in A375 cells (Fig. 7F), which was verified in B16F10 (Fig. 7G). In light of these findings, it is reasonable to deduce that BRQ induced mtDNA release is orchestrated through VDAC pores. Collectively, our investigations establish VDAC pores triggers BRQ induced mtDNA release (Fig. 7H).

In summary, the release of mtDNA from compromised mitochondria instigates the activation of the cGAS-STING signaling pathway. Concurrently, mitochondrial oxidative stress plays a pivotal role in expediting pyroptosis [39]. Cumulatively, these findings underscore the critical role of damaged mitochondria as the underlying source driving BRQ-induced antitumor immunity.

### EA6, a more effective DHODH inhibitor

In the quest to enhance the antitumor efficacy of BRQ, a series of derivatives based on BRQ were designed and synthesized (Fig. 8A). Initially, by introducing electron-withdrawing groups such as F and Cl to the 6th position of the quinoline ring, compounds AA3, EA3 were obtained, and by introducing electron-donating groups such as -CH3 and -OCH3 to the quinoline ring, compounds BA3, BA4, CA2, CA4, and CA8 were obtained. Activity screening revealed that, compared to BRQ, replacing electron-withdrawing groups on the 6th position of the quinoline ring with electron-donating groups resulted in better antitumor activity.

**Fig. 8.**
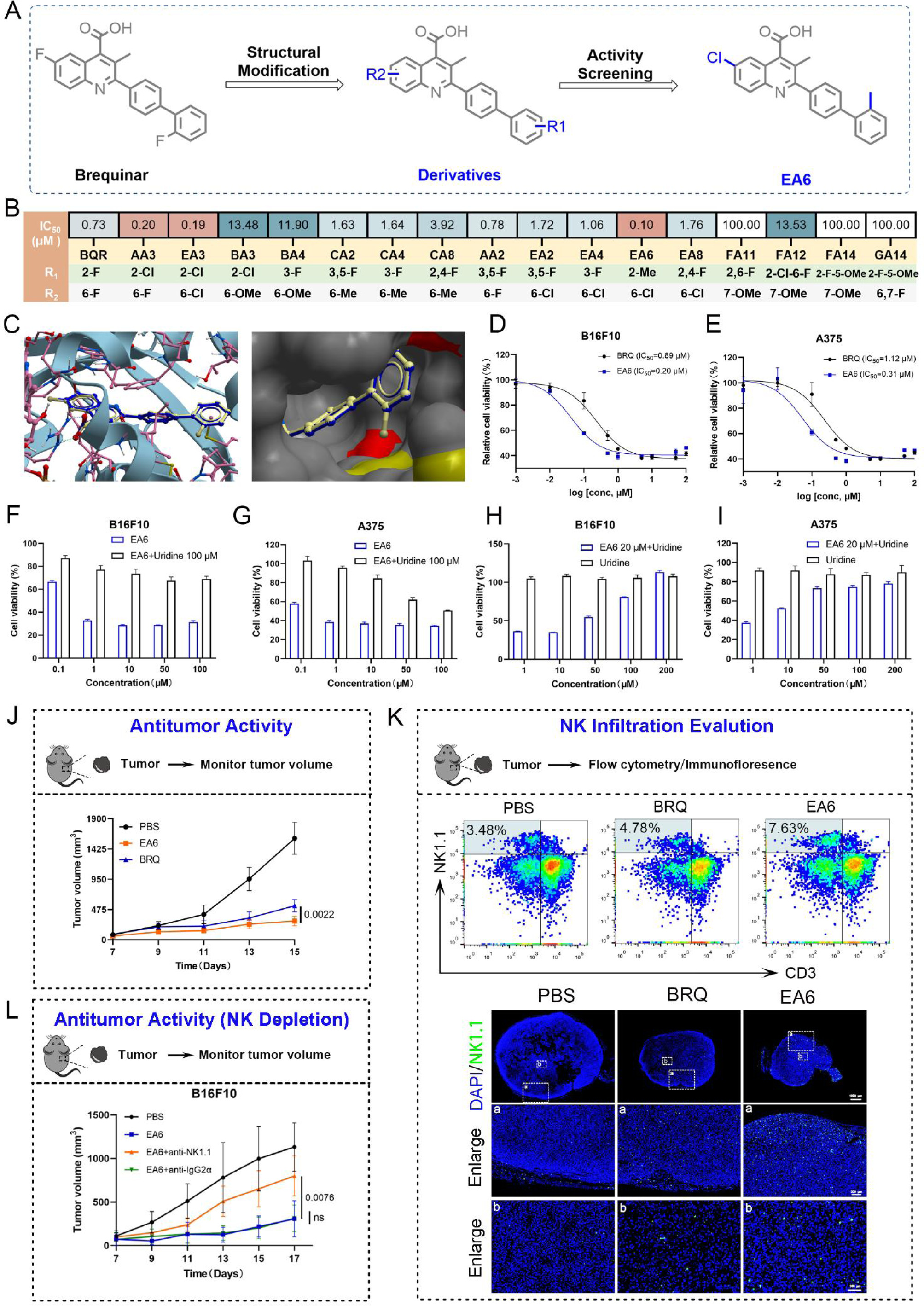
EA6, a more effective DHODH inhibitor. (A) Structural transformation from BRQ to EA6. (B) IC50 values of BRQ and BRQ derivatives drugs to B16F10 cell lines at 72 h. (C) Comparison of the binding sites of BRQ (blue) and EA6 (gray) with DHODH (PDB ID:1D3G). (D-E) Cell viability in B16F10 or A375 treated with BRQ or EA6. (F-G) Cell viability in B16F10 or A375 treated with EA6 or EA6+uridine. (H-I) Cell viability in B16F10 or A375 treated with uridine or uridine+EA6. (J) B16F10 tumors growth in mice treated with vehicle, BRQ (30 mg/kg, i.p.) or EA6 (30 mg/kg, i.p.) once ever two days. (K) Representative flow cytometry images and immunofluorescence images of NK cells in tumors from PBS, BRQ (30 mg/kg, i.p.) or EA6 (30 mg/kg, i.p.) treated mice. Mice were treated every two days, and after three times the tumors was collected for testing. (L) B16F10 tumor growth in mice treated with PBS, EA6 (30 mg/kg, i.p.), the combination of EA6 (30 mg/kg, i.p.) and anti-IgG2α (300 μg, i.p.) or the combination of EA6 (30 mg/kg, i.p.) and anti-anti-NK1.1 (300 ug, i.p.).

Subsequently, on the basis of introducing F and Cl at the 6th position of the quinoline ring, we further explored the effects of substituents at different positions on the biphenyl ring on its activity, then designed and synthesized compounds AA2, EA2, EA4, EA6, and EA8. The results showed that the compound with -CH3 substitution on the biphenyl ring exhibited significantly better antitumor activity than the F and Cl substitutions.

In addition, to further investigate whether antitumor activity could be effectively enhanced by introducing -OCH3 at the 7th position of the quinoline ring and introducing F at both the 6th and 7th positions simultaneously, compounds FA11, FA12, FA14, and GA14 were synthesized. Unfortunately, these structural modifications still exhibited poor antitumor activity. Ultimately, EA6 was selected as the compound with the highest activity (Fig. 8B and Table 1).

To further investigate the reasons for the increased activity of EA6, molecular docking was performed between EA6 and BRQ with DHODH protein (PDB ID: 1D3G). The results indicated that the methyl substitution on the biphenyl ring of EA6 increased its binding to the hydrophobic pocket relative to BRQ, thereby providing a theoretical basis for its enhanced activity (Fig. 8C). Subsequent cytotoxicity tests in A375 and B16F10 cell lines revealed that EA6 exhibited approximately four times higher activity compared to BRQ (Fig. 8D-E). The results of cAM/PI staining further confirmed that EA6 significantly induced cells death at a concentration of 500 nM (Fig. S7A).

To substantiate that EA6 induced cells death relied on the inhibition of DHODH, uridine was employed to mitigate cellular damage. EA6 displayed a dose-dependent inhibition of melanoma cells proliferation, which could be fully reversed upon supplementation with 100 μM uridine (Fig. 8F-G). Simultaneously, 20 μM EA6 notably decreased cells viability, and this effect was progressively reversed in a dose-dependent manner by uridine (Fig. 8H-I). Consequently, EA6 emerged as a potential *in vitro* inhibitor of DHODH.

Subsequently, the *in vivo* antitumor effect of EA6 was assessed in C57BL/6 mice bearing B16F10 tumors. Both BRQ and EA6 robustly inhibited tumor growth, with EA6 exhibiting superior effectiveness compared to BRQ (Fig. 8J). Additionally, the expression levels of GSDME and p-STING were upregulated by EA6 (Fig. S7B). And post-treatment examination of major organs revealed no severe structural or pathological changes (Fig. S7C). Taken together, EA6 induces melanoma cells pyroptosis and activates the cGAS-STING signaling pathway *in vivo*, which demonstrated superior antitumor activity.

To explore whether EA6 has a promising potential in NK cells based cancer immunotherapy, flow cytometry and immunofluorescence were used to investage NK cells inflitration. Flow cytometry analysis showed that EA6 treatment increased the proportion of NK cells from 3.48% to 7.63% of CD45^+^ lymphocytes, indicating a greater potential to induce NK cells infiltration compared to BRQ (Fig. 8K and Fig. S7D). Tumor-infiltrating NK cells were also assessed through immunofluorescence staining, as depicted in Fig. 8K. In the PBS group, NK cells were mainly aggregated at the tumors periphery and were less abundant. Conversely, following treatment with BRQ/EA6, a significant infiltration of NK cells into the tumor core was evident. Remarkably, EA6 exhibited a higher capacity to promote NK cells infiltration within the tumor microenvironment compared to the BRQ treated group. To verify that the antitumor activity of EA6 is dependent on NK cells, anti-NK1.1 antibody was used to deplete NK cells in C57BL/6 mice, and the antitumor activity of EA6 was explored in NK-depleted mice. NK depletion diminished the antitumor efficacy of EA6, emphasizing that its therapeutic efficacy is dependent on NK cells (Fig. 8L and Fig. S7E-F).

In summary, EA6 emerged as a more potent DHODH inhibitor, demonstrating strong tumor regression in melanoma and acting as an NK cells-dependent therapeutic agent.

## DISCUSSION

In recent years, our research group has primarily focused on tumor metabolism and cancer immunotherapy [40, 41]. Restraining nucleotide metabolism is not only a therapeutic target for chemotherapy but also for improving the therapeutic efficacy of cancer immunotherapy. Disrupting nucleotide metabolism increases the genomic instability via destructing of purine and pyrimidine pools which finally promoting immunogenicity [3]. DHODH, a key rate-limiting enzyme in *de novo* pyrimidine synthesis, plays an important part in tumor immunity [10, 12]. However, the effect of DHODH inhibition on NK cells remains unknown, and malignant melanomas escape immunosurveillance via the loss/down-regulation of MHC-I expression which are susceptible to NK cells induced toxicity. Thus, it is worthwhile to explore that weather NK cells paly a potential part in antitumor immunity responded to DHODH inhibition in melanoma.

In our study, we investigated the differential response of immune effector cells, specifically CD8^+^ T cells and NK cells, to DHODH inhibition. Our findings revealed that while CD8^+^ T cells exhibited no significant increase in tumor tissues following BRQ treatment, NK cells infiltration saw a remarkable surge. The depletion of NK cells in C57BL/6 mice led to a notable reduction in the antitumor effect of DHODH inhibition, underscoring the indispensable role of NK cells in DHODH induced antitumor responses.

The pivotal role of mitochondria in cellular energy production is well-documented, but their emerging importance as centers of immune responses has opened new avenues of research [22]. DHODH, situated on the outer surface of the mitochondrial inner membrane and interconnected with the respiratory chain, provides a compelling connection between mitochondrial function and immune activation. It has been revealed that high concentrations of BRQ induced mitochondria-associated ferroptosis in cancer cells [34], further substantiating the link between DHODH and mitochondrial induced immune responses.

Our investigation focused on the hypothesis that DHODH inhibition induced mitochondrial damage serves as the primary source of immune activation. In this regard, DHODH inhibition triggered mitochondrial oxidative stress, culminating in VDAC oligomerization. VDAC oligomers formed pores in the mitochondrial outer membrane, facilitating the release of mtDNA and proteins. Free mtDNA in the cytoplasm was subsequently recognized by cGAS, initiating the STING pathway, a crucial driver of the antitumor immune response. Simultaneously, caspase 3, released from damaged mitochondria, cleaved GSDME, triggering pyroptosis—a form of immunogenic cell death that releases DAMPs and fuels the immune response. In concert, mitochondrial oxidative stress induced by DHODH inhibition thus stimulated immune activation via pyroptosis and the cGAS-STING pathway.

The production of IFN-Ⅰ stands as a hallmark of STING pathway activation, with IFN-Ⅰ playing a crucial role in NK cells homeostasis, activation, and antitumor functionality [27]. We found that BRQ promoted the antitumor activity of NK cells by stimulating of IFN-β release from tumor cells.

And our *in vivo* findings indicated a marked up-regulation of both IFN-β in the tumor microenvironment. These observations further elucidate the role of DHODH inhibition in facilitating NK cells infiltration through the up-regulation of IFN-Ⅰ.

NK cells release Granzyme B, which can cleave GSDME at the caspase 3 cleavage site [30], a process that was confirmed in our study. Intriguingly, N-GSDME was upregulated in cancer cells following coculture with NK-92 cells, indicating that the interaction between NK cells and tumor cells enhances pyroptosis, further contributing to the antitumor immune response.

Lastly, our research resulted in the design of EA6, an enhanced DHODH inhibitor. EA6 demonstrated significantly higher activity compared to BRQ, inducing NK cells infiltration and tumor growth inhibition.

In conclusion, our study underscores the capacity of DHODH inhibition to trigger pyroptosis and the cGAS-STING signaling pathways, leading to NK cells induced antitumor immunity. This finding opens new avenues for enhancing the efficacy of targeted nucleotide metabolism in cancer therapy. This innovative approach also holds promise for the synergistic combination of chemotherapy and immunotherapy.

While our findings shed light on the promising synergy between DHODH inhibition and NK cells mediated antitumor immunity, several research gaps and future directions should be addressed. First, a deeper understanding of the specific regulatory mechanisms governing NK cells activation and their interactions with other immune cells remains essential. Moreover, a critical future avenue should encompass the development of novel therapeutic strategies building upon our findings. Exploring combinatory approaches, such as combining DHODH inhibitors with immunotherapies, may hold the key to unlocking enhanced anticancer efficacy while minimizing side effects. Such an approach could lead to more targeted and personalized cancer treatments, capitalizing on the synergistic effects of nucleotide metabolism disruption and immune stimulation.

## MATERIALS AND METHODS

### Cell culture

The B16F10 (murine melanoma cells), A375 (human melanoma cells), 786-O (human renal carcinoma cells), PUMC-HUVEC-T1 (human endothelial cells) and NK-92 (human natural killer cells) were procured from Procell Life Science & Technology (Wuhan, China). B16F10, 786-O, PUMC-HUVEC-T1 and A375 cells were cultured in DMEM/RPMI-1640 medium supplemented with 10% fetal bovine serum (Excell Bio, Shanghai, FSP500) and 1% penicillin/streptomycin (MacGene, Beijing, CC004). NK-92 cells were cultured with specific medium (CM-0530) procured from Procell Life Science & Technology. Incubation of all cell lines was carried out at 37 °C under a 5% CO2 atmosphere.

### Mice and antitumor therapy

Female C57BL/6 mice aged 6-8 weeks were obtained from SPF (Beijing) Biotechnology Co., Ltd. B16F10 cells were harvested and suspended in serum-free RPMI-1640 medium. Subsequently, 5×10^6^ cells were subcutaneously injected into each mouse. When the tumor size reached approximately 50 mm^3^, the mice were treated with PBS (intraperitoneal injection, once every two days), BRQ (intraperitoneal injection, 30 mg/kg, once every two days), or EA6 (intraperitoneal injection, 30 mg/kg, once every two days).

For NK cells depletion, 250 μg of anti-NK1.1 or IgG2α antibodies were diluted in dilution buffer (InVivoPure pH 7.0 Dilution Buffer) and intraperitoneally injected into the mice on days 6, 8, 10, and 12. Spleens were collected 24 hours after the first injection of anti-NK1.1 antibody for flow cytometry analysis to verify the depletion of NK cells.

The tumor volumes and body weights of the mice were measured every 2 days. The tumor volume (V) was calculated using the following equation, where ’a’ represents the longest dimension of the tumor and ’b’ represents the shortest dimension of the tumor: V = a*b^2^/2

All procedures conducted on animals complied with the ARRIVE guidelines (EU Directive 2010/63/ EU for animal experiments). Ethics approval was granted by the Ethics Committee of Northwestern Polytechnical University. Animal experiments were carried out according to the Animal Experiment Center of Northwestern Polytechnical University Animal Care and Use Guidelines (NO. 202201173).

### Western blot

After the indicated treatment duration, cells were collected and lysed for subsequent immunoblotting analysis. The protein concentration was determined using the BCA Protein Assay Kit (P0010, Beyotime). Approximately 20 μg of protein from each group cells lysate was loaded onto an SDS-PAGE gel and electrophoresed at 120 V. The proteins were then transferred to PVDF membranes at 250 mA for the specified period of time. Following transfer, the membranes were blocked with 5% non-fat dry milk in PBST for 1 h at room temperature. Subsequently, the membranes were incubated overnight with the primary antibody, followed by three washes with PBST. After that, the membranes were incubated with the secondary antibody for 1 h at room temperature and washed three times with PBST. Finally, the membranes were immersed in an ECL substrate, and the images were captured using a Chemiluminescence Imaging System. Detailed information of antibodies as shown in **Table S2** in supporting information.

### Enzyme-linked immunosorbent assay

After a 36 h treatment with BRQ, the culture medium was collected from the cells and centrifuged for subsequent analysis. In the animal study, tumors were collected at the indicated time points and lysed with ELISA extraction buffer on ice for 2 h. The supernatant was then centrifuged for further analysis. The concentrations of IL-10, IL-12, IL-1B, TGF-β, and IFN-γ were determined using an ELISA kit (Fanke, Shanghai) according to the manufacturer’s instructions.

### Cytotoxicity

Cytotoxicity was assessed using the MTT assay. Cells in the logarithmic growth phase were harvested and seeded in a 96-well plate at a density of approximately 5000 cells/well. The respective drugs were diluted in the indicated concentrations in the culture medium. The cells were then treated with the prepared drug solutions for 72 h. Afterward, the cells were incubated with serum-free medium containing 0.5 mg/ml MTT for 4 h in a carbon dioxide cell culture box. The formazan crystals were dissolved in DMSO, and the optical absorbance values at 450 nm were measured using a microplate multifunctional reader (Tecan--Tecan Spark, Switzerland).

### NK cell-induced cytotoxicity to BRQ-treated A375 cells

A375 cells were pretreated with BRQ for 30 h and then replacement of fresh medium to continuation incubation for 6 h. Subsequently, NK-92 cells were added to A375 cells in a ratio of 1:5 and cocultured for 12 h. Finally, cell viability of A375 were evaluated by MTT assay.

To characterize NK-92 cells recognition of A375 cells, briefly, A375 cells were labeled with Hochest 33342 and NK92 cells were labeled with CSFE. After co-culture with the transwell system, the contact between NK-92 cells and A375 cells were evaluated by CLSM.

### Immunofluorescence

Cells were grown on coverslips, and the treated cells were washed twice with PBS. Subsequently, the cells were fixed with 4% paraformaldehyde for 10 minutes. For intracellular proteins such as p-STING, which required permeabilization, the cells were treated with 0.2% Triton X-100 for 10 minutes. After washing with cold PBS three times, the cells were blocked with 3% BSA in PBST (containing 22.5 mg/ml glycine). The cells were then incubated with the indicated primary antibody overnight at 4 °C. After three washes, corresponding secondary antibodies conjugated with fluorescence dyes were incubated with the cells at room temperature for 1 h. Following another round of washing, the cells were stained with DAPI for 10 minutes. CLSM was used to collect the images.

### Mitochondrial function analysis

The mitochondrial membrane potential was evaluated using the Enhanced Mitochondrial Membrane Potential Assay Kit with JC-1 (C2003S, Beyotime). The status of the mitochondrial permeability transition pore (mPTP) was assessed using the Mitochondrial Permeability Transition Pore Assay Kit (C2009S, Beyotime). A375 and B16F10 cells were treated with the indicated concentration of BRQ for 24 h and then stained with JC-1 or Calcein AM following the manufacturer’s instructions. The cells fluorescence were detected using a fluorescence microscope (OLYMPUS-OLYMPUS IX73) or a CLSM.

### Clinical date analysis

RNA-sequencing expression data and detailed clinical information for Skin Cutaneous Melanoma (SKCM) were obtained from The Cancer Genome Atlas (TCGA, https://portal.gdc.com). The immune score was evaluated using the immuneeconv package. Immuneeconv is an R software package that incorporates six algorithms, namely TIMER, EPIC, quanTIseq, MCP-counter, CIBERSORT, and xCell. In this study, we only presented the results obtained using the EPIC algorithm. All data analysis methods and R packages were implemented using R version. The significance of the results were determined using the student’s t-test.

### LDH and ATP release assay

A375 and B16F10 cells were treated with the specified concentrations of BRQ for 48 h. After centrifugation at 3000 rpm for 5 minutes, the supernatant was collected for the detection of LDH release and ATP levels. The LDH Release Assay Kit and Enhanced ATP Assay Kit were used, following the provided instructions, to perform these measurements.

### Flow cytometry analysis

After three rounds of treatment, the mice were euthanized to examine their immune phenotype. The analysis focused on mature DC cells (CD11c^+^CD80^+^CD86^+^), T cells (CD45^+^CD3^+^CD4^+^/CD8^+^), and NK cells (CD45^+^CD3^-^NK1.1^+^). Tumors, spleens, and lymph nodes were dissected, followed by digestion or homogenization to obtain single cells. These cells were then stained with specific antibodies and analyzed using flow cytometry. FlowJo_V10 software was utilized for data analysis and interpretation.

### RNA-seq

After treating B16F10 cells with DMSO or BRQ for 24 h, total RNA was extracted using Trizol (Thermofisher, 15596018) for further analysis. The extracted RNA samples were then processed for library preparation and sequenced on the illumina Novaseq™ 6000 platform by LC Bio Technology CO., Ltd (Hangzhou, China). Bioinformatic analysis of the sequencing data was conducted using the OmicStudio tools available at https://www.omicstudio.cn/tool. In the bioinformatic analysis, differentially expressed genes were identified based on specific criteria: genes with an absolute log2 fold change (|log2FC|) greater than or equal to 1 and a q-value (adjusted *P* value) less than 0.05 were considered as differentially expressed. These differentially expressed genes represent the subset of genes that showed significant changes in expression levels between the DMSO and BRQ treated groups.

### Tissue microarrays chips assay

The tissue microarray chips (TMA) used in this study were obtained from ZhuoLi Biotech Co., Ltd. (Shanghai, China). The TMA consisted of 5 normal skin tissues and 44 tumor tissues from patients with melanoma. Each case was represented twice on the TMA to ensure reproducibility and reliability of the results. Detailed information about the patients, including clinical and pathological characteristics, was provided by the company along with the TMA. This comprehensive dataset allowed for the correlation of molecular findings with clinicopathological features of the melanoma samples.

### Statistical analysis

The data analysis was performed using GraphPad Prism 9.5 software. The results are presented as mean ± SEM (standard error of the mean) or mean ± SD (standard deviation), depending on the specific analysis. Each experiment was repeated independently at least three times to ensure the reliability of the findings. Statistical significance was determined using the student’s t-test for comparisons between two groups, and One-way ANOVA followed by post hoc tests for comparisons among multiple groups. *P* value of less than 0.05 was considered statistically significant.

## DATA AVAILABILITY

The authors declare that all relevant data supporting the findings of this study are available within the paper, the Supplementary Information, and Source data. The RNA-sequencing data generated by this study have been deposited to GEO database under accession number: GSE245151.

## Supporting information

Supporting Information: Part 1

Supporting Information: Part 2

## ACKNOWLEDGEMENTS

This work was supported by the National Natural Science Foundation of China (82173682, 22367006, 82260545), Shenzhen Science and Technology Program (JCYJ20240813150807010), the Young Elite Scientist Sponsorship Program by CAST (YESS20200108), Shaanxi Provincial Health Research Project (2022A016), Xi’an Science and Technology Programme Project (23YXYJ0109).

## REFERENCES

1. Ali ES, Ben-Sahra I. Regulation of nucleotide metabolism in cancers and immune disorders. Trends Cell Biol. 2023; 3311: 950–966.

2. Mullen NJ, Singh PK. Nucleotide metabolism: a pan-cancer metabolic dependency. Nat Rev Cancer. 2023; 235: 275–294.

3. Wu HL, Gong Y, Ji P, Xie YF, Jiang YZ, Liu GY. Targeting nucleotide metabolism: a promising approach to enhance cancer immunotherapy. J Hematol Oncol. 2022; 151: 45.

4. Siddiqui A, Ceppi P. A non-proliferative role of pyrimidine metabolism in cancer. Mol Metab. 2020; 35: 100962.

5. Zhou Y, Tao L, Zhou X, Zuo Z, Gong J, Liu X, et al. DHODH and cancer: promising prospects to be explored. Cancer Metab. 2021; 91: 22.

6. Ladds M, van Leeuwen IMM, Drummond CJ, Chu S, Healy AR, Popova G, et al. A DHODH inhibitor increases p53 synthesis and enhances tumor cell killing by p53 degradation blockage. Nat Commun. 2018; 91: 1107.

7. Pal S, Kaplan JP, Nguyen H, Stopka SA, Savani MR, Regan MS, et al. A druggable addiction to de novo pyrimidine biosynthesis in diffuse midline glioma. Cancer Cell. 2022; 409: 957–972 e910.

8. Li L, Ng SR, Colón CI, Drapkin BJ, Hsu PP, Li Z, et al. Identification of DHODH as a therapeutic target in small cell lung cancer. Sci Transl Med. 2019; 11517.

9. Sykes DB, Kfoury YS, Mercier FE, Wawer MJ, Law JM, Haynes MK, et al. Inhibition of Dihydroorotate Dehydrogenase Overcomes Differentiation Blockade in Acute Myeloid Leukemia. Cell. 2016; 1671: 171–186 e115.

10. Sprenger HG, MacVicar T, Bahat A, Fiedler KU, Hermans S, Ehrentraut D, et al. Cellular pyrimidine imbalance triggers mitochondrial DNA-dependent innate immunity. Nat Metab. 2021; 35: 636–650.

11. Peeters MJW, Aehnlich P, Pizzella A, Molgaard K, Seremet T, Met O, et al. Mitochondrial-Linked De Novo Pyrimidine Biosynthesis Dictates Human T-Cell Proliferation but Not Expression of Effector Molecules. Front Immunol. 2021; 12: 718863.

12. Colligan SH, Amitrano AM, Zollo RA, Peresie J, Kramer ED, Morreale B, et al. Inhibiting the biogenesis of myeloid-derived suppressor cells enhances immunotherapy efficacy against mammary tumor progression. J Clin Invest. 2022; 13223: e158661.

13. Halbrook CJ, Pontious C, Kovalenko I, Lapienyte L, Dreyer S, Lee HJ, et al. Macrophage-Released Pyrimidines Inhibit Gemcitabine Therapy in Pancreatic Cancer. Cell Metab. 2019; 296: 1390–1399 e1396.

14. Maskalenko NA, Zhigarev D, Campbell KS. Harnessing natural killer cells for cancer immunotherapy: dispatching the first responders. Nat Rev Drug Discov. 2022; 218: 559–577.

15. Kyrysyuk O, Wucherpfennig KW. Designing Cancer Immunotherapies That Engage T Cells and NK Cells. Annu Rev Immunol. 2022; 41: 17–38.

16. White RM, Cech J, Ratanasirintrawoot S, Lin CY, Rahl PB, Burke CJ, et al. DHODH modulates transcriptional elongation in the neural crest and melanoma. Nature. 2011; 4717339: 518–522.

17. Wu YT, Fang Y, Wei Q, Shi H, Tan H, Deng Y, et al. Tumor-targeted delivery of a STING agonist improvescancer immunotherapy. Proc Natl Acad Sci USA. 2022; 11949: e2214278119.

18. Wang H, Wang X, Zhang X, Xu W. The promising role of tumor-associated macrophages in the treatment of cancer. Drug Resist Updat. 2024; 73: 101041.

19. Takashi N, Hiroko M, Mamoru H, Yusuke S, Yoshihiro H, Hideyoshi H. Liposomes loaded with a STING pathway ligand, cyclic di-GMP, enhance cancer immunotherapy against metastatic melanoma. Journal of Controlled Release. 2015; 216:149–57.

20. Batlle E, Massague J. Transforming Growth Factor-beta Signaling in Immunity and Cancer. Immunity. 2019; 504: 924–940.

21. Baumgartner CK, Ebrahimi-Nik H, Iracheta-Vellve A, Hamel KM, Olander KE, Davis TGR, et al. The PTPN2/PTPN1 inhibitor ABBV-CLS-484 unleashes potent anti-tumour immunity. Nature. 2023; 7984: 850–962.

22. West AP, Khoury-Hanold W, Staron M, Tal MC, Pineda CM, Lang SM, et al. Mitochondrial DNA stress primes the antiviral innate immune response. Nature. 2015; 5207548: 553–557.

23. Chin EN, Sulpizio A, Lairson LL. Targeting STING to promote antitumor immunity. Trends Cell Biol. 2023; 333: 189–203.

24. Feng B, Lu X, Zhang G, Zhao L, Mei D. STING agonist delivery by lipid calcium phosphate nanoparticles enhances immune activation for neuroblastoma. Acta Materia Medica. 2023; 22: 216–227.

25. Marcus A, Mao AJ, Lensink-Vasan M, Wang L, Vance RE, Raulet DH. Tumor-Derived cGAMP Triggers a STING-Mediated Interferon Response in Non-tumor Cells to Activate the NK Cell Response. Immunity. 2018; 4: 753–764.

26. Nicolai CJ, Wolf N, Chang IC, Kirn G, Marcus A, Ndubaku CO, et al. NK cells mediate clearance of CD8(+) T cell-resistant tumors in response to STING agonists. Sci Immunol. 2020; 545: aaz2738.

27. Swann JB, Hayakawa Y, Zerafa N, Sheehan KC, Scott B, Schreiber RD, et al. Type I IFN contributes to NK cell homeostasis, activation, and antitumor function. J Immunol. 2007; 17812: 7540–7549.

28. Wei X, Xie F, Zhou X, Wu Y, Yan H, Liu T, et al. Role of pyroptosis in inflammation and cancer. Cell Mol Immunol. 2022; 199: 971–992.

29. Zhou Z, He H, Wang K, Shi X, Wang Y, Su Y, et al. Granzyme A from cytotoxic lymphocytes cleaves GSDMB to trigger pyroptosis in target cells. Science. 2020; 3686494: eaaz7548.

30. Zhang Z, Zhang Y, Xia S, Kong Q, Li S, Liu X, et al. Gasdermin E suppresses tumour growth by activating anti-tumour immunity. Nature. 2020; 5797799: 415–420.

31. Wang Y, Gao W, Shi X, Ding J, Liu W, He H, et al. Chemotherapy drugs induce pyroptosis through caspase-3 cleavage of a gasdermin. Nature. 2017; 5477661: 99–103.

32. Boukalova S, Hubackova S, Milosevic M, Ezrova Z, Neuzil J, Rohlena J. Dihydroorotate dehydrogenase in oxidative phosphorylation and cancer. Biochim Biophys Acta Mol Basis Dis. 2020; 18666: 165759.

33. Mills EL, Kelly B, O’Neill LAJ. Mitochondria are the powerhouses of immunity. Nat Immunol. 2017; 185: 488–498.

34. Mao C, Liu X, Zhang Y, Lei G, Yan Y, Lee H, et al. DHODH-mediated ferroptosis defence is a targetable vulnerability in cancer. Nature. 2021; 5937860: 586–590.

35. Bock FJ, Tait SWG. Mitochondria as multifaceted regulators of cell death. Nat Rev Mol Cell Biol. 2020; 212: 85–100.

36. Rogers C, Erkes DA, Nardone A, Aplin AE, Fernandes-Alnemri T, Alnemri ES. Gasdermin pores permeabilize mitochondria to augment caspase-3 activation during apoptosis and inflammasome activation. Nat Commun. 2019; 101: 1689.

37. McArthur K, Whitehead LW, Heddleston JM, Li L, Padman BS, Oorschot V, et al. BAK/BAX macropores facilitate mitochondrial herniation and mtDNA efflux during apoptosis. Science. 2018; 3596378: eaao6047.

38. Kim J, Gupta R, Blanco LP, Yang S, Shteinfer-Kuzmine A, Wang K, et al. VDAC oligomers form mitochondrial pores to release mtDNA fragments and promote lupus-like disease. Science. 2019; 3666472: 1531–1536.

39. Kang R, Zeng L, Zhu S, Xie Y, Liu J, Wen Q, et al. Lipid Peroxidation Drives Gasdermin D-Mediated Pyroptosis in Lethal Polymicrobial Sepsis. Cell Host Microbe. 2018; 241: 97–108 e104.

40. Fan R, Deng A, Qi B, Zhang S, Sang R, Luo L, et al. CJ(2): A Novel Potent Platinum(IV) Prodrug Enhances Chemo-Immunotherapy by Facilitating PD-L1 Degradation in the Cytoplasm and Cytomembrane. J Med Chem. 2023; 661: 875–889.

41. Sang R, Fan R, Deng A, Gou J, Lin R, Zhao T, et al. Degradation of Hexokinase 2 Blocks Glycolysis and Induces GSDME-Dependent Pyroptosis to Amplify Immunogenic Cell Death for Breast Cancer Therapy. J Med Chem. 2023; 6613: 8464–8483.

