## Supporting Information: Part 1 for "Inhibition of DHODH Activates Pyroptosis and cGAS-STING Pathways to Enhance NK cell Infiltration Mediated Anti-tumor Immunity in Melanoma"

**Table S1：** $IC_{50}$ values of drugs to variable cell lines at 72 hours [a].

| Compounds | R1 | R2 | B16F10 |
|---|---|---|---|
| | | | $IC_{50}$ (μM) |
| BQR | - | - | 0.73±0.08 |
| AA2 | 3,5-F | 6-F | 0.78±0.32 |
| AA3 | 2-Cl | 6-F | 0.20±0.06 |
| BA3 | 2-Cl | 6-$OCH_3$ | 13.48±5.32 |
| BA4 | 3-F | 6-$OCH_3$ | 11.90±4.76 |
| CA2 | 3,5-F | 6-$CH_3$ | 1.63±0.41 |
| CA4 | 3-F | 6-$CH_3$ | 1.64±0.39 |
| CA8 | 2,4-F | 6-$CH_3$ | 3.92±0.56 |
| EA2 | 3,5-F | 6-Cl | 1.72±0.64 |
| EA3 | 2-Cl | 6-Cl | 0.19±0.07 |
| EA4 | 3-F | 6-Cl | 1.06±0.84 |
| EA6 | 2-$CH_3$ | 6-Cl | 0.10±0.05 |
| EA8 | 2,4-F | 6-Cl | 1.76±0.22 |
| FA11 | 2,6-F | 7-$OCH_3$ | ＞100 |
| FA12 | 2-Cl-6-F | 7-$OCH_3$ | 13.53±2.36 |
| FA14 | 2-F-5-$OCH_3$ | 7-$OCH_3$ | ＞100 |
| GA14 | 2-F-5-$OCH_3$ | 6,7-F | ＞100 |

[a] $IC_{50}$ values are represented by mean ± SD of three independent experiments.

**Table S2：** Detailed information of antibodies.

| Antibody | Supplier | Cat. Num. |
|---|---|---|
| DHODH Polyclonal antibody | Proteintech | 14877 |
| GSDME antibody | CST | 40618 |
| Caspase-3(D3R6Y) Rabbit mAb | CST | 14220 |
| Anti-Vinculin antibody | Abcam | ab129002 |
| STING(D1V5L) Rabbit mAb | CST | 50494 |
| P-STING(S365) Rabbit mAb | CST | 72971 |
| cGAS(D3080) Rabbit mAb | CST | 31659 |
| TBK1/NAK(D1B4) Rabbit mAb | CST | 3504 |
| p-TBK1/NAK(S172) Rabbit mAb | CST | 5483 |
| IRF-3(D83B9) Rabbit mAb | CST | 4302 |
| p-IRF-3(S396) Rabbit mAb | CST | 29047 |
| Anti-cGAS antibody | abcam | 302617 |
| P-STING(S365) Rabbit mAb | CST | 50907 |
| Anti-Glutathione peroxidase antibody | abcam | Ab125066 |
| Recombinant Anti-beta Actin antibody (Mouse mAb) | ServiceBio | GB15001 |
| VDAC(D73D12) Rabbit mAb | CST | 4661 |
| CD11c (3.9) Mouse mAb (FITC Conjugate) | CST | 69627 |
| PE Anti-Mouse CD80 | Proteintech | PE-65076 |
| APC Anti-Mouse CD86 | Proteintech | APC-65068 |

| Antibody | Supplier | Cat. Num. |
|---|---|---|
| CD8α (RPA-T8) Mouse mAb (APC Conjugate) | CST | 64915 |
| CD3 (UCHT1) Mouse mAb (PE Conjugate) | CST | 46233 |
| CD4(RM4-5) Rabbit mAb (FITC Conjugate) | CST | 96127 |
| PerCP/Cyanine5.5 anti-mouse CD45 Antibody | Biolegend | 103132 |
| PE anti-mouse NK1.1 Antibody | Biolegend | 108707 |
| DFNA5(G-9) | Santa | 393162 |
| P-STING(S365) Rabbit mAb | CST | 62912 |
| Anti-NKR-P1C antibody | abcam | ab289542 |
| InvivoMAb mouse IgG2α isotype control | Biocell | BE0085 |
| InvivoMAb anti-mouse NK1.1 | Biocell | BE0036 |

**Supplemental Figures and Figure legends**

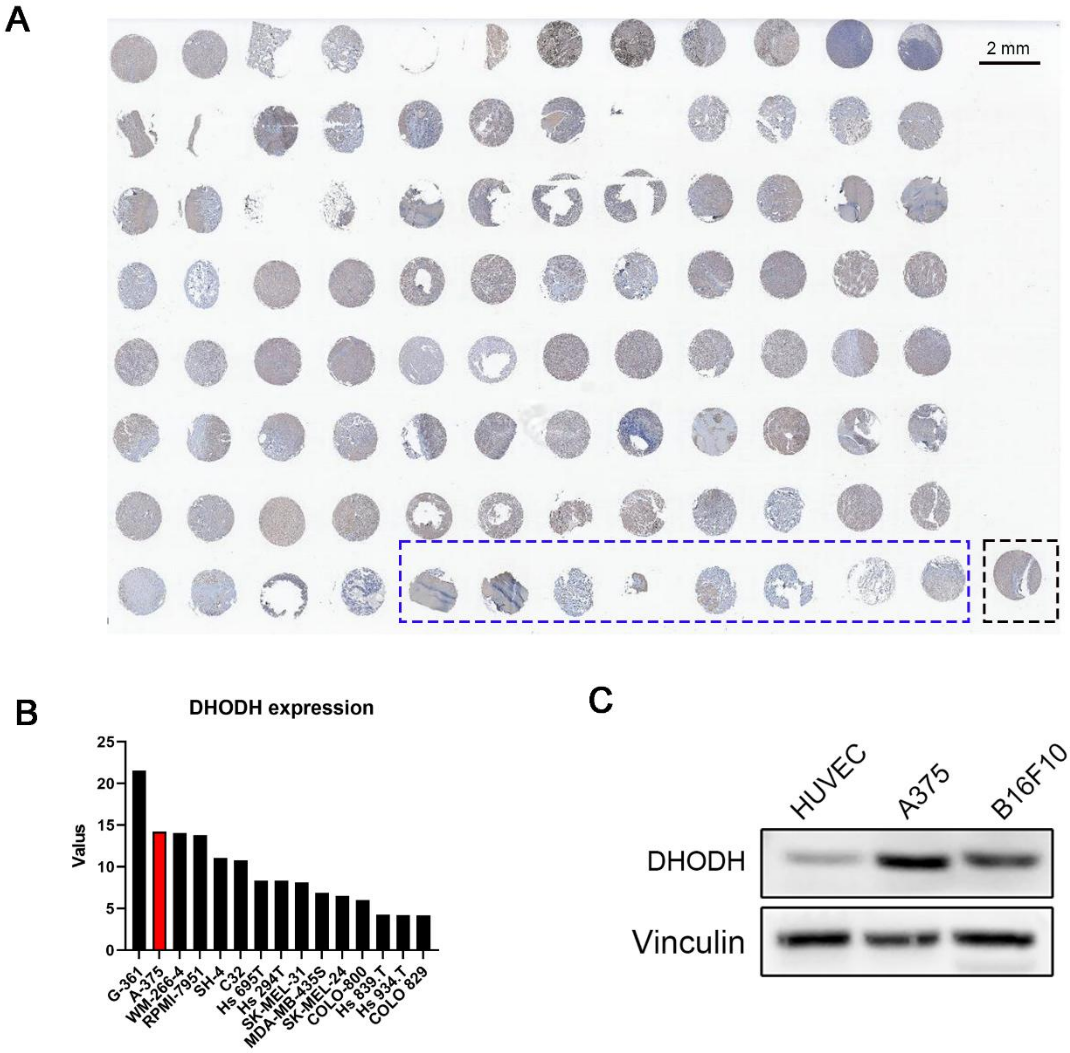


**Fig. S1. Over-expressed DHODH relates to immunosuppressive in melanoma.** (A) Full view of melanoma tissue microarray. (Black bordered rectangle: localization point; Blue bordered rectangle: normal tissues; others: melanoma tissues). (B) DHODH expression levels in different human-derived melanoma cell line. Dates were obtained from Cancer Cell Line Encyclopedia (CCLE). (C) Western blot analysis of DHODH in HUVEC (normal cells), A375 and B16F10 cells.

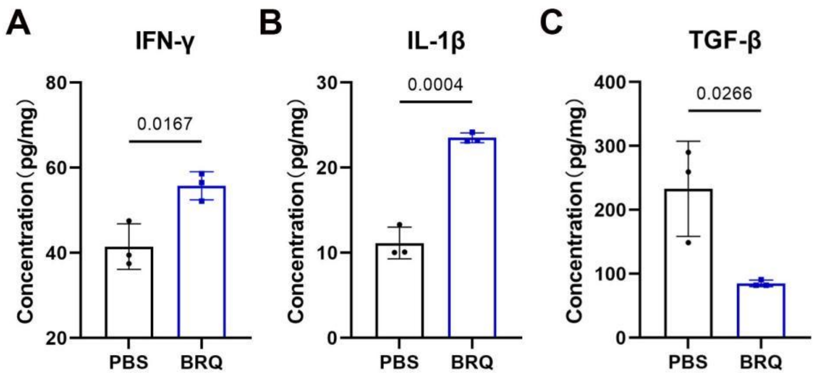


**Fig. S2. BRQ restrains melanoma growth.** (A-C) The concentration of IFN-γ, IL-1β and TGF-β in tumors were analyzed by ELISA (n = 3).

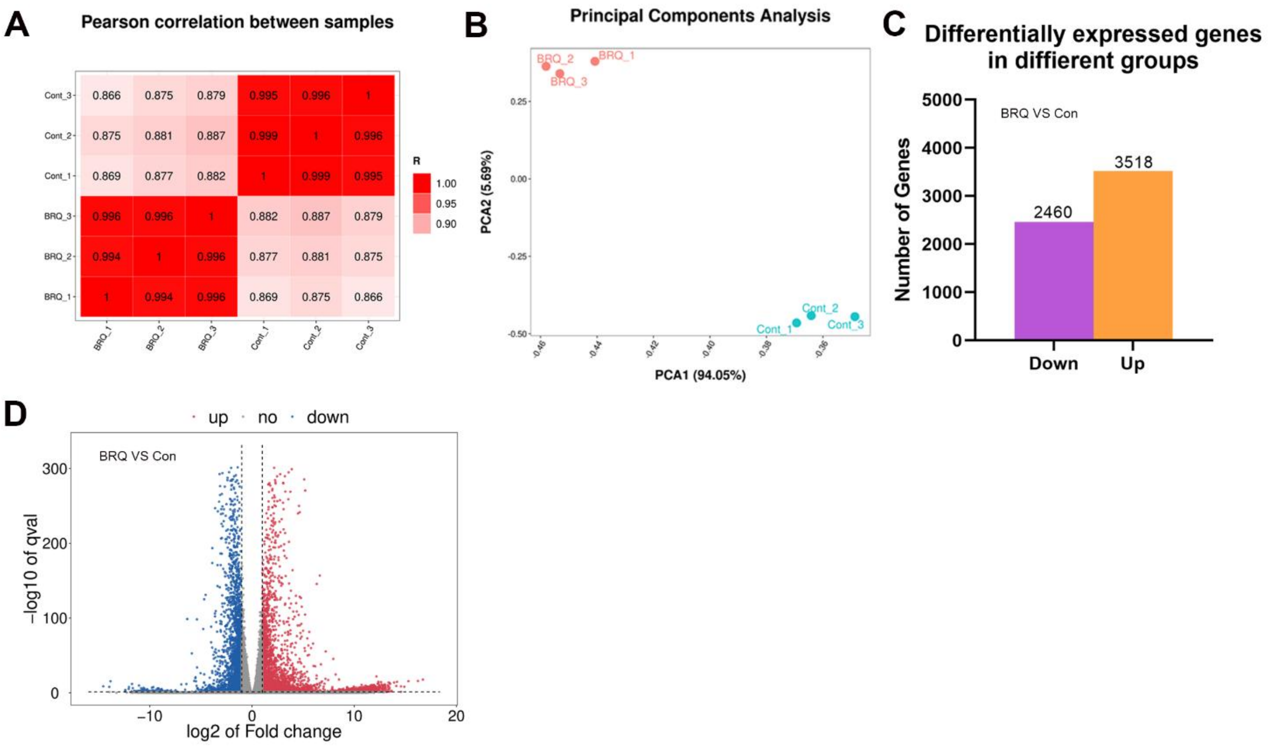


**Fig. S3. RNA-Seq analysis of B16F10 cells treated with BRQ.** (A) Pearson correlation between the BRQ group and the control group. (B) PCA analysis between the BRQ and the control group. (C) The number of DEGs after BRQ treatment. (D) The volcano plot of DEGs in BRQ and the control group.

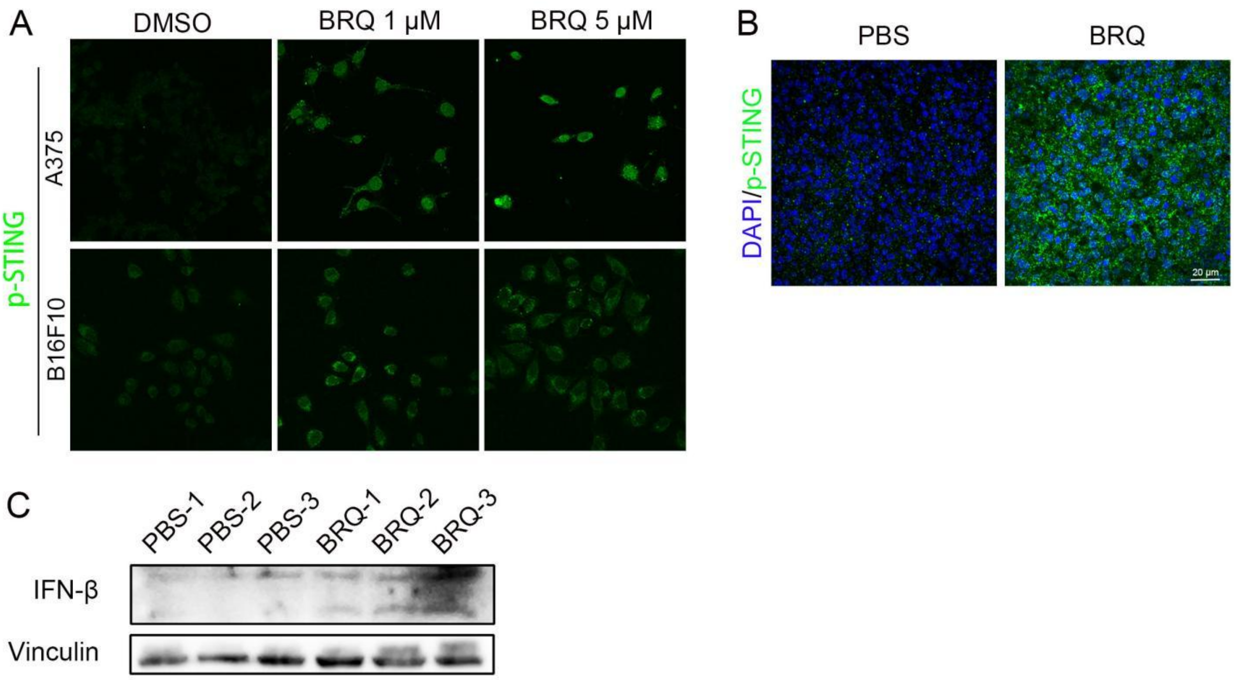


**Fig. S4. BRQ activation of the cGAS-STING pathway enhances the antitumor immunity of NK cells.** (A) CLSM images of p-STING expression in BRQ treatment and control group. (B) Immunostaining of p-STING in tumor tissue (Blue: DAPI; green: p-Sting). (C) Western blot analysis of IFN-β in tumor tissue (n = 3 for each group).

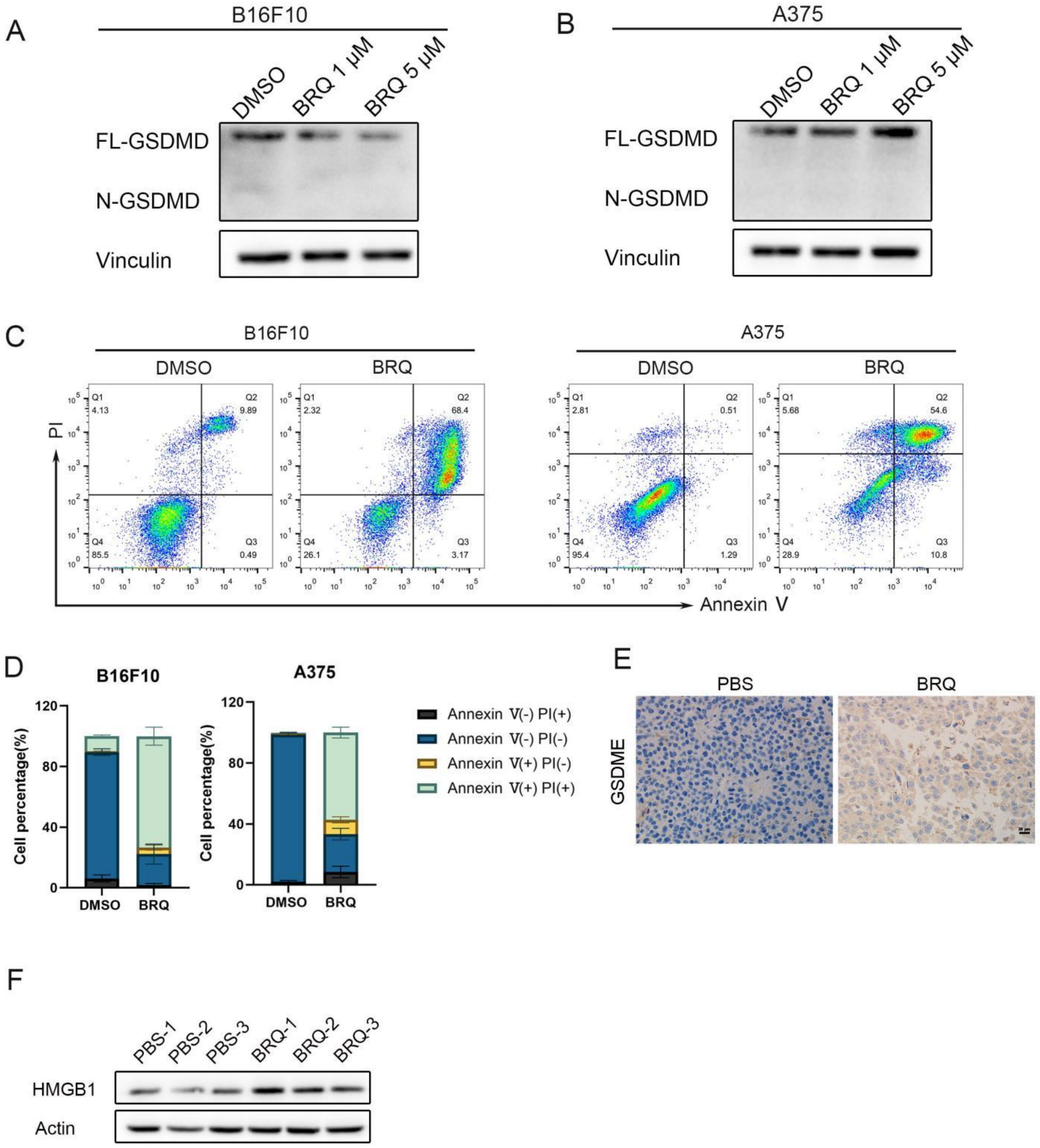


**Fig. S5. BRQ induces pyroptosis in melanoma cells and synergizes with NK cells.** (A-B) Western blot analysis of GSDMD in B16F10 and A375 cells. (C) Flow analysis of cells death after BRQ treatment. (D) Quantification of single PI positive, single FITC-Annexin V positive, and FITC-Annexin V/PI double positive or negative cells. (E) Immunohistochemistry analysis of GSDME expression in tumor tissue in BRQ treatment and control group. (F) Western blot analysis of HMGB1 in tumor tissue (n = 3 for each group).

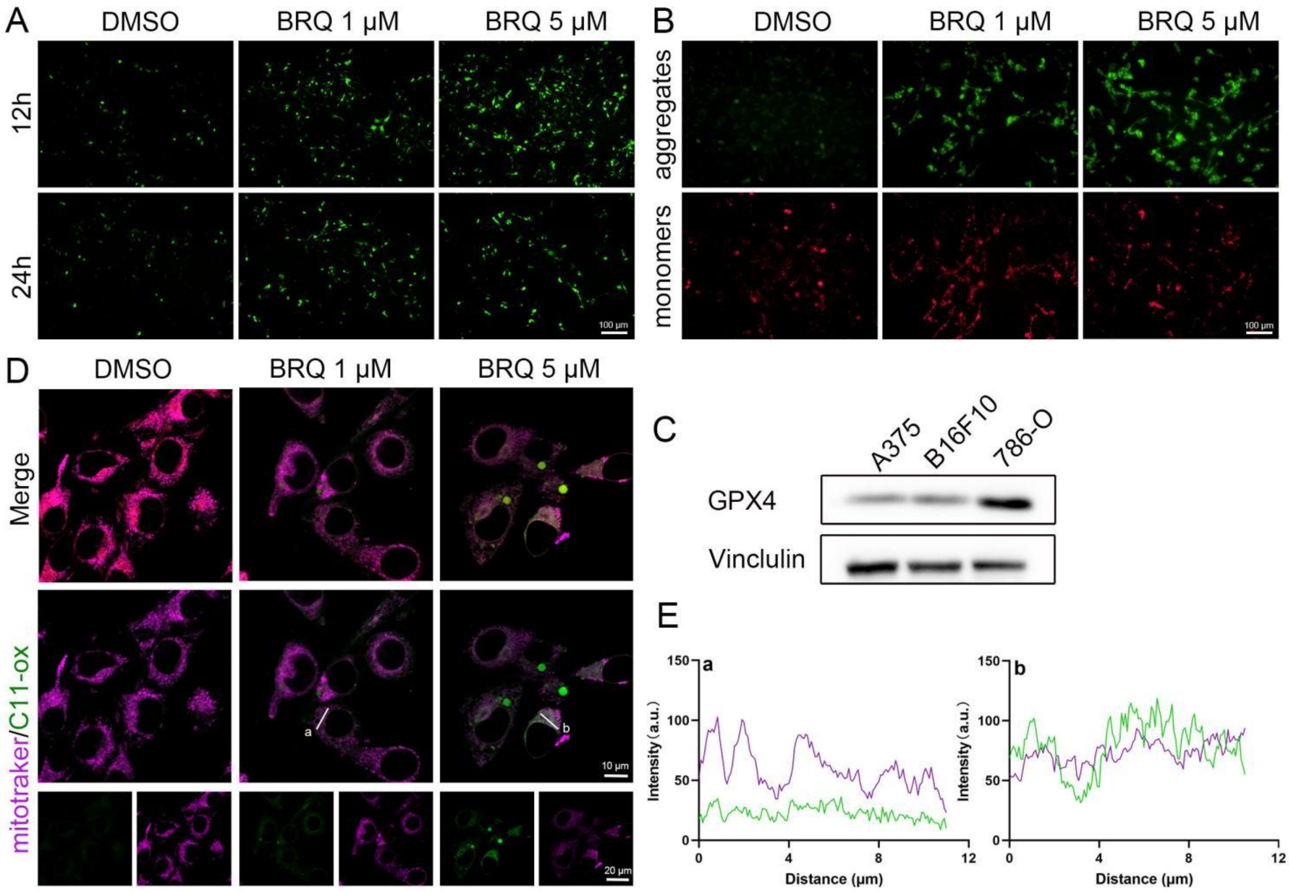


**Fig. S6. BRQ induced mitochondrial oxidative stress and mtDNA released via VDAC.**

(A) ROS was detected by DCFH-DA in B16F10 cells. (B) JC-1 analysis in B16F10 cells. (C) Western blot analysis of GPX4 in A375, B16F10 and 786-O cell lines. (D) Representative images of B16F10 cells co-stained BODIPY C11 (Oxidized BODIPY-green/non oxidized BODIPY-red) and mitochondrial(purple). (E) Colocalization analysis of mitochondrial and Oxidized BODIPY B16F10 cells.

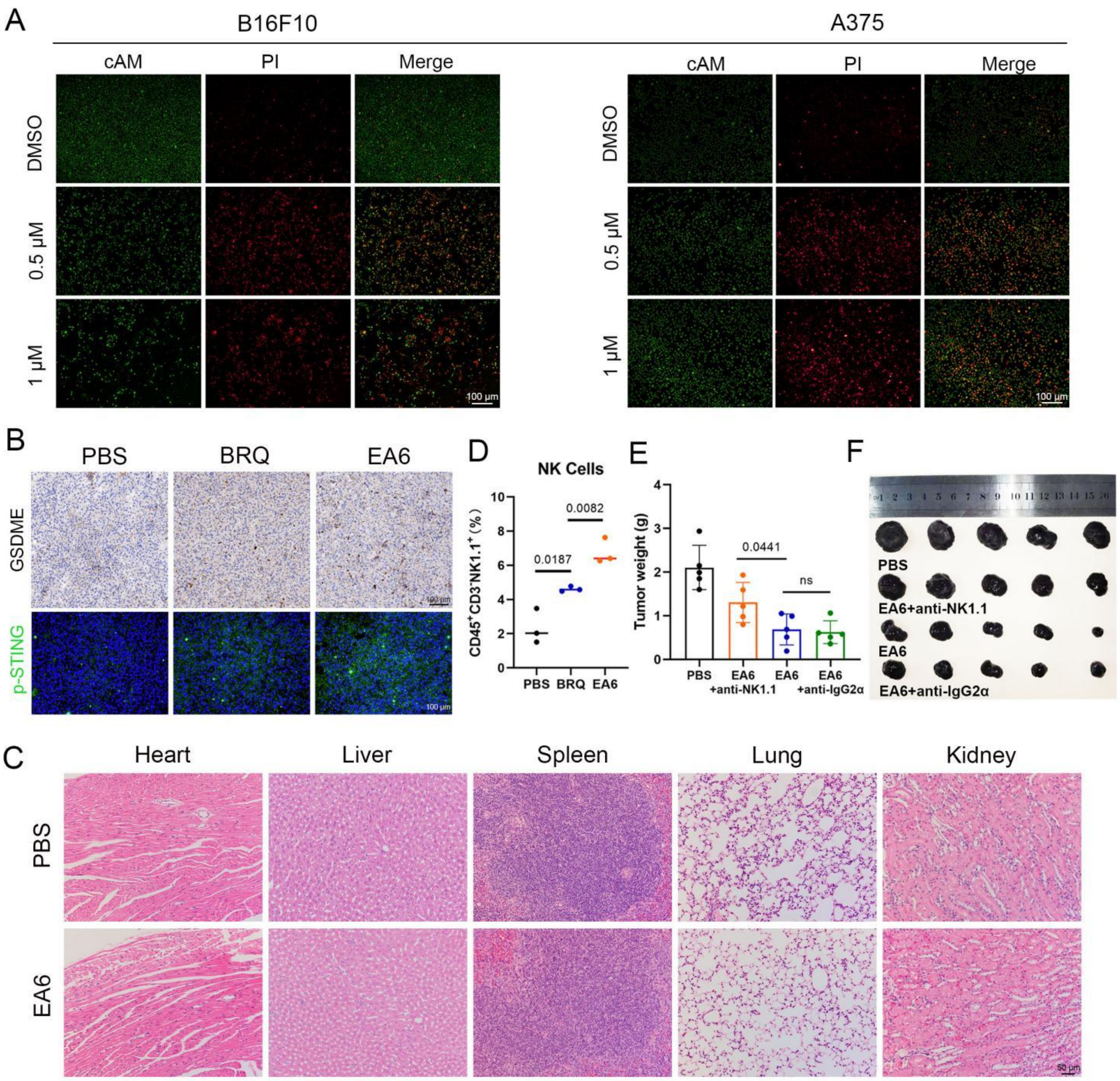


**Fig. S7. EA6, a more effective DHODH inhibitor.**

(A) Live/dead staining analysis in B16F10 and A375 cells. Red indicated dead cells (PI), green indicated live cells (cAM). (B) The expression of GSDEM and p-STING in indicted tumor tissue. (C) H&E stain of heart, liver, spleen, lung and kidney. (D) Quantitative graph of flow cytometry analysis for tumor-infiltrating NK cells. (E) Average tumor weight at Day 17 of each group. two-tailed Student's t-test. (F) Images of isolated tumors for each group in NK-depletion experiment.
