## Supporting Information: Part 2 for "Inhibition of DHODH Activates Pyroptosis and cGAS-STING Pathways to Enhance NK cell Infiltration Mediated Anti-tumor Immunity in Melanoma"

#### Brequinar derivatives synthesis and characterization

**Chemicals and Materials.** All commercial reagents and synthesized materials obtained from Energy-Chemical, Adamas-Beta, or Topbiochem were used without purification unless otherwise specified. Flash-column chromatography was realized with 200-300 mesh silica gel (Qingdao Haiyang Chemical, China). HPLC chromatograms were obtained on a Waters Alliance e2695 with a reverse-phase column (C18, Agilent) (1S0 mm X 4.6 mm) using a mixture of solvents acetonitrile/water (70:30) at a flow rate of 1.0 mL min, peak detection at 254 nm under UV, an injection volume of 10 μL, and an injection time of 10min. $^{1}H$ and $^{13}C$ NMR spectra were recorded on a Bruker ARX 600 MHz spectrometer: chemical shifts and coupling constants (J) were shown in parts per million and in hertz, respectively. HRMS spectra were measured on a Bruker micrOTOF Q spectrometer.

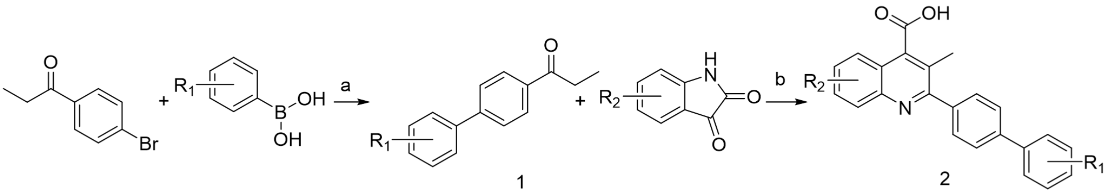

**Reaction and conditions:** (a) $(PPh_3)_4Pd$, $K_2HPO_4$, dioxane:$H_2O$ 3:1; (b) KOH, EtOH:$H_2O$ 3:1.

**General synthesis for intermediate 1**

1-(4-Bromophenyl) propan-1-one ( 1 equiv ), the corresponding phenylboronic acid ( 1 equiv ), $(PPH_3)_4Pd$ ( 0.05 equiv ) and $K_2HPO_4$ ( 2 equiv ) were added in 3:1 1,4- dioxane / $H_2O$ under inert atmosphere . The reaction was heated to 130 $^{o}C$ for reflux for 24 h. After the reaction was complete, the obtained mixture was concentrated, and the mixture was extracted with ethyl acetate three times

(3×). The organic layer was dried with MgSO4. The residue was purified by column chromatography.

**General synthesis for product 2**

The corresponding isatin, KOH, were added in 3:1 EtOH/$H_2O$. The reaction was heated to 100 °C for reflux for 0.5 h. The product from the first step was added. The reaction was heated to 80 °C for reflux for 9 h. The mixture was concentrated, and the mixture was extracted with ethyl acetate three times (3×). The aqueous layer was acidified with HCl until pH 2−3 was reached, the sediment is filtered and washed in deionized water and product was dried under vacuum.

**1-(3',5'-difluoro-[1,1'-biphenyl]-4-yl)propan-1-one**

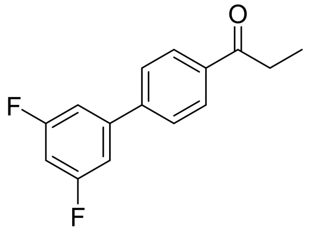

***A2*** The synthesis of general synthesis for intermediate 1, white solid, 75%. $^1$H NMR (62 MHz, DMSO-*d6*): δ 8.24 – 7.74 (m, 4H), 7.71 – 7.53 (m, 1H), 7.52 – 7.04 (m, 2H), 3.10 (q, *J* = 7.1 Hz, 2H), 1.11 (t, *J* = 7.1 Hz, 3H). $^{13}$C NMR (101 MHz, DMSO-*d6*): δ 200.45, 164.65, 164.51, 162.20, 162.07, 143.08, 142.98, 142.88, 142.06, 142.04, 142.01, 136.85, 128.96, 127.67, 110.80, 110.73, 110.61, 110.54, 104.30, 104.05, 103.79, 31.83, 8.53. HRMS (ESI) calcd for $C_{15}H_{12}F_2O$ $[M+H]^+$ 247.0934, found 247.0928.

**1-(2'-chloro-[1,1'-biphenyl]-4-yl)propan-1-one**

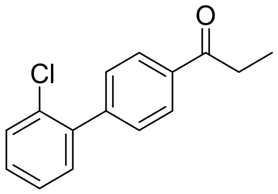

***A3*** The synthesis of general synthesis for intermediate 1, white solid, 77%. $^1$H NMR (62 MHz, DMSO-*d6*): δ 8.24 – 7.87 (m, 2H), 7.75 – 7.29 (m, 6H), 3.10 (q, *J* = 7.1 Hz, 2H), 1.11 (t, *J* = 7.1 Hz, 3H). $^{13}$C NMR (101 MHz, DMSO-*d6*): δ 200.52, 143.54, 139.34, 136.25, 131.84, 131.62, 130.43, 130.25, 130.08, 128.23, 128.11, 31.78, 8.57. HRMS (ESI) calcd for $C_{15}H_{13}ClO$ $[M+H]^+$ 245.0733, found 245.0728.

**1-(3'-fluoro-[1,1'-biphenyl]-4-yl)propan-1-one**

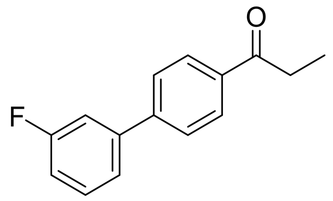

*A4* The synthesis of general synthesis for intermediate 1, white solid, 79%. $^{1}$H NMR (62 MHz, DMSO-*d6*): δ 8.26 – 6.98 (m, 8H), 3.08 (q, *J* = 7.1 Hz, 2H), 1.10 (t, *J* = 7.1 Hz, 3H). $^{13}$C NMR (101 MHz, DMSO-*d6*): δ 200.43, 164.39, 161.97, 143.31, 143.29, 141.87, 141.80, 136.37, 131.54, 131.45, 128.99, 127.53, 123.55, 123.52, 115.62, 115.41, 114.30, 114.08, 31.78, 8.56. HRMS (ESI) calcd for $C_{15}H_{13}ClO$ $[M+H]^+$ 229.1029, found 229.1031.

**1-(2'-methyl-[1,1'-biphenyl]-4-yl)propan-1-one**

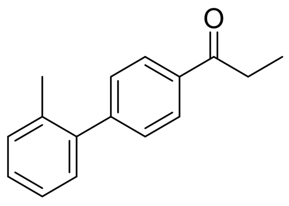

*A6* The synthesis of general synthesis for intermediate 1, white solid, 79%. $^{1}$H NMR (500 MHz, DMSO-*d6*): δ 8.03 (d, J = 8.3 Hz, 2H), 7.49 (d, J = 8.3 Hz, 2H), 7.34 – 7.26 (m, 3H), 7.24 – 7.21 (m, 1H), 3.09 (q, J = 7.2 Hz, 2H), 2.24 (s, 3H), 1.11 (t, J = 7.2 Hz, 3H).8.03 (d, J = 8.3 Hz, 2H), 7.49 (d, J = 8.3 Hz, 2H), 7.34 – 7.26 (m, 3H), 7.24 – 7.21 (m, 1H), 3.09 (q, J = 7.2 Hz, 2H), 2.24 (s, 3H), 1.11 (t, J = 7.2 Hz, 3H). $^{13}$C NMR (101 MHz, DMSO-*d6*) δ: 200.46, 146.33, 140.76, 135.59, 135.13, 130.96, 129.80, 129.77, 128.34, 128.28, 126.53, 31.69, 20.55, 8.59. HRMS (ESI) calcd for $C_{16}H_{16}O$ $[M+H]^+$ 225.1279, found 225.1269.

**1-(2',4'-difluoro-[1,1'-biphenyl]-4-yl)propan-1-one**

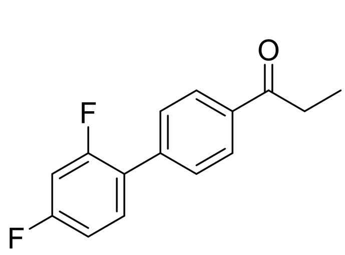

*A8* The synthesis of general synthesis for intermediate 1, white solid, 79%. $^{1}$H NMR (62 MHz, DMSO-*d6*): δ 8.26 – 7.91 (m, 2H), 7.88 – 7.00 (m, 5H), 3.09 (q, *J* = 7.1 Hz, 2H), 1.10 (t, *J* = 7.1 Hz, 3H). $^{13}$C NMR (101 MHz, DMSO-*d6*): δ 200.47, 163.94, 163.82, 161.48, 161.35, 160.97, 160.85, 158.49, 158.37, 139.07, 139.05, 136.26, 132.57, 132.52, 132.47, 132.43, 129.52, 129.49, 128.63, 124.44, 124.41, 124.31, 124.28, 112.87, 112.83, 112.66, 112.62, 105.43, 105.17,

104.91, 31.78, 8.55. HRMS (ESI) calcd for $C_{15}H_{12}F_2O$ $[M+H]^+$ 247.0934, found 247.0931

**1-(2',6'-difluoro-[1,1'-biphenyl]-4-yl)propan-1-one**

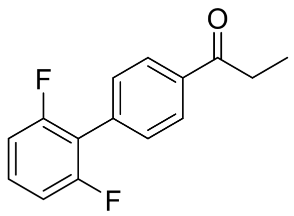

***A11*** The synthesis of general synthesis for intermediate 1, white solid, 72%. $^1$H NMR (62 MHz, DMSO-*d6*):δ 8.21 – 7.94 (m, 2H), 7.77 – 7.03 (m, 5H), 3.10 (q, J = 7.1 Hz, 2H), 1.11 (t, J = 7.1 Hz, 3H). $^{13}$C NMR (101 MHz, DMSO-*d6*): δ 200.52, 160.95, 160.88, 158.49, 158.42, 136.80, 133.55, 131.29, 131.18, 131.08, 130.98, 130.96, 130.94, 128.36, 117.51, 117.32, 117.14, 112.76, 112.70, 112.57, 112.51, 31.79, 8.52. HRMS (ESI) calcd for $C_{15}H_{12}F_2O$ $[M+H]^+$ 247.0934, found 247.0916.

**1-(2'-chloro-6'-fluoro-[1,1'-biphenyl]-4-yl)propan-1-one**

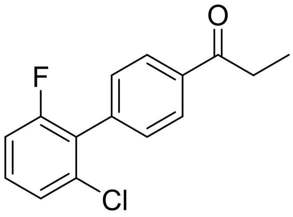

***A12*** The synthesis of general synthesis for intermediate 1, white solid, 72%. $^1$H NMR (400 MHz, DMSO-*d6*): δ 8.07 (d, J = 8.6 Hz, 2H), 7.53 – 7.43 (m, 4H), 7.38 – 7.31 (m, 1H), 3.09 (q, J = 7.1 Hz, 2H), 1.11 (t, J = 7.1 Hz, 3H). $^{13}$C NMR (101 MHz, DMSO-*d6*): δ 200.40, 161.17, 158.71, 137.01, 136.83, 133.45, 133.41, 131.21, 131.12, 130.84, 130.82, 128.25, 127.99, 127.80, 126.21, 126.18, 115.41, 115.19, 31.77, 8.46. HRMS (ESI) calcd for $C_{15}H_{12}ClFO$ $[M+H]^+$ 263.0639, found 263.0616.

**1-(2'-chloro-6'-fluoro-[1,1'-biphenyl]-4-yl)propan-1-one**

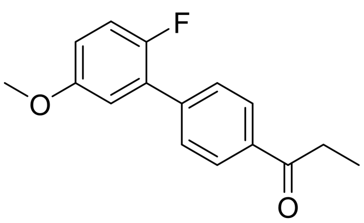

***A14*** The synthesis of general synthesis for intermediate 1, white solid, 79%. $^1$H NMR (62 MHz, DMSO-*d6*): δ 8.06 (d, *J* = 8.5 Hz, 2H), 7.70 (m, 2H), 7.48 – 6.82 (m, 3H), 3.81 (s, 3H), 3.09 (q, *J* = 7.1 Hz, 2H), 1.11 (t, *J* = 7.1 Hz, 3H). $^{13}$C NMR (101 MHz, DMSO-*d6*): δ 200.52, 156.24, 156.23, 155.03, 152.66, 139.95, 139.94, 136.28, 129.60, 129.57, 128.55, 128.28, 128.14, 117.57, 117.33, 115.83, 115.75, 115.63, 115.60, 56.21, 31.79, 8.57. HRMS (ESI) calcd for $C_{16}H_{15}FO_2$ $[M+H]^+$ 259.1134, found

259.1122.

**2-(3',5'-difluoro-[1,1'-biphenyl]-4-yl)-6-fluoro-3-methylquinoline-4-carboxylic acid**

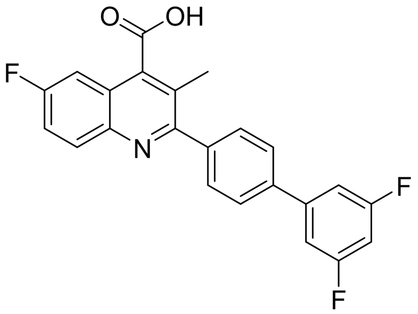

***AA2*** The synthesis of General synthesis for product 2, white solid, 55%. $^{1}$H NMR (400 MHz, DMSO-*d6*): δ 8.01 (dd, J = 9.3, 5.6 Hz, 1H), 7.88 (d, J = 8.4 Hz, 2H), 7.68 (d, J = 8.4 Hz, 2H), 7.65 – 7.50 (m, 4H), 7.27 (tt, J = 9.3, 2.4 Hz, 1H), 2.38 (s, 3H). $^{13}$C NMR (101 MHz, DMSO-*d6*): δ 164.69, 164.55, 162.25, 162.11, 161.26, 159.47, 158.84, 143.92, 143.82, 143.72, 143.59, 141.50, 137.64, 132.07, 131.97, 130.17, 127.06, 124.58, 124.48, 123.23, 119.20, 118.95, 110.46, 110.40, 110.28, 110.21, 109.90, 109.67, 103.64, 103.38, 103.13, 18.08. 394.1148. HRMS (ESI) calcd for $C_{23}H_{14}F_{3}NO_{2}$ $[M+H]^{+}$ 394.1055, found 394.1148. HPLC purity at 254 nm, 99.52%.

**2-(2'-chloro-[1,1'-biphenyl]-4-yl)-6-fluoro-3-methylquinoline-4-carboxylic acid**

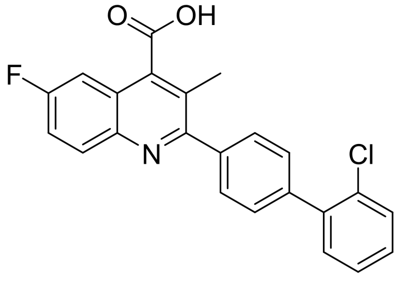

***AA3*** The synthesis of General synthesis for product 2, yellow solid, 50%. $^{1}$H NMR (400 MHz, DMSO-*d6*): δ 8.02 (dd, *J* = 9.2, 5.8 Hz, 1H), 7.67 (d, *J* = 8.2 Hz, 2H), 7.65 – 7.58 (m, 3H), 7.56 (d, *J* = 8.2 Hz, 2H), 7.51 (dd, *J* = 7.2, 2.4 Hz, 1H), 7.45 (dtd, *J* = 12.9, 7.2, 1.9 Hz, 2H), 2.41 (s, 3H).$^{13}$C NMR (101 MHz, DMSO-*d6*): δ 170.66, 161.32, 159.71, 159.68, 158.89, 143.59, 140.53, 139.92, 138.87, 132.12, 132.03, 131.83, 130.41, 129.82, 129.40, 129.34, 128.09, 124.48, 124.38, 123.41, 119.27, 119.02, 109.73, 109.50, 18.11. HRMS (ESI) calcd for $C_{23}H_{15}ClFNO_{2}$ $[M+H]^{+}$ 392.0854, found 392.0832. HPLC purity at 254 nm, 99.16%

**2-(2'-chloro-[1,1'-biphenyl]-4-yl)-6-methoxy-3-methylquinoline-4-carboxylic acid**

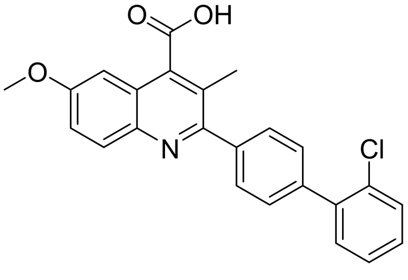

***BA3*** The synthesis of General synthesis for product 2, yellow solid, 44%. $^{1}$H NMR (400 MHz, DMSO-*d6*): δ 7.94 (d, *J* = 9.2 Hz, 1H),

7.71 – 7.67 (m, 2H), 7.61 (dd, *J* = 7.1, 2.1 Hz, 1H), 7.58 – 7.55 (m, 2H), 7.54 – 7.39 (m, 5H), 7.13 (d, *J* = 2.8 Hz, 1H), 3.89 (s, 3H), 2.41 (s, 3H). $^{13}$C NMR (101 MHz, DMSO-*d*6): δ 169.94, 158.14, 157.44, 142.41, 140.30, 139.92, 138.84, 132.04, 131.83, 131.16, 130.41, 129.82, 129.46, 129.42, 128.09, 124.20, 124.03, 121.99, 103.23, 55.93, 18.09. HRMS (ESI) calcd for $C_{24}H_{18}ClNO_3$ $[M+H]^+$ 404.1053, found 404.1037. HPLC purity at 254 nm, 98.01%.

**2-(3'-fluoro-[1,1'-biphenyl]-4-yl)-6-methoxy-3-methylquinoline-4-carboxylic acid**

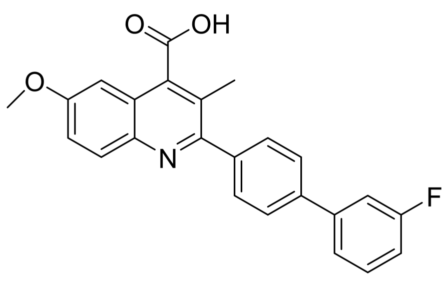

***BA4*** The synthesis of General synthesis for product 2, white solid, 47%. $^{1}$H NMR (400 MHz, DMSO-*d*6): δ 7.98 (d, *J* = 9.2 Hz, 1H), 7.87 – 7.83 (m, 2H), 7.72 (dt, *J* = 6.4, 1.9 Hz, 2H), 7.62 (ddd, *J* = 7.5, 4.2, 1.8 Hz, 2H), 7.55 (td, *J* = 8.1, 6.2 Hz, 1H), 7.45 (dd, *J* = 9.2, 2.7 Hz, 1H), 7.27 – 7.21 (m, 1H), 7.08 (d, *J* = 2.7 Hz, 1H), 3.91 (s, 3H), 2.43 (s, 3H).$^{13}$C NMR (101 MHz, DMSO-*d*6): δ 169.50, 167.43, 164.45, 162.03, 158.43, 157.35, 142.64, 142.56, 142.37, 140.67, 140.33, 138.94, 138.92, 132.19, 131.98, 131.46, 131.38, 130.29, 129.13, 127.01, 124.68, 124.01, 123.31, 123.29, 122.23, 114.97, 114.76, 114.05, 113.83, 102.72, 55.96, 18.11. HRMS (ESI) calcd for $C_{24}H_{18}FNO_3$ $[M+H]^+$ 388.1349, found 388.1325. HPLC purity at 254 nm, 91.66%.

**2-(3',5'-difluoro-[1,1'-biphenyl]-4-yl)-3,6-dimethylquinoline-4-carboxylic acid**

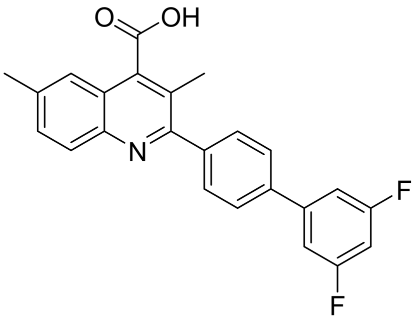

***CA2*** The synthesis of General synthesis for product 2, white solid, 50%. $^{1}$H NMR (400 MHz, DMSO-*d*6): δ 7.96 (d, *J* = 8.5 Hz, 1H), 7.91 – 7.87 (m, 2H), 7.75 – 7.71 (m, 2H), 7.62 (dd, *J* = 8.7, 1.9 Hz, 1H), 7.58 – 7.53 (m, 3H), 7.27 (tt, *J* = 9.3, 2.3 Hz, 1H), 2.55 – 2.53 (m, 3H), 2.43 (s, 3H). $^{13}$C NMR (101 MHz, DMSO-*d*6): δ 169.47,

164.69, 164.55, 162.25, 162.11, 158.90, 144.83, 143.84, 143.75, 143.65, 141.18, 140.90, 137.89, 137.86, 137.84, 137.68, 132.12, 131.97, 130.28, 129.48, 129.13, 127.12, 124.22, 123.46, 122.93, 110.49, 110.42, 110.31, 110.24, 103.68, 103.42, 103.17, 21.87, 18.01. HRMS (ESI) calcd for $C_{24}H_{17}F_2NO_2$ $[M+H]^+$ 390.1306, found 390.1293. HPLC purity at 254 nm, 89.85%.

**2-(3'-fluoro-[1,1'-biphenyl]-4-yl)-3,6-dimethylquinoline-4-carboxylic acid**

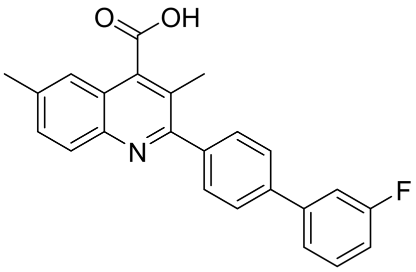

***CA4*** The synthesis of General synthesis for product 2, yellow solid, 59%. $^{1}$H NMR (400 MHz, DMSO-*d6*): δ 7.82 (dd, *J* = 8.4, 1.5 Hz, 3H), 7.69 – 7.65 (m, 3H), 7.61 (ddd, *J* = 7.7, 4.0, 1.8 Hz, 2H), 7.55 (td, *J* = 8.0, 6.1 Hz, 1H), 7.49 (dd, *J* = 8.6, 2.0 Hz, 1H), 7.26 – 7.21 (m, 1H), 2.47 (s, 3H), 2.36 (s, 3H). $^{13}$C NMR (101 MHz, DMSO-*d6*): δ 171.13, 164.45, 162.03, 158.99, 145.05, 142.77, 142.69, 141.41, 138.56, 135.39, 131.46, 131.37, 131.01, 130.17, 128.90, 126.88, 125.59, 123.73, 123.28, 123.25, 121.73, 114.88, 114.67, 114.00, 113.78, 21.83, 18.04. HRMS (ESI) calcd for $C_{24}H_{18}FNO_2$ $[M+H]^+$ 372.1400, found 372.1388. HPLC purity at 254 nm, 96.11%.

**2-(2',4'-difluoro-[1,1'-biphenyl]-4-yl)-3,6-dimethylquinoline-4-carboxylic acid**

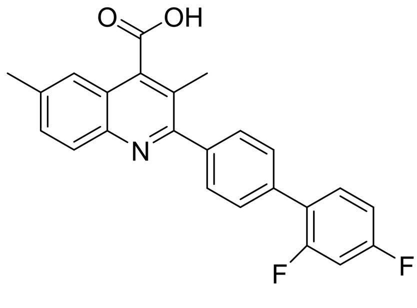

***CA8*** The synthesis of General synthesis for product 2, yellow solid, 54%. $^{1}$H NMR (400 MHz, DMSO-*d6*): δ 7.95 (d, *J* = 8.5 Hz, 1H), 7.74 – 7.72 (m, 2H), 7.67 (dq, *J* = 8.9, 1.7 Hz, 3H), 7.63 (dd, *J* = 8.7, 1.9 Hz, 1H), 7.56 (t, *J* = 1.4 Hz, 1H), 7.42 (ddd, *J* = 11.6, 9.3, 2.6 Hz, 1H), 7.27 – 7.22 (m, 1H), 2.54 (s, 3H), 2.43 (s, 3H). $^{13}$C NMR (101 MHz, DMSO-*d6*): δ 169.49, 167.43, 159.03, 144.83, 140.19, 137.63, 134.62, 132.53, 132.48, 132.43, 132.38, 132.12, 131.99, 129.91, 129.46, 129.14, 128.99, 128.96, 125.01, 124.88, 124.16, 123.49, 122.92, 112.79,

112.58, 105.36, 105.10, 104.84, 21.87, 18.04. HRMS (ESI) calcd for $C_{24}H_{17}F_2NO_2$ $[M+H]^+$ 390.1306, found 390.1292. HPLC purity at 254 nm, 97.10%.

**6-chloro-2-(3',5'-difluoro-[1,1'-biphenyl]-4-yl)-3-methylquinoline-4-carboxylic acid**

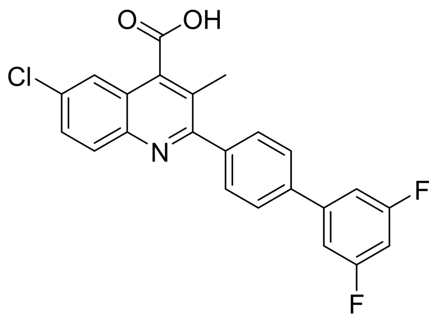

***EA2*** The synthesis of General synthesis for product 2, white solid, 54%. $^1$H NMR (400 MHz, DMSO-*d6*): δ 8.00 (d, *J* = 2.4 Hz, 1H), 7.95 (d, *J* = 9.0 Hz, 1H), 7.89 – 7.85 (m, 2H), 7.69 – 7.64 (m, 3H), 7.54 (dt, *J* = 7.3, 2.2 Hz, 2H), 7.27 (tt, *J* = 9.3, 2.3 Hz, 1H), 2.38 (s, 3H). $^{13}$C NMR (101 MHz, DMSO-*d6*): δ 170.80, 164.68, 164.55, 162.24, 162.11, 160.44, 149.43, 144.81, 143.89, 143.79, 143.70, 141.52, 137.66, 131.29, 130.63, 130.15, 129.37, 127.04, 125.75, 124.69, 123.03, 110.45, 110.38, 110.27, 110.20, 103.65, 103.39, 103.14, 18.09. HRMS (ESI) calcd for $C_{23}H_{14}ClF_2NO_2$ $[M-H]^-$ 408.0603, found 408.0576. HPLC purity at 254 nm, 99.21%.

**6-chloro-2-(2'-chloro-[1,1'-biphenyl]-4-yl)-3-methylquinoline-4-carboxylic acid**

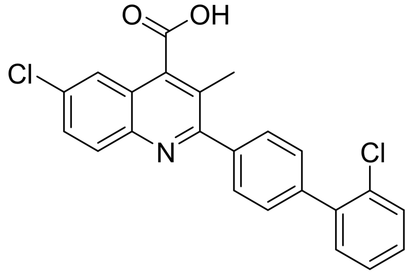

***EA3*** The synthesis of General synthesis for product 2, white solid, 64%. $^1$H NMR (400 MHz, DMSO-*d6*): δ 7.97 (d, J = 2.4 Hz, 1H), 7.94 (d, J = 8.9 Hz, 1H), 7.69 – 7.63 (m, 3H), 7.61 (dd, J = 7.3, 1.9 Hz, 1H), 7.59 – 7.54 (m, 2H), 7.53 – 7.43 (m, 3H), 2.40 (s, 3H). $^{13}$C NMR (101 MHz, DMSO-*d6*): δ 170.45, 160.70, 144.85, 140.75, 139.94, 138.84, 132.03, 131.82, 131.24, 130.42, 129.83, 129.38, 129.32, 129.25, 128.10, 125.88, 124.77, 122.70, 18.10. HRMS (ESI) calcd for $C_{23}H_{15}Cl_2NO_2$ $[M-H]^-$ 406.0402, found 406.0362. HPLC purity at 254 nm, 97.17%.

**6-chloro-2-(3'-fluoro-[1,1'-biphenyl]-4-yl)-3-methylquinoline-4-carboxylic acid**

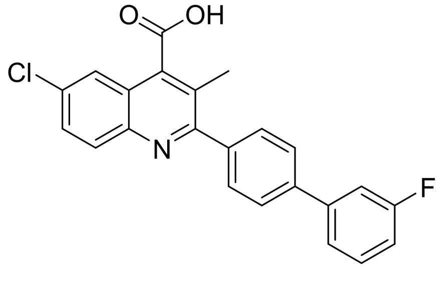

***EA4*** The synthesis of General synthesis for product 2, white

solid, 64%. $^{1}$H NMR (400 MHz, DMSO-*d6*): δ 8.13 – 8.08 (m, 1H), 7.87 (d, J = 8.4 Hz, 2H), 7.82 (d, J = 8.5 Hz, 2H), 7.75 (d, J = 8.3 Hz, 2H), 7.66 – 7.61 (m, 2H), 7.55 (td, J = 7.8, 5.9 Hz, 1H), 7.28 – 7.22 (m, 1H), 2.46 (s, 3H). $^{13}$C NMR (101 MHz, DMSO-*d6*): δ 168.78, 164.45, 162.03, 160.73, 144.61, 142.52, 142.45, 140.66, 139.83, 139.36, 139.34, 132.53, 131.99, 131.49, 131.40, 130.56, 130.26, 127.09, 126.07, 123.68, 123.44, 123.36, 123.34, 115.07, 114.86, 114.10, 113.88, 18.25. HRMS (ESI) calcd for $C_{23}H_{15}ClFNO_2$ $[M+H]^+$ 392.0854, found 392.0846. HPLC purity at 254 nm, 89.58%.

**6-chloro-3-methyl-2-(2'-methyl-[1,1'-biphenyl]-4-yl)quinoline-4-carboxylic acid**

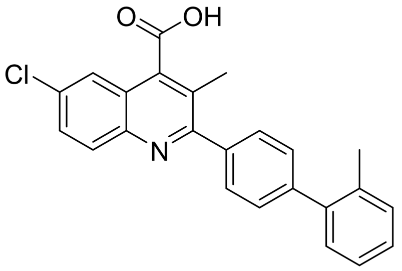

***EA6*** The synthesis of General synthesis for product 2, white solid, 49%. $^{1}$H NMR (500 MHz, DMSO-*d6*): δ 8.12 (d, J = 8.8 Hz, 1H), 7.84 (dd, J = 8.8, 2.3 Hz, 1H), 7.81 (d, J = 2.3 Hz, 1H), 7.73 – 7.70 (m, 2H), 7.52 – 7.49 (m, 2H), 7.36 – 7.29 (m, 4H), 2.49 (s, 3H), 2.32 (s, 3H). $^{13}$C NMR (101 MHz, DMSO-*d6*): δ 170.82, 160.87, 149.91, 144.84, 141.45, 141.38, 139.89, 135.26, 131.23, 130.90, 130.41, 130.01, 129.32, 129.24, 129.08, 127.94, 126.51, 125.86, 124.72, 122.88, 20.73, 18.15. HRMS (ESI) calcd for $C_{24}H_{18}ClNO_2$ $[M-H]^-$ 386.0948, found 386.0900. HPLC purity at 254 nm, 94.94%.

**6-chloro-2-(2',4'-difluoro-[1,1'-biphenyl]-4-yl)-3-methylquinoline-4-carboxylic acid**

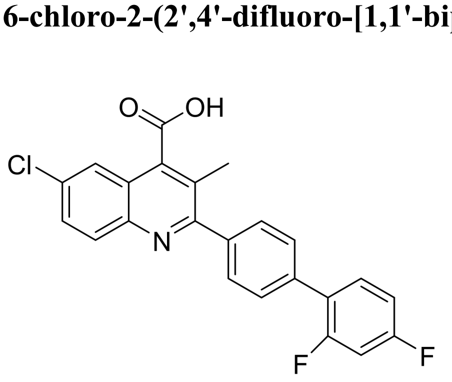

***EA8*** The synthesis of General synthesis for product 2, white solid, 49%. $^{1}$H NMR (400 MHz, DMSO-*d6*): 8.10 (d, *J* = 8.8 Hz, 1H), 7.84 – 7.80 (m, 2H), 7.75 (d, *J* = 8.3 Hz, 2H), 7.69 (ddt, *J* = 8.8, 4.0, 2.3 Hz, 3H), 7.42 (ddd, *J* = 11.5, 9.3, 2.6 Hz, 1H), 7.28 – 7.22 (m, 1H), 2.47 (s, 3H). $^{13}$C NMR (101 MHz, DMSO-*d6*): δ 168.73, 161.08, 160.73, 144.60,

140.56, 139.69, 134.96, 132.58, 132.50, 132.00, 130.60, 129.90, 129.06, 129.03, 126.10, 123.67, 123.42, 112.77, 112.56, 105.37, 105.11, 104.85, 18.25. HRMS (ESI) calcd for $C_{23}H_{14}ClF_2NO_2$ $[M+H]^+$ 410.0759, found 410.0722. HPLC purity at 254 nm, 97.28%.

**2-(2',6'-difluoro-[1,1'-biphenyl]-4-yl)-7-methoxy-3-methylquinoline-4-carboxylic acid**

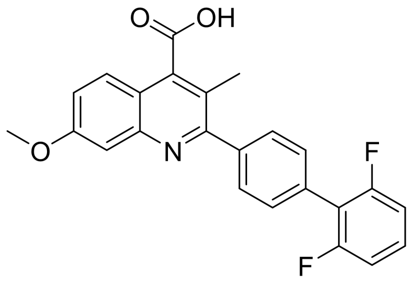

***FA11*** The synthesis of General synthesis for product 2, white solid, 46%. $^1$H NMR (400 MHz, DMSO-*d*6): δ 14.19 (s, 1H), 7.77 – 7.69 (m, 3H), 7.68 – 7.65 (m, 1H), 7.61 (d, *J* = 7.9 Hz, 1H), 7.52 (dt, *J* = 8.2, 1.8 Hz, 1H), 7.45 (t, *J* = 2.2 Hz, 1H), 7.33 (dt, *J* = 9.1, 2.8 Hz, 1H), 7.28 (t, *J* = 8.0 Hz, 1H), 7.02 – 6.90 (m, 1H), 3.93 (s, 3H), 2.41 (s, 3H).$^{13}$C NMR (101 MHz, DMSO-*d*6): δ 172.47, 169.44, 167.43, 160.54, 160.51, 160.08, 147.91, 140.77, 131.99, 131.77, 130.68, 130.34, 129.64, 129.14, 129.04, 128.98, 126.10, 121.78, 120.86, 120.75, 118.01, 117.94, 112.75, 112.49, 107.94, 56.02, 17.74. HRMS (ESI) calcd for $C_{24}H_{17}F_2NO_3$ $[M+H]^+$ 406.1255, found 406.1239. HPLC purity at 254 nm, 96.21%.

**2-(2'-chloro-6'-fluoro-[1,1'-biphenyl]-4-yl)-7-methoxy-3-methylquinoline-4-carboxylic acid**

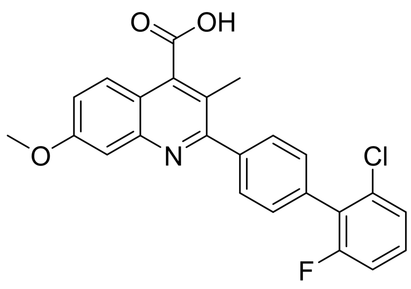

***FA12*** The synthesis of General synthesis for product 2, white solid, 59%. $^1$H NMR (400 MHz, DMSO-*d*6): δ 7.75 (d, J = 9.1 Hz, 1H), 7.68 (q, J = 2.8, 2.1 Hz, 4H), 7.34 (d, J = 2.6 Hz, 1H), 7.31 – 7.26 (m, 1H), 7.17 (dd, J = 9.1, 2.6 Hz, 1H), 7.15 – 7.12 (m, 1H), 7.01 – 6.97 (m, 1H), 3.90 (s, 3H), 3.83 (s, 3H), 2.35 (s, 3H). $^{13}$C NMR (101 MHz, DMSO-*d*6): δ 168.73, 160.73, 144.60, 140.56, 139.69, 134.96, 132.58, 132.00, 130.60, 129.90, 129.06, 129.03, 126.10, 123.67, 123.42, 112.77, 112.56, 105.37, 105.11, 104.85, 18.25. HRMS (ESI) calcd for $C_{24}H_{17}ClFNO_3$ $[M+H]^+$ 422.0959, found 422.0939. HPLC purity at 254 nm, 95.16%.

**2-(2'-fluoro-5'-methoxy-[1,1'-biphenyl]-4-yl)-7-methoxy-3-methylquinoline-4-carboxylic acid**

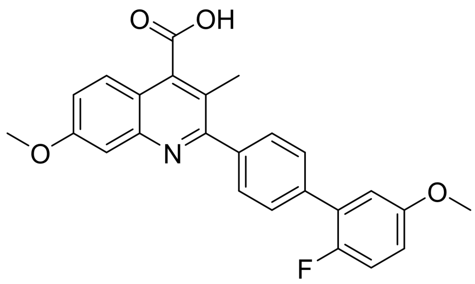

***FA14*** The synthesis of General synthesis for product 2, white solid, 70%. $^{1}$H NMR (400 MHz, DMSO-*d6*): δ 7.75 (d, J = 9.1 Hz, 1H), 7.68 (q, J = 2.8, 2.1 Hz, 4H), 7.34 (d, J = 2.6 Hz, 1H), 7.31 – 7.26 (m, 1H), 7.17 (dd, J = 9.1, 2.6 Hz, 1H), 7.15 – 7.12 (m, 1H), 7.01 – 6.97 (m, 1H), 3.90 (s, 3H), 3.83 (s, 3H), 2.35 (s, 3H).$^{13}$C NMR (101 MHz, DMSO-*d6*): δ 171.00, 159.99, 156.25, 156.23, 155.15, 152.78, 147.97, 141.12, 135.12, 129.72, 128.88, 128.85, 127.71, 120.15, 119.28, 118.69, 117.48, 117.23, 115.60, 115.57, 115.20, 115.13, 107.44, 56.18, 55.82, 17.77. HRMS (ESI) calcd for $C_{25}H_{20}FNO_4$ $[M+H]^+$ 418.1455, found 418.1427. HPLC purity at 254 nm, 97.53%.

**6,7-difluoro-2-(2'-fluoro-5'-methoxy-[1,1'-biphenyl]-4-yl)-3-methylquinoline-4-carboxylic acid**

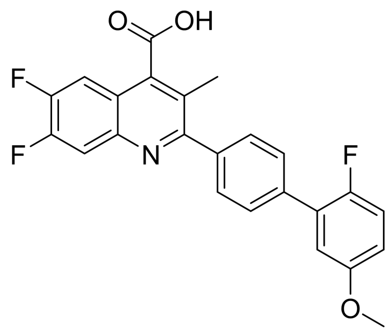

***GA14*** The synthesis of General synthesis for product 2, white solid, 60%. $^{1}$H NMR (400 MHz, DMSO-*d6*): δ 7.71 – 7.65 (m, 4H), 7.56 (d, J = 12.4 Hz, 1H), 7.52 (d, J = 8.5 Hz, 1H), 7.28 (dd, J = 10.3, 9.0 Hz, 1H), 7.13 (dd, J = 6.4, 3.2 Hz, 1H), 6.99 (dt, J = 9.0, 3.5 Hz, 1H), 3.83 (s, 3H), 2.36 (s, 3H). $^{13}$C NMR (101 MHz, DMSO-*d6*): δ 170.08, 159.53, 156.24, 155.15, 153.25, 152.78, 150.77, 149.20, 149.07, 144.59, 140.73, 135.24, 129.73, 128.92, 128.89, 128.77, 121.59, 118.00, 117.92, 117.49, 117.24, 115.60, 115.57, 115.24, 115.16, 110.91, 109.80, 56.18, 17.86. HPLC purity at 254 nm, 96.00%.

#### NMR Spectrum

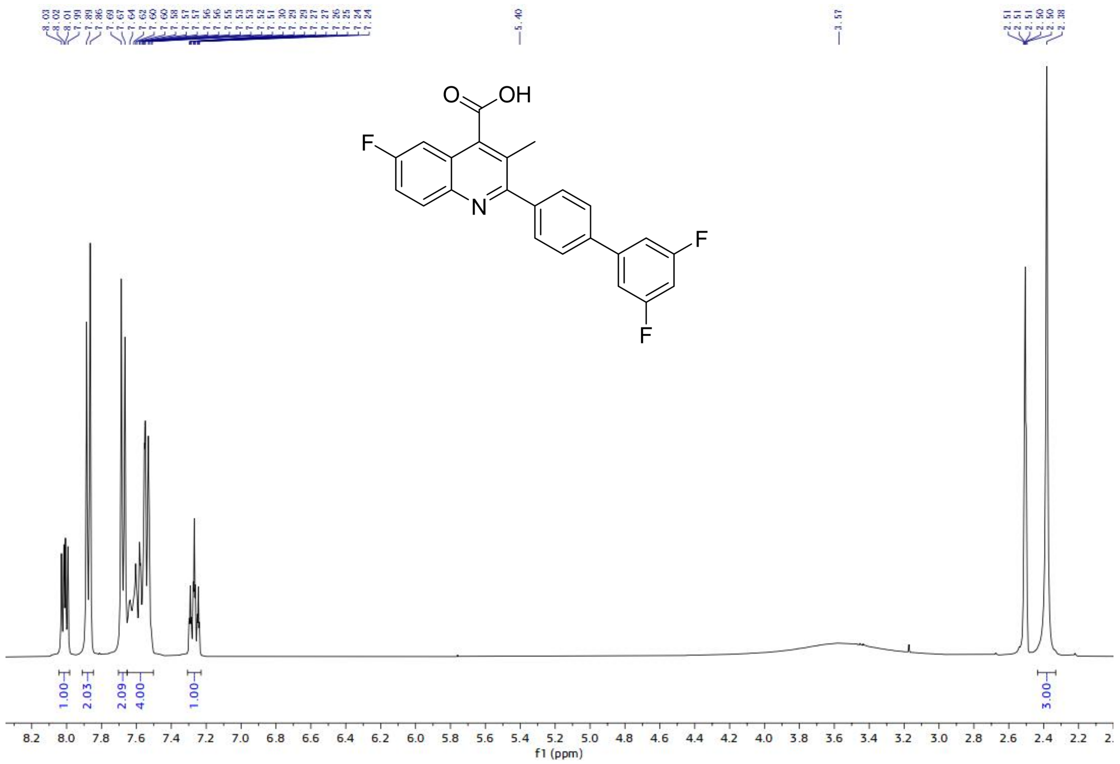

$^{1}$H-NMR spectrum of AA2

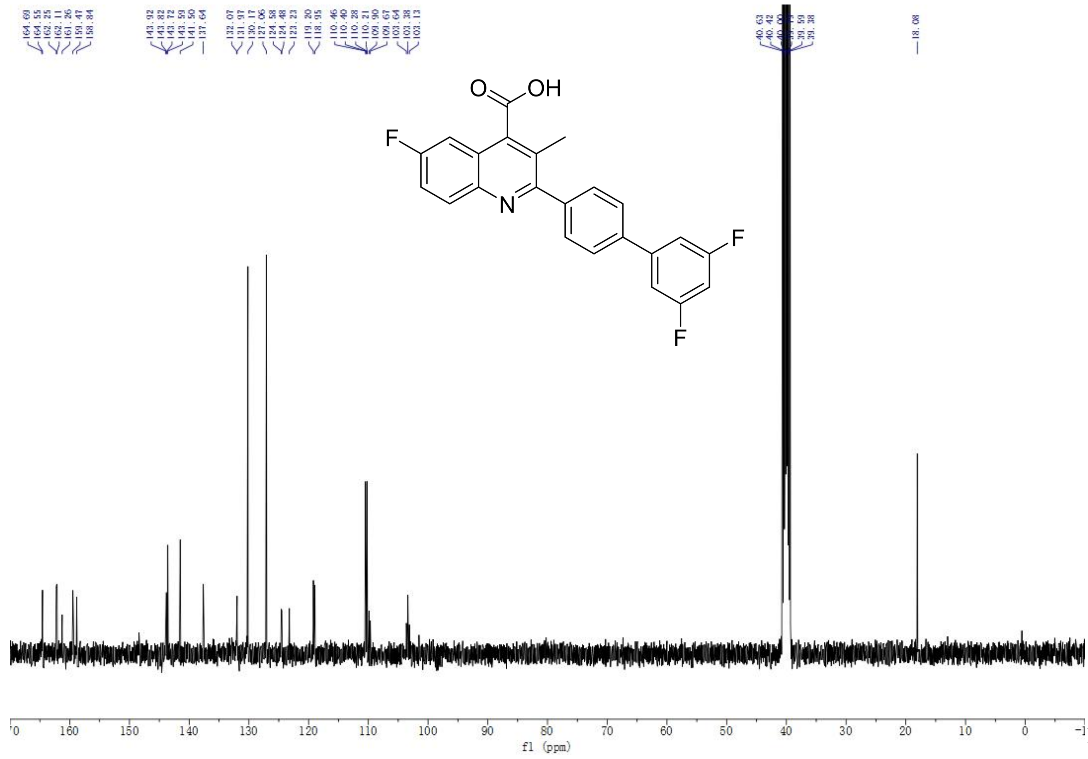

$^{13}$C-NMR spectrum of AA3

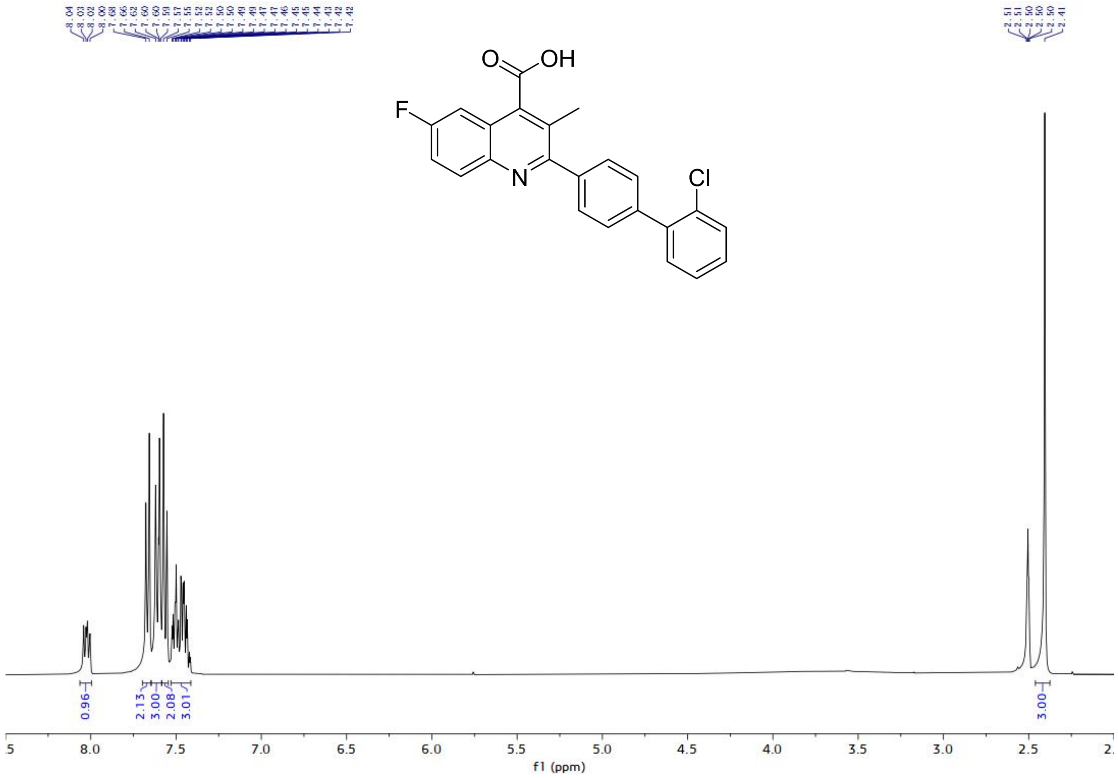

$^{1}$H-NMR spectrum of AA3

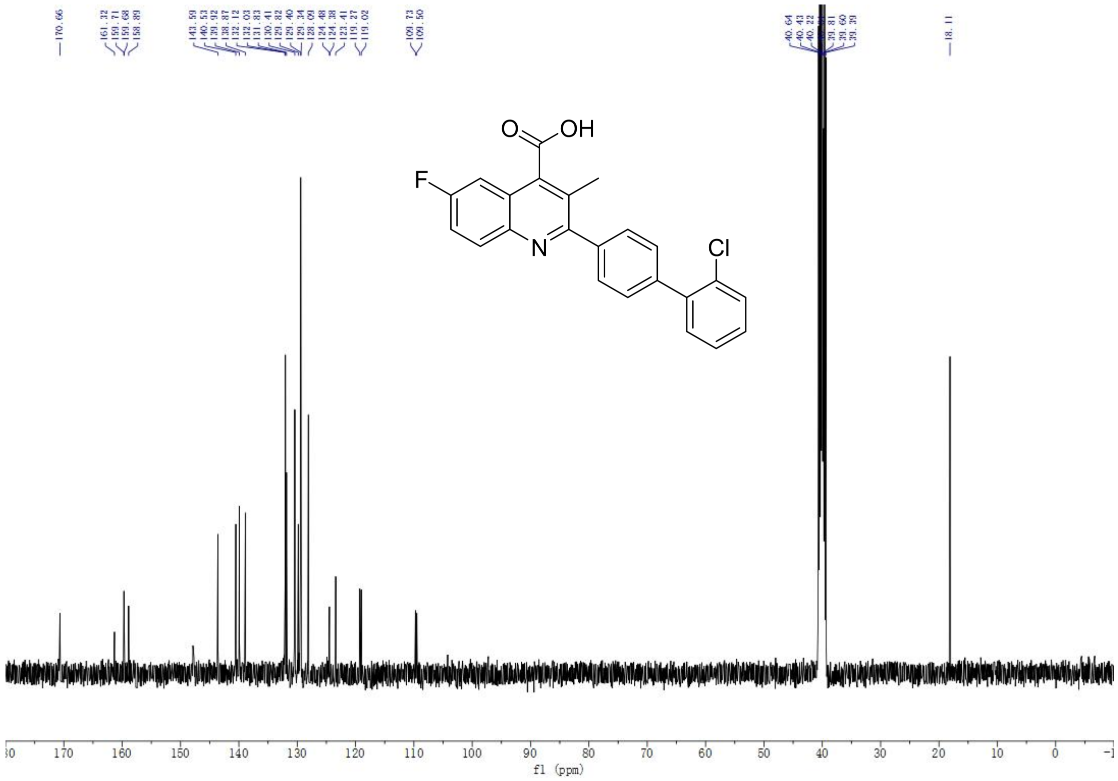

$^{13}$C-NMR spectrum of AA3

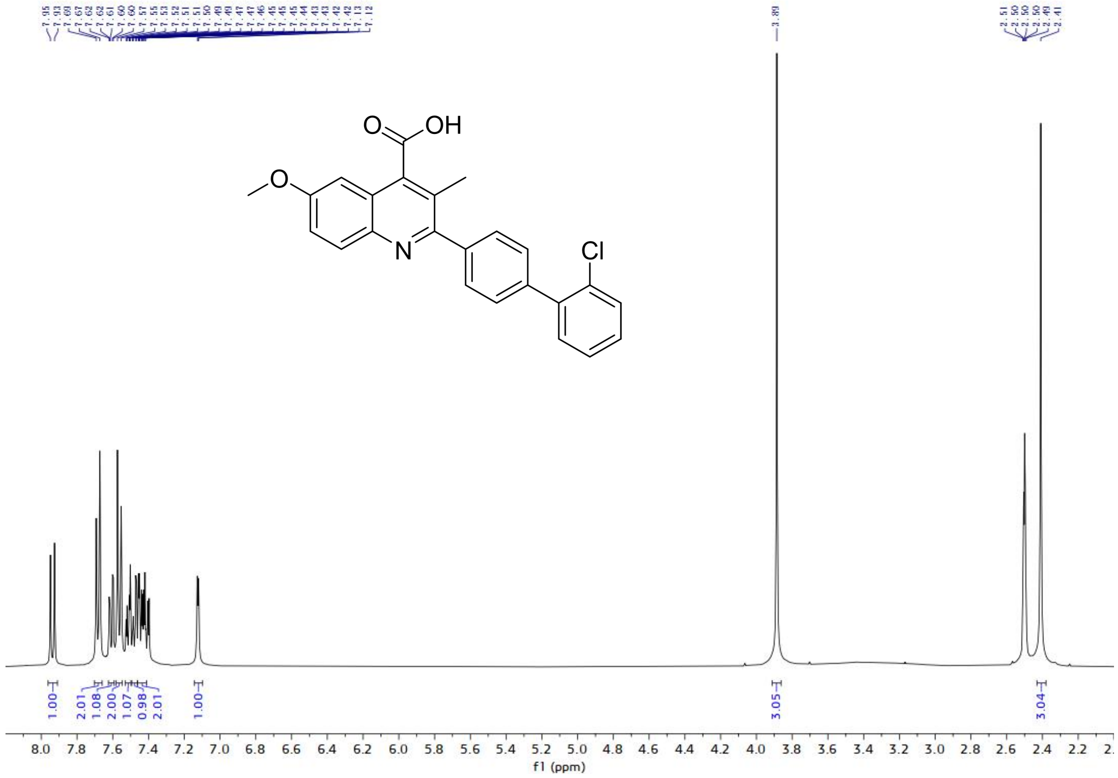

$^{1}$H-NMR spectrum of BA3

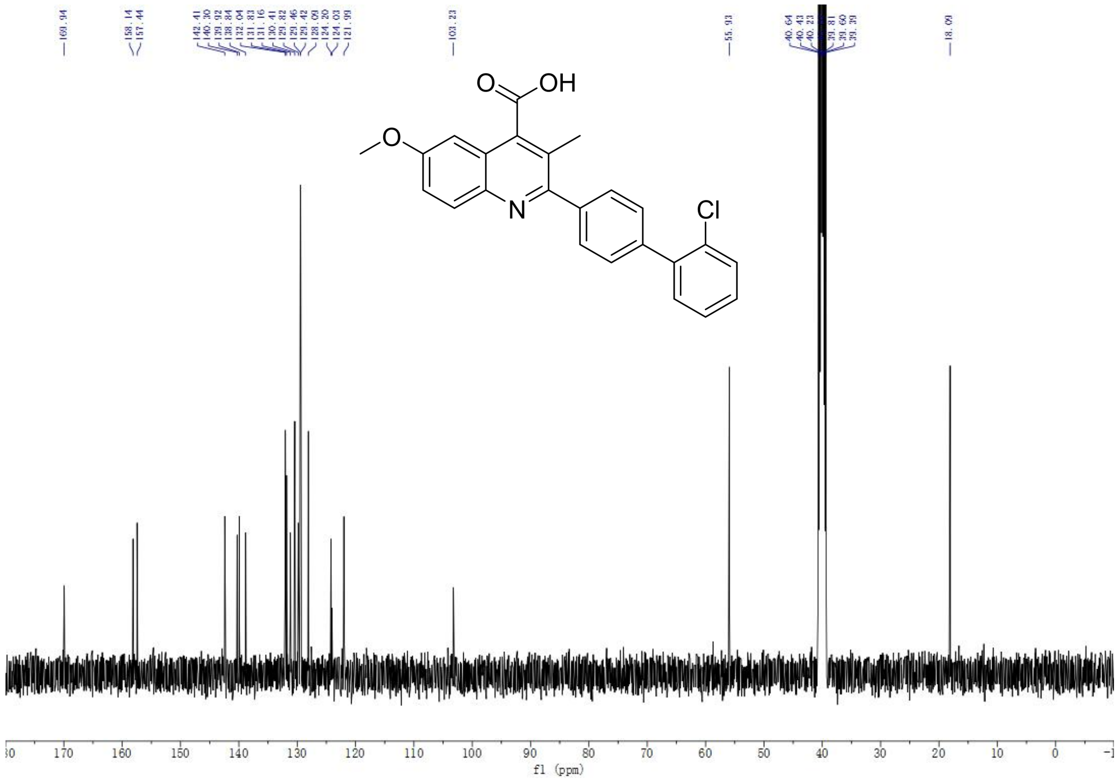

$^{13}$C-NMR spectrum of BA3

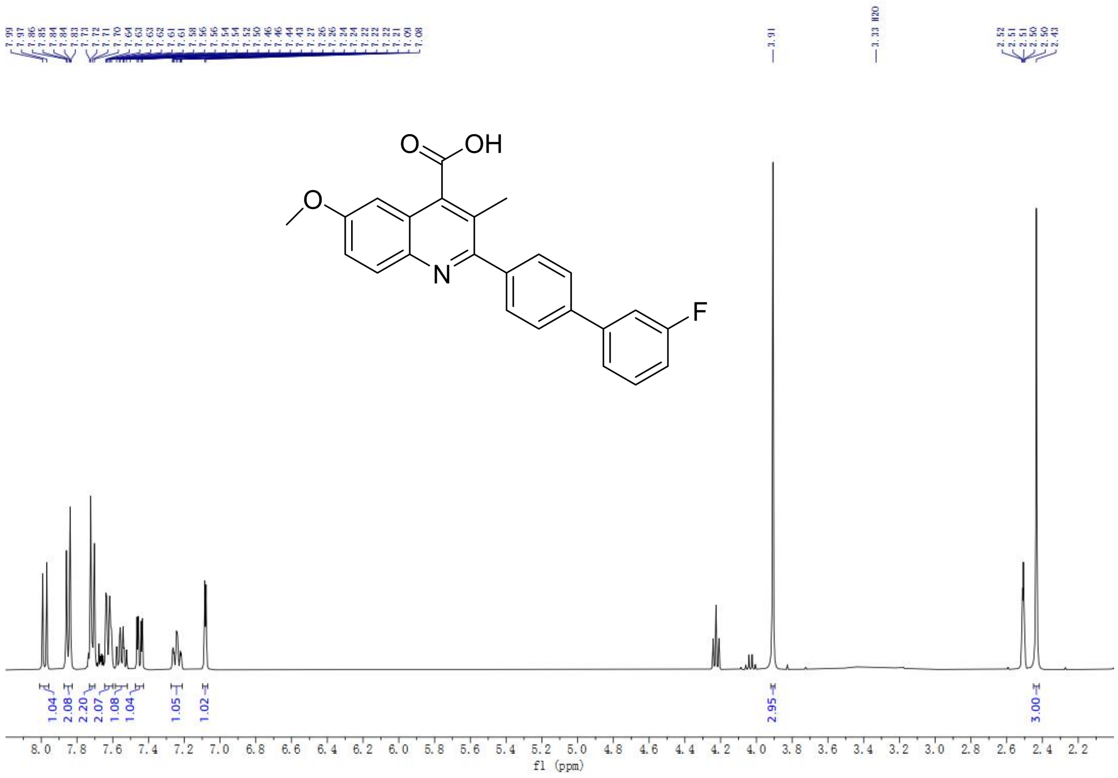

$^{1}H$-NMR spectrum of BA4

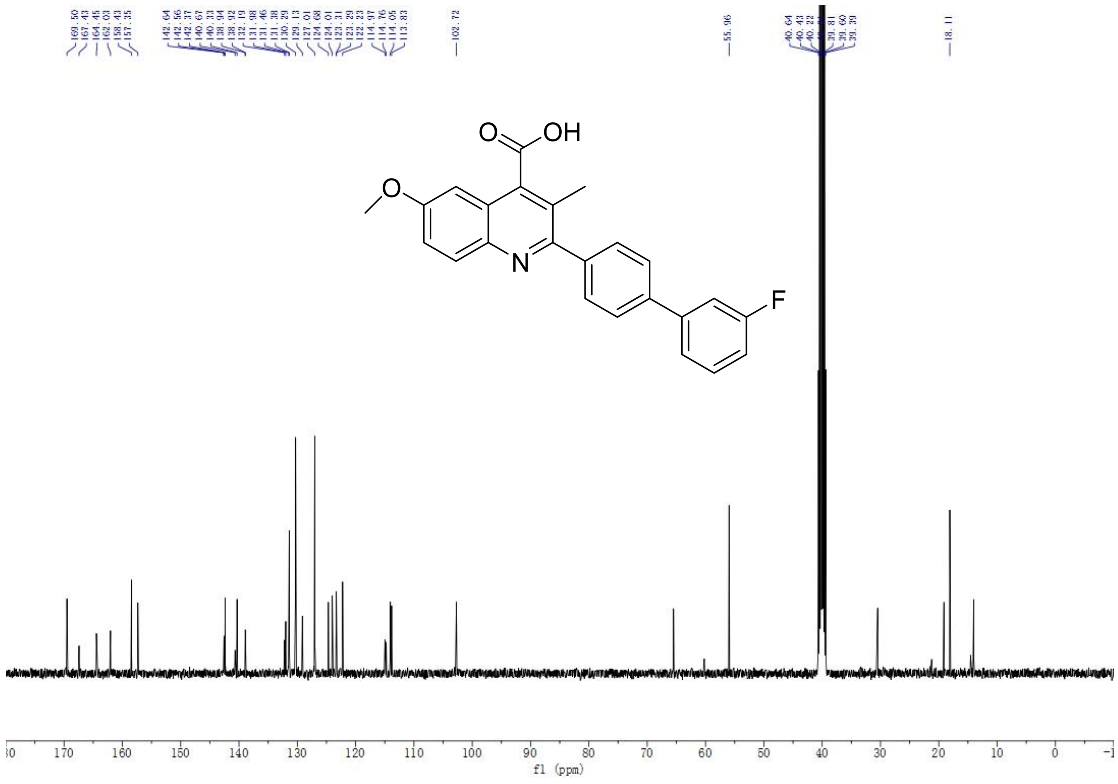

$^{13}C$-NMR spectrum of BA4

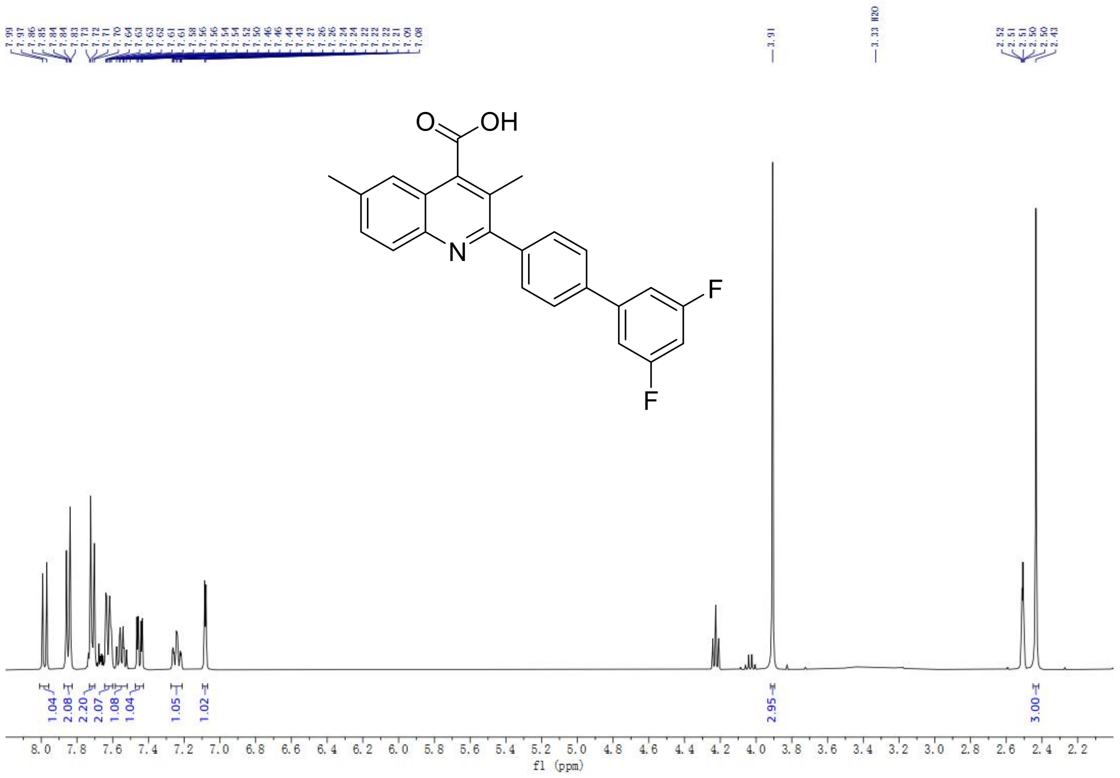

$^{1}$H-NMR spectrum of CA2

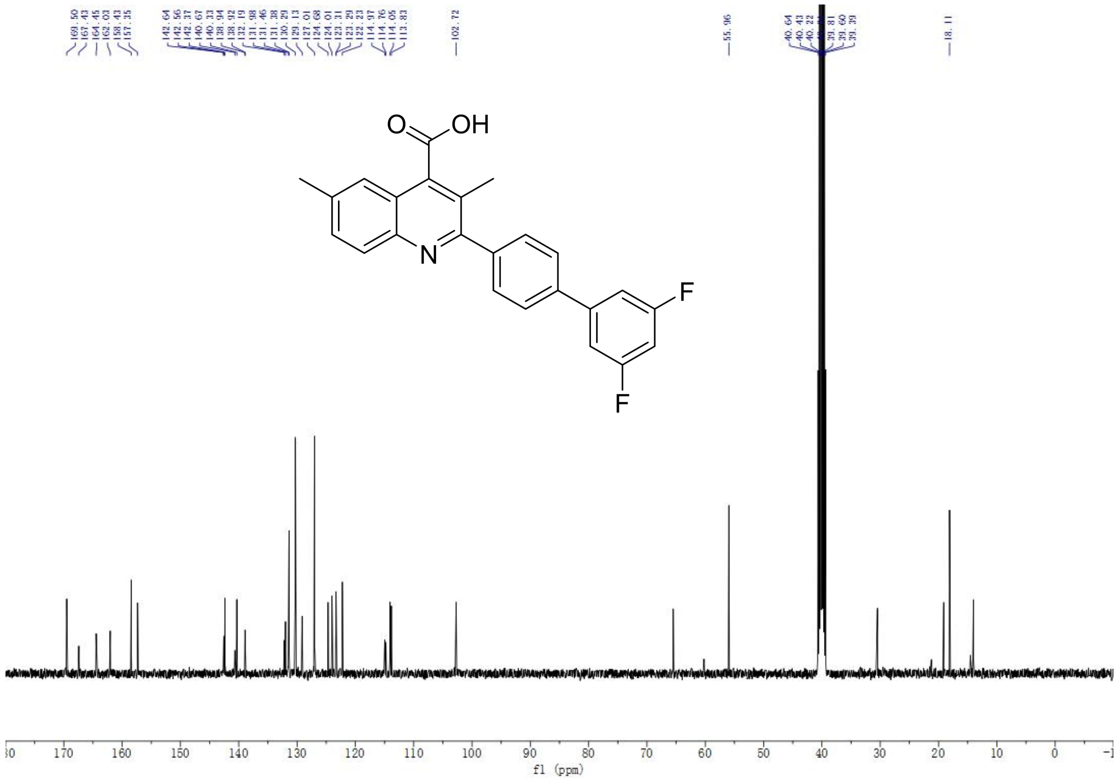

$^{13}$C-NMR spectrum of CA2

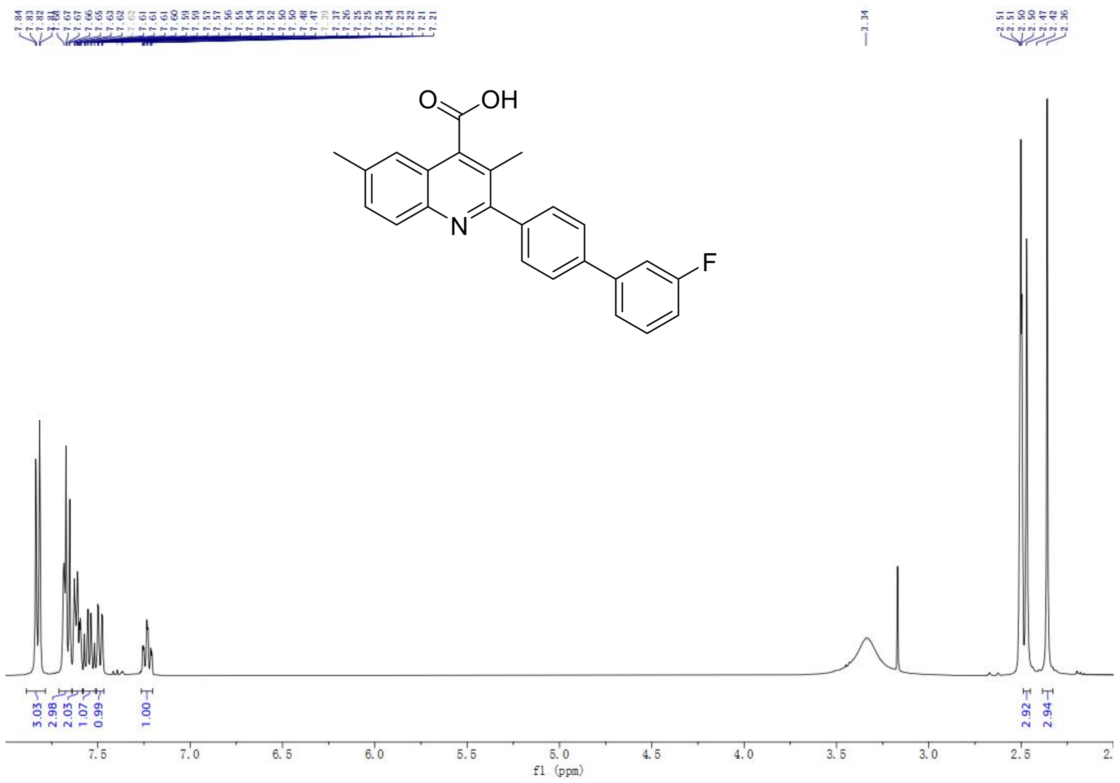

[1]H-NMR spectrum of CA4

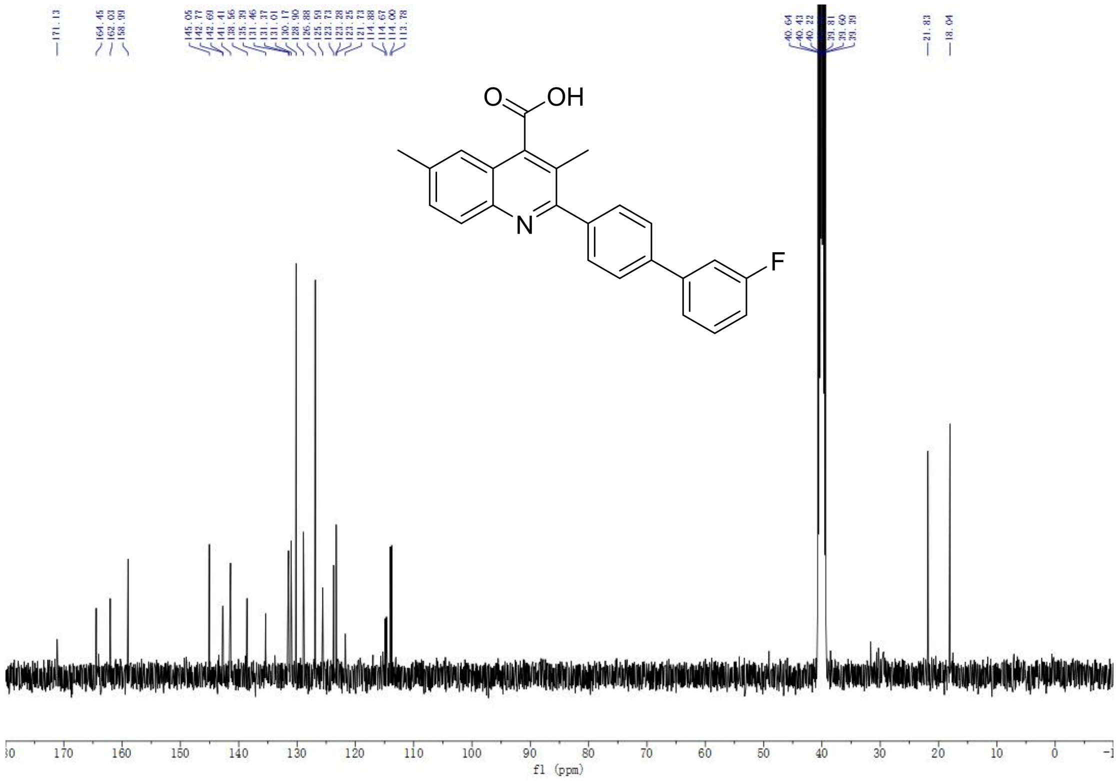

[13]C-NMR spectrum of CA4

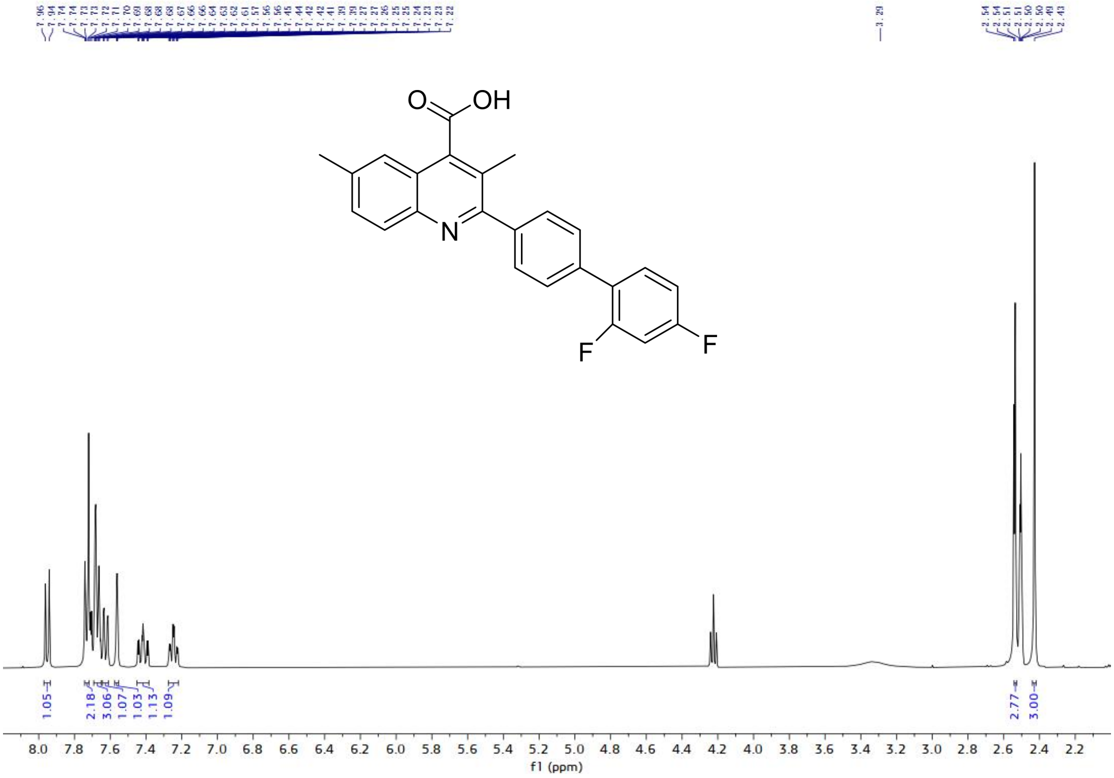

$^1$H-NMR spectrum of CA8

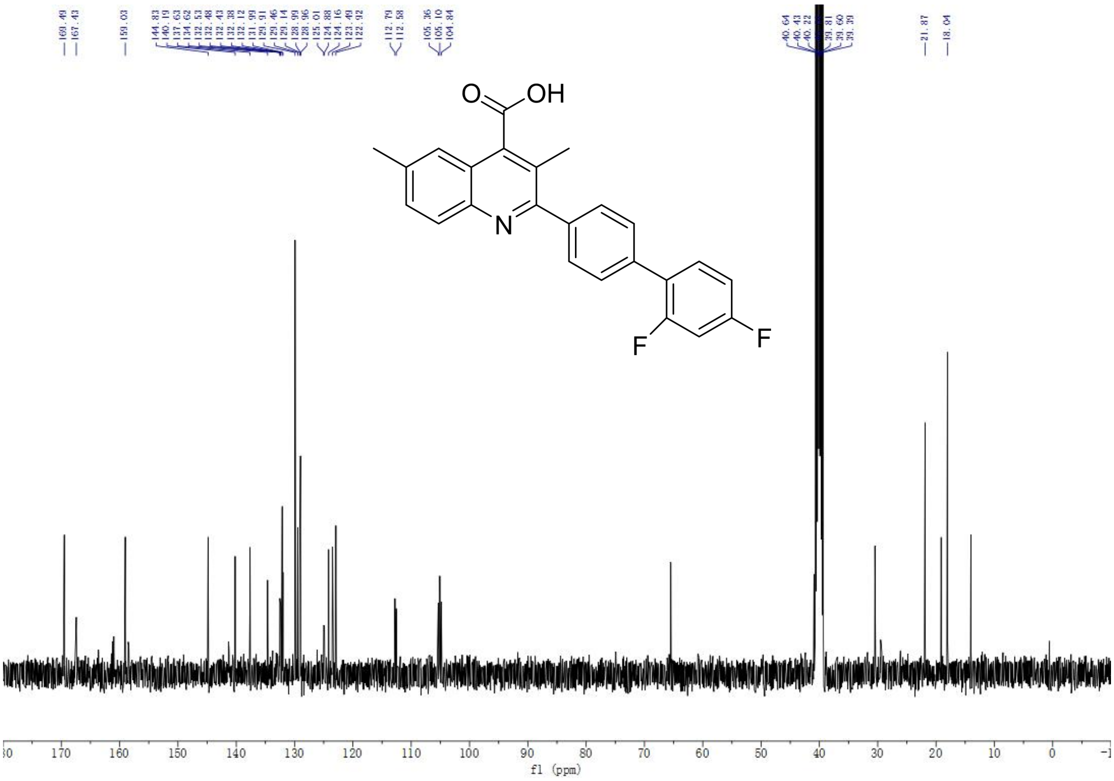

$^{13}$C-NMR spectrum of CA8

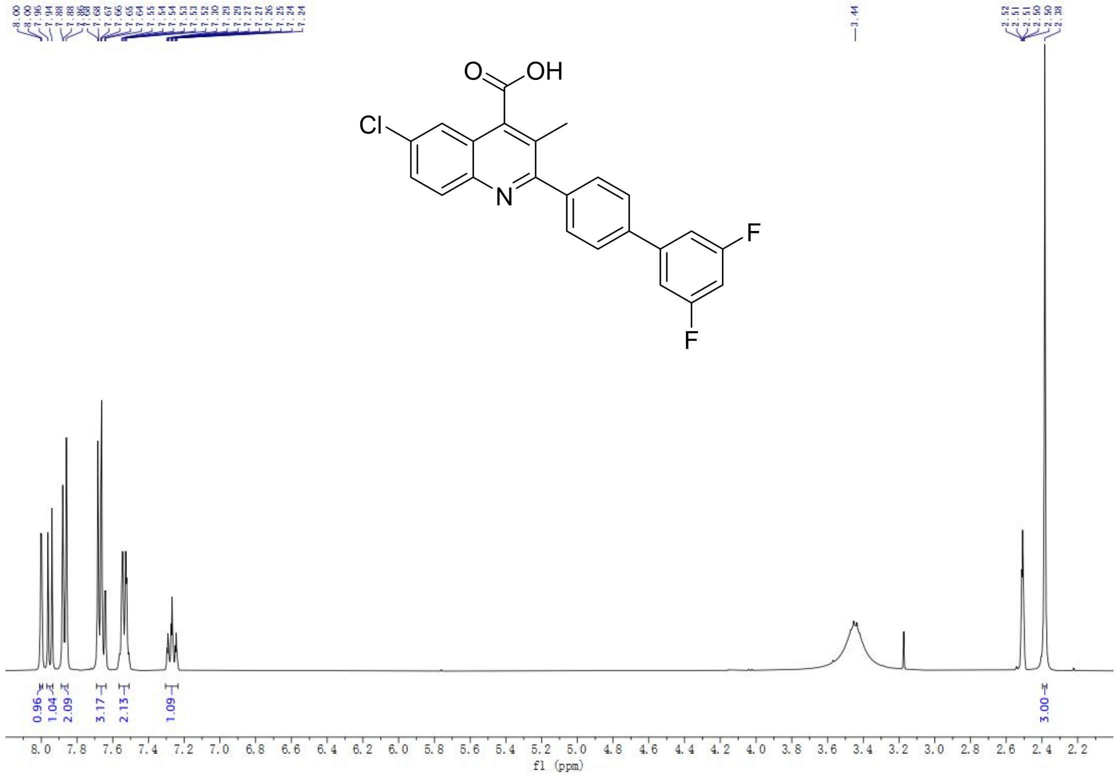

$^{1}$H-NMR spectrum of EA2

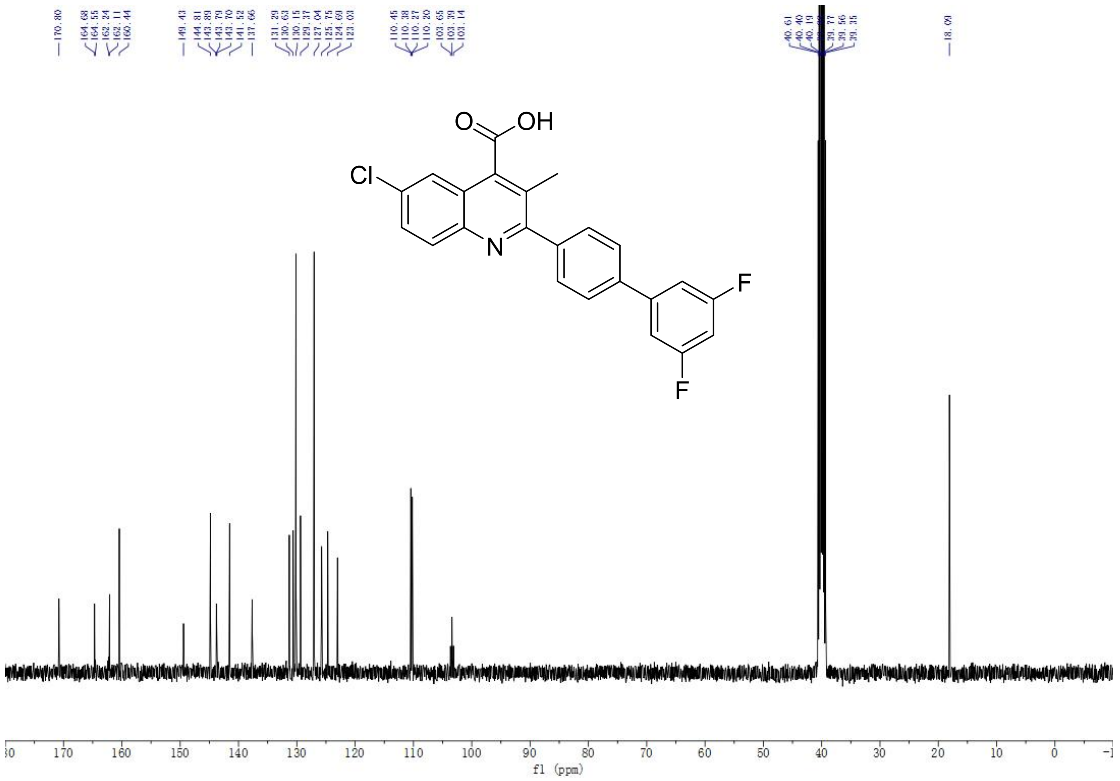

$^{13}$C-NMR spectrum of EA2

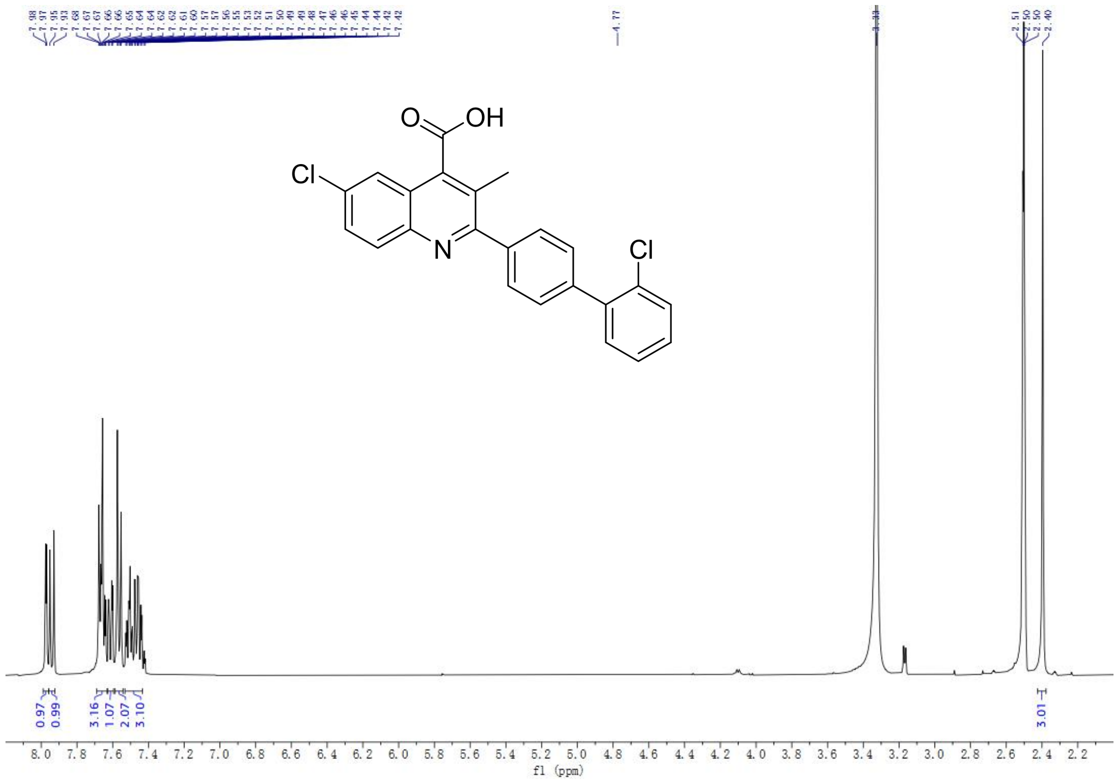

$^{1}$H-NMR spectrum of EA3

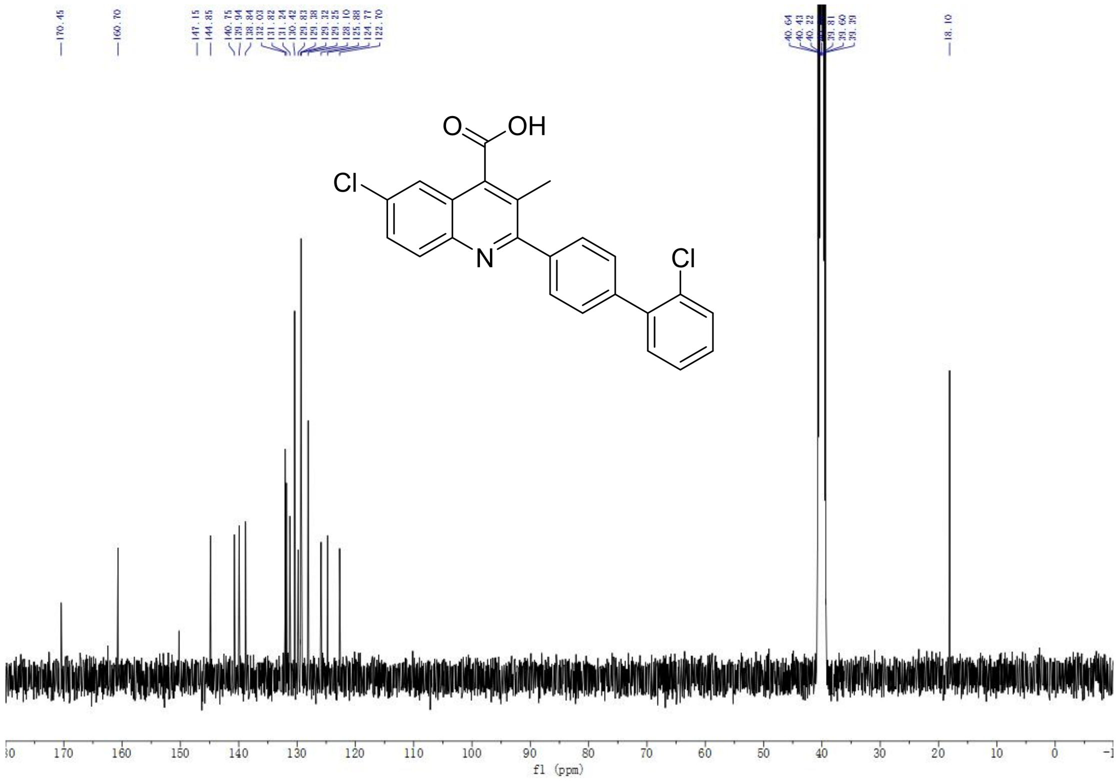

$^{13}$C-NMR spectrum of EA3

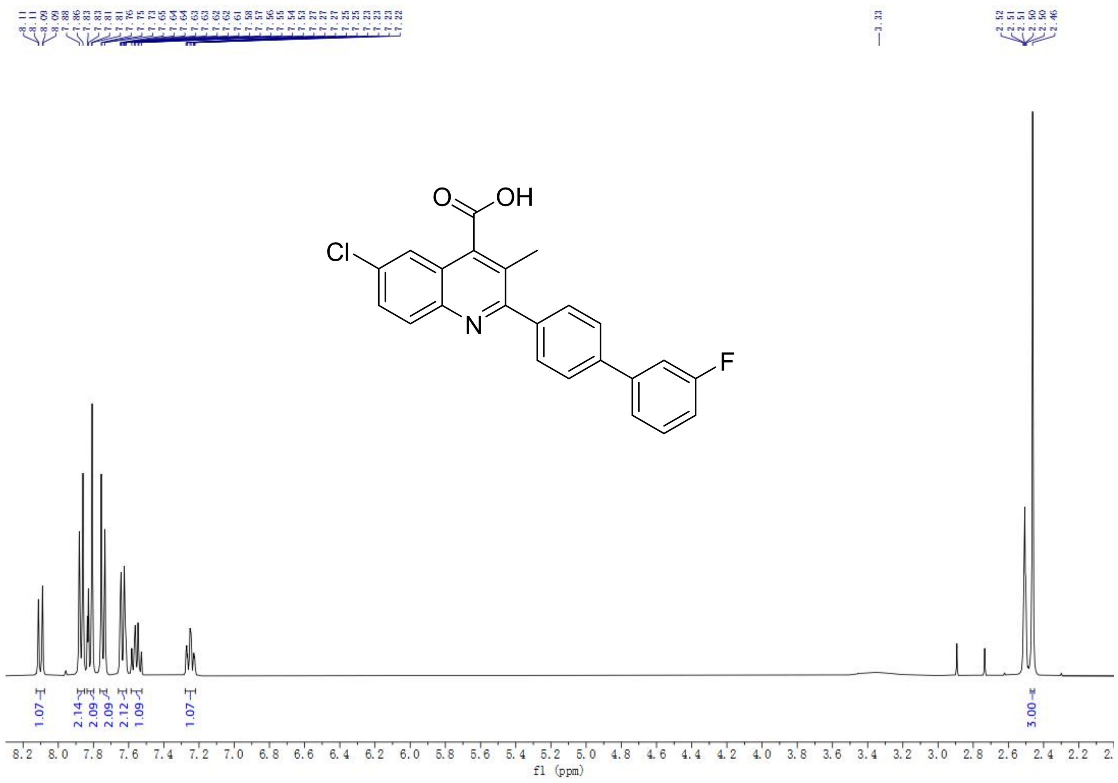

$^{1}$H-NMR spectrum of EA4

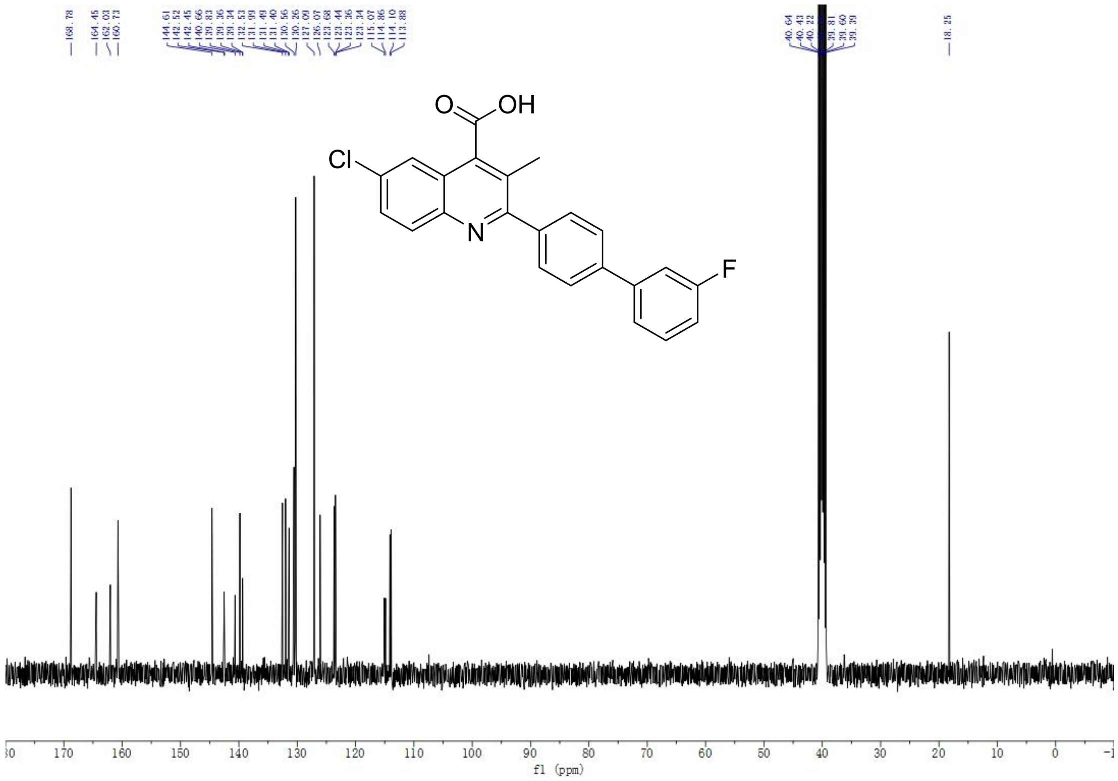

$^{13}$C-NMR spectrum of EA4

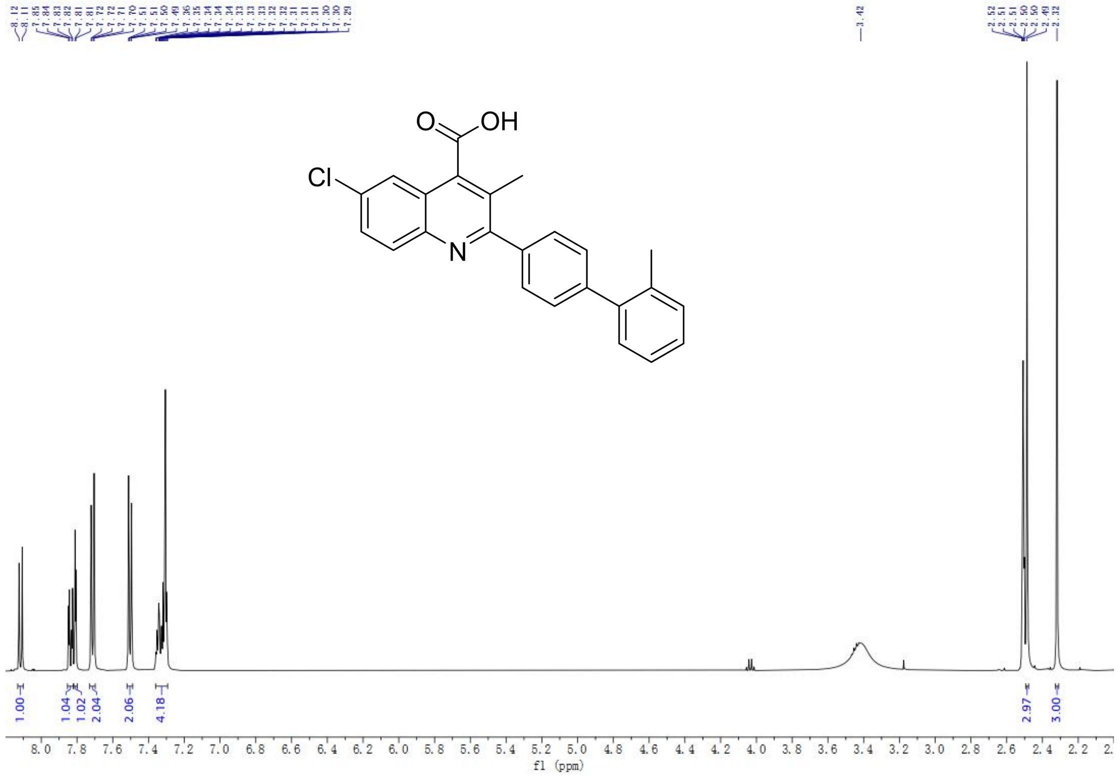

[1]H-NMR spectrum of EA6

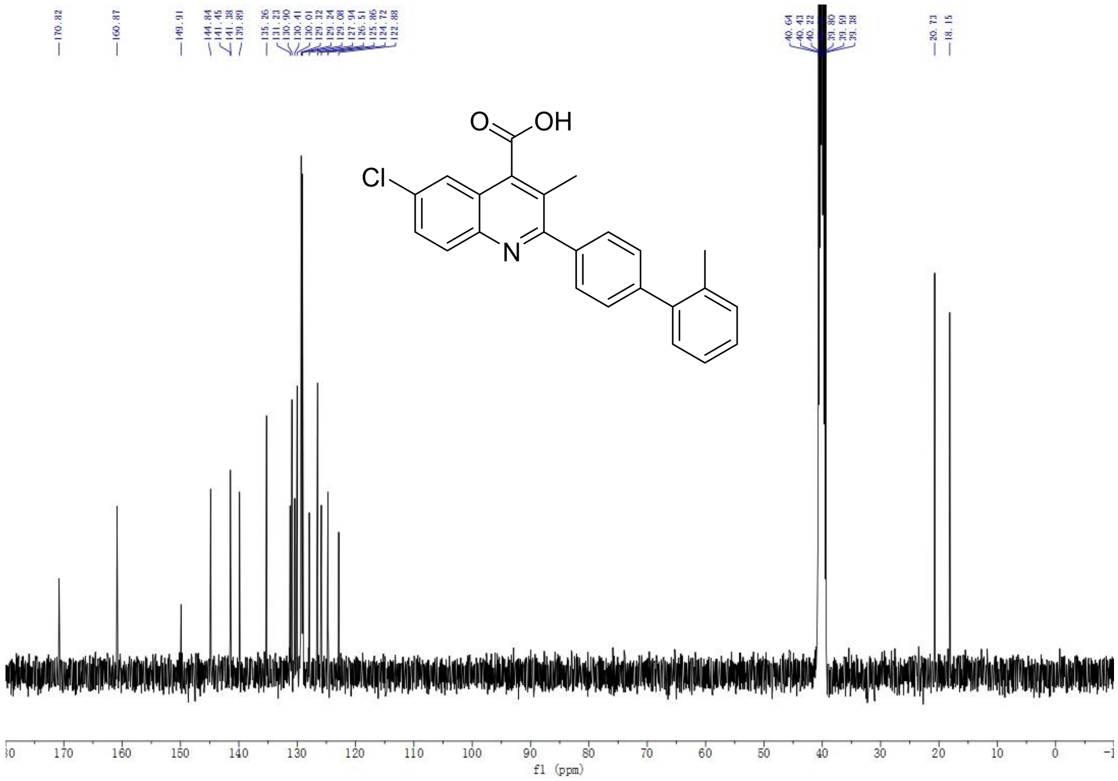

[13]C-NMR spectrum of EA6

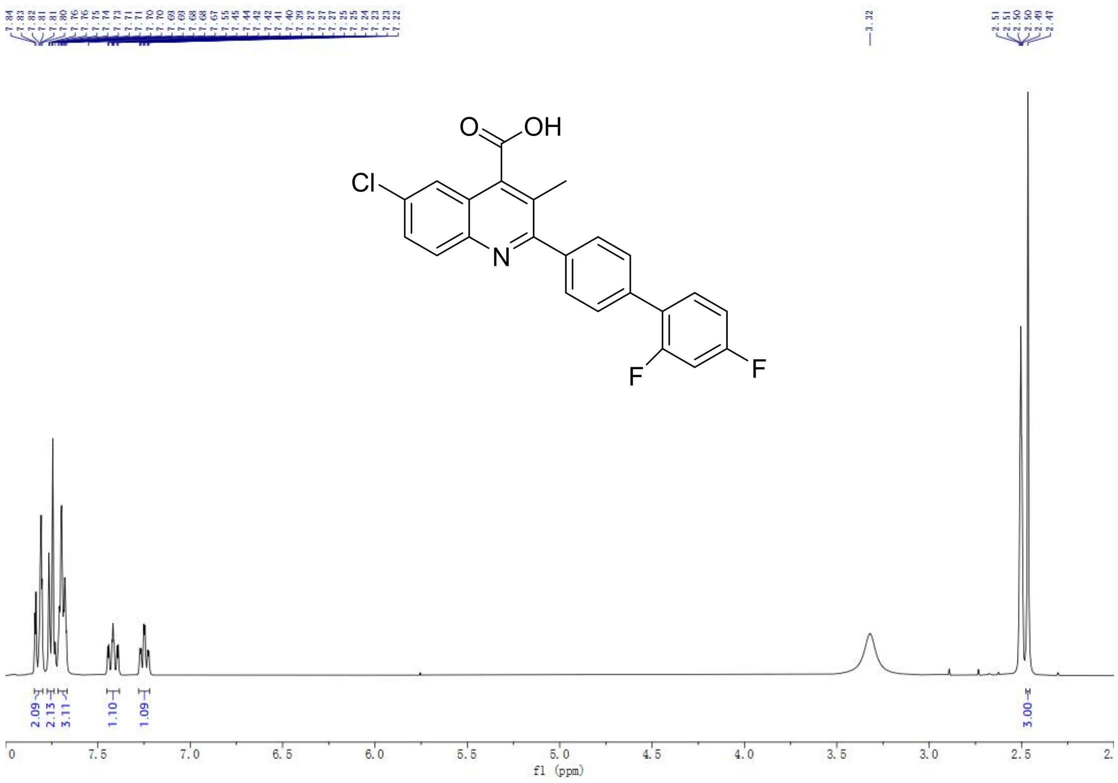

$^{1}$H-NMR spectrum of EA8

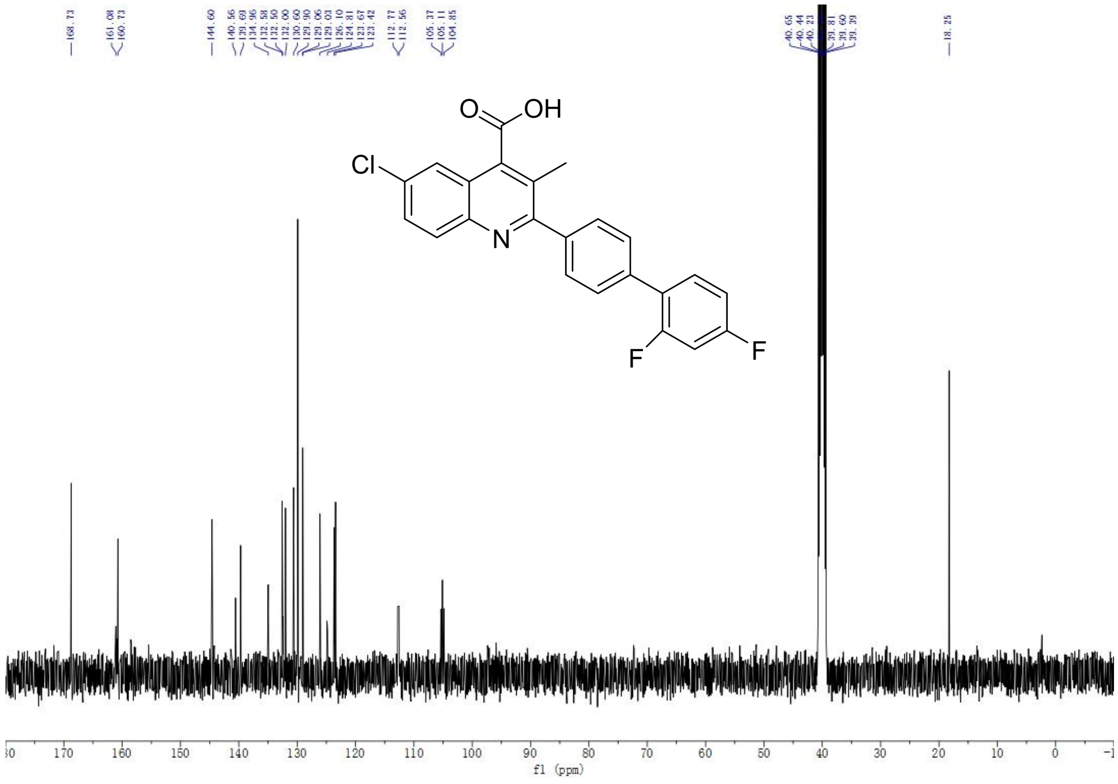

$^{13}$C-NMR spectrum of EA8

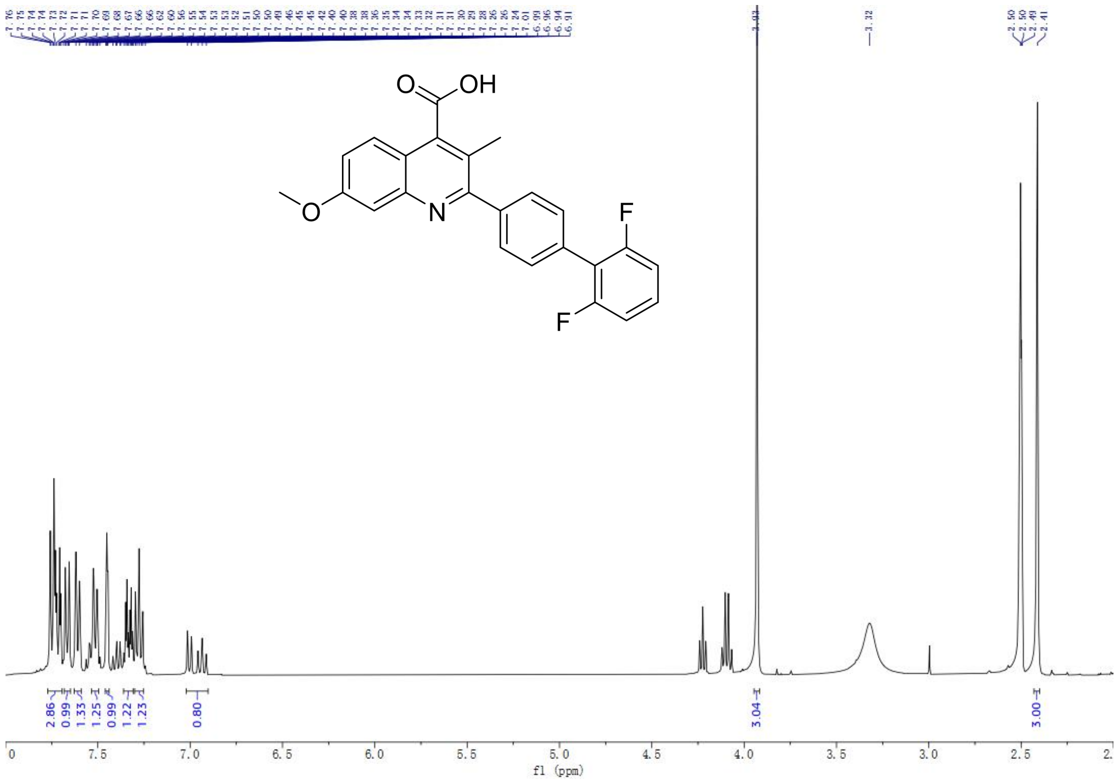

[1]H-NMR spectrum of FA11

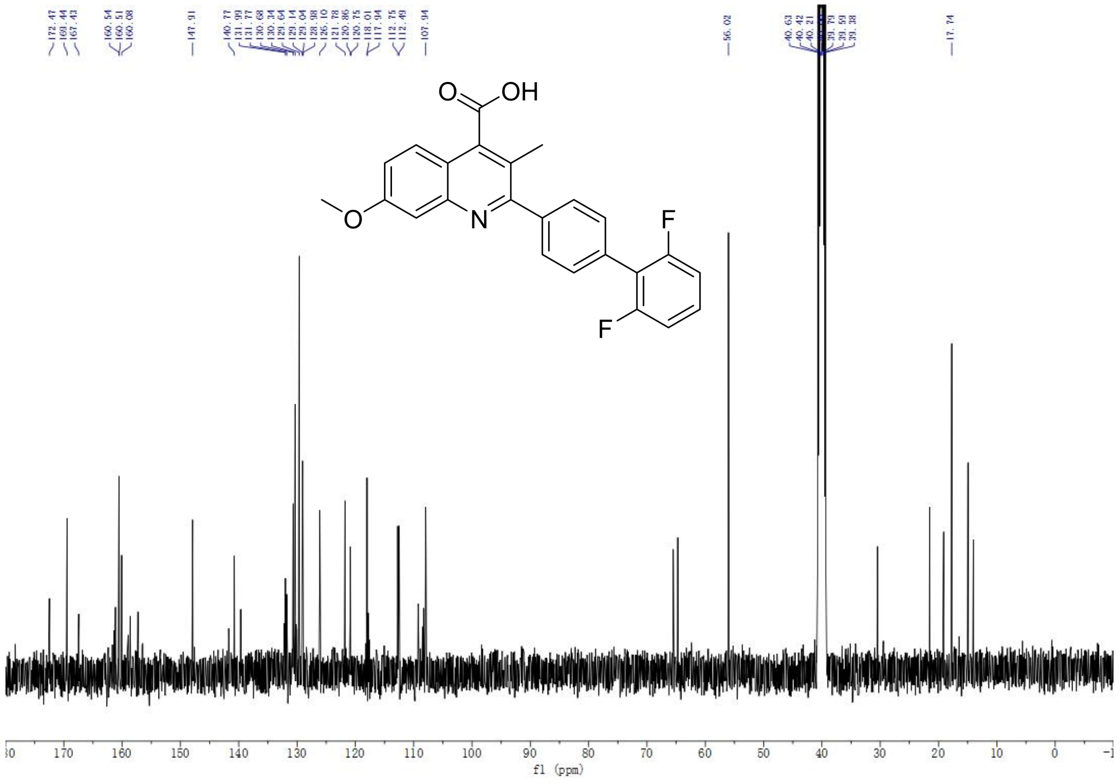

[13]C-NMR spectrum of FA11

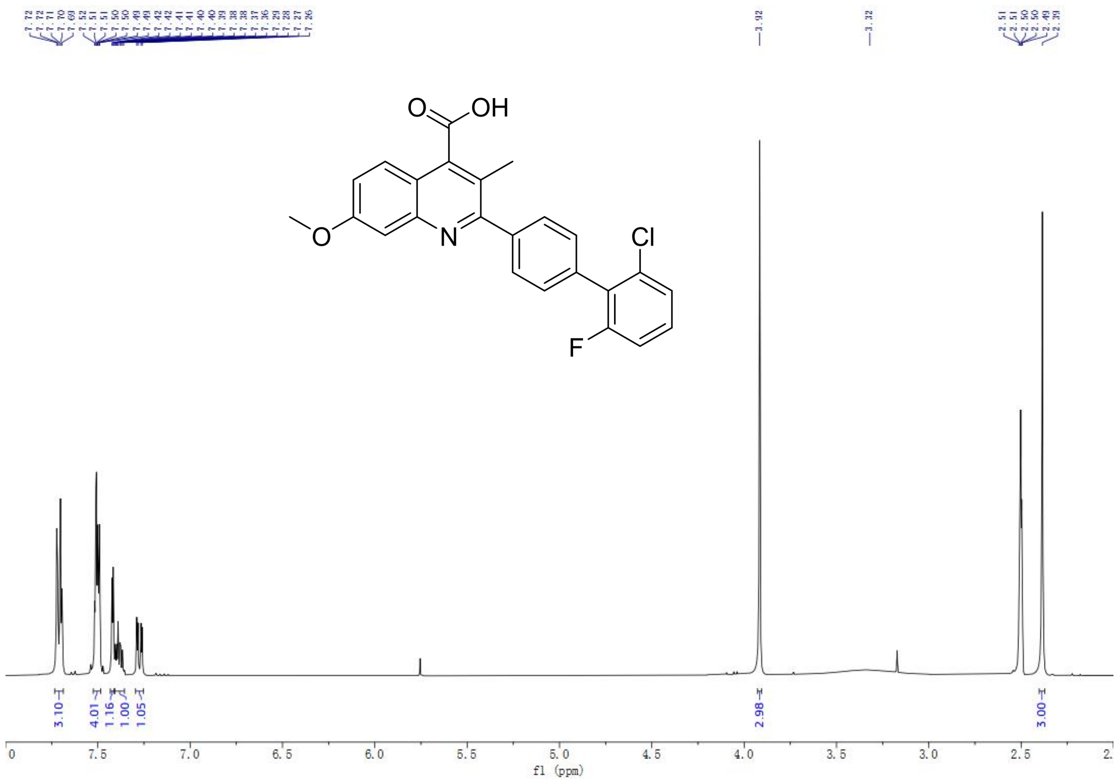

[1]H-NMR spectrum of FA12

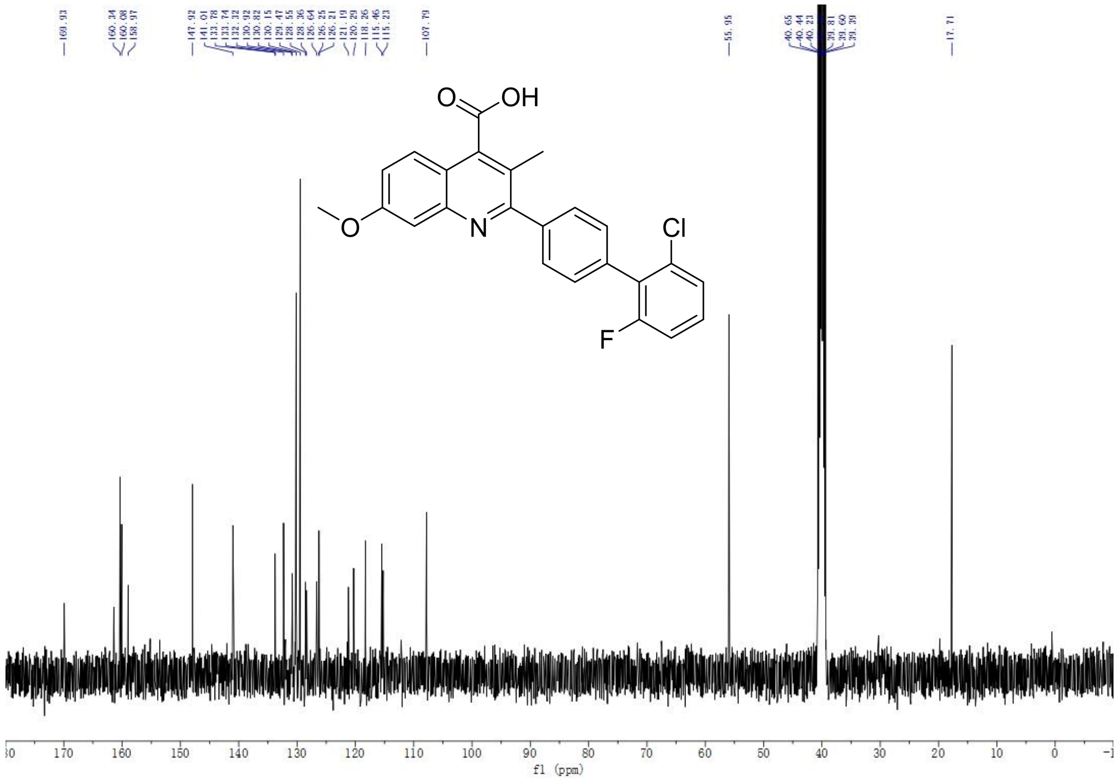

[13]C-NMR spectrum of FA12

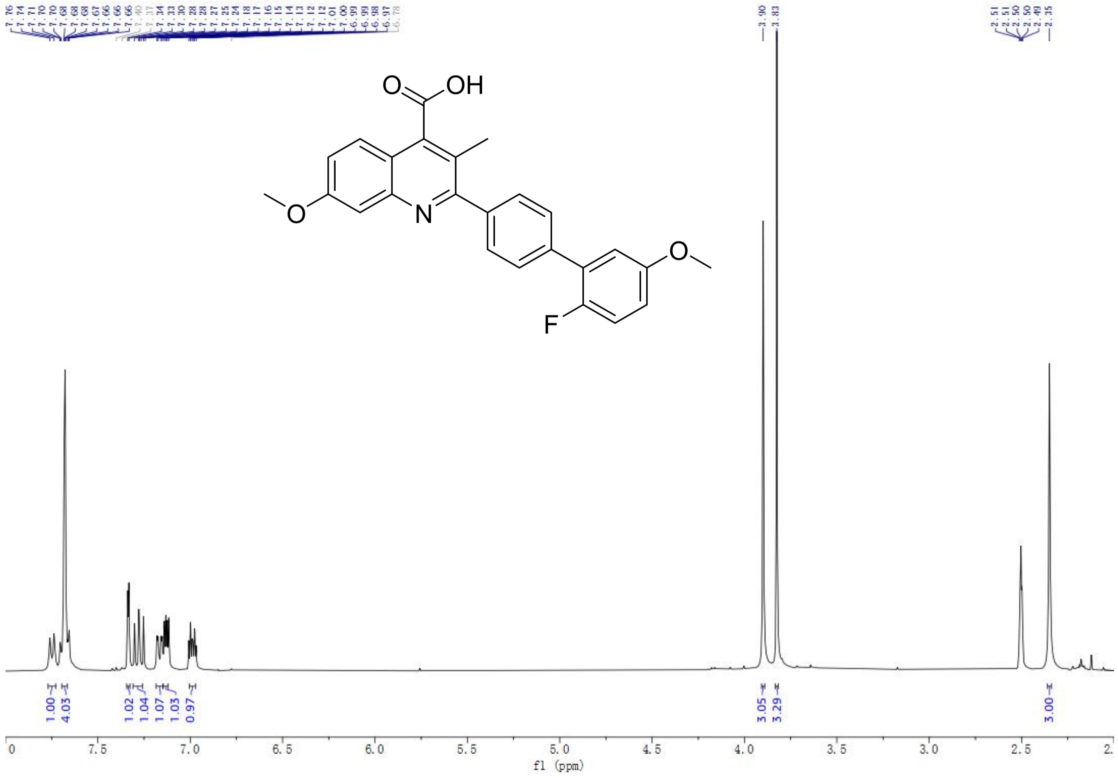

[1]H-NMR spectrum of FA14

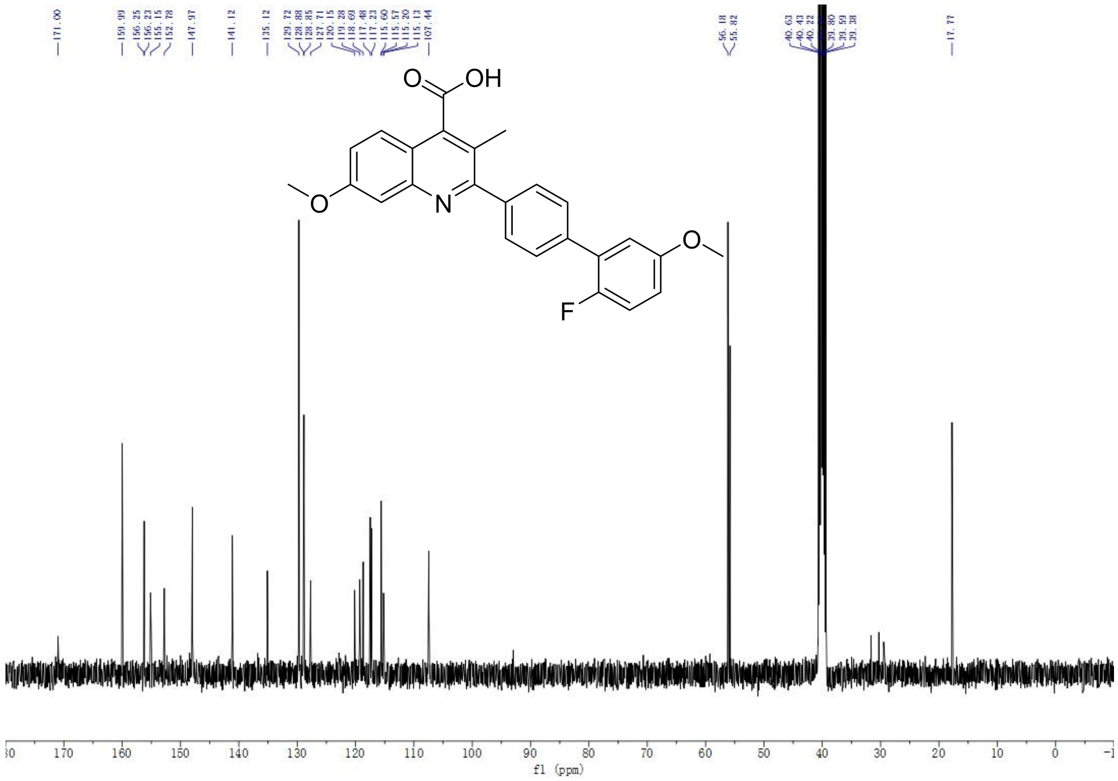

[13]C-NMR spectrum of FA14

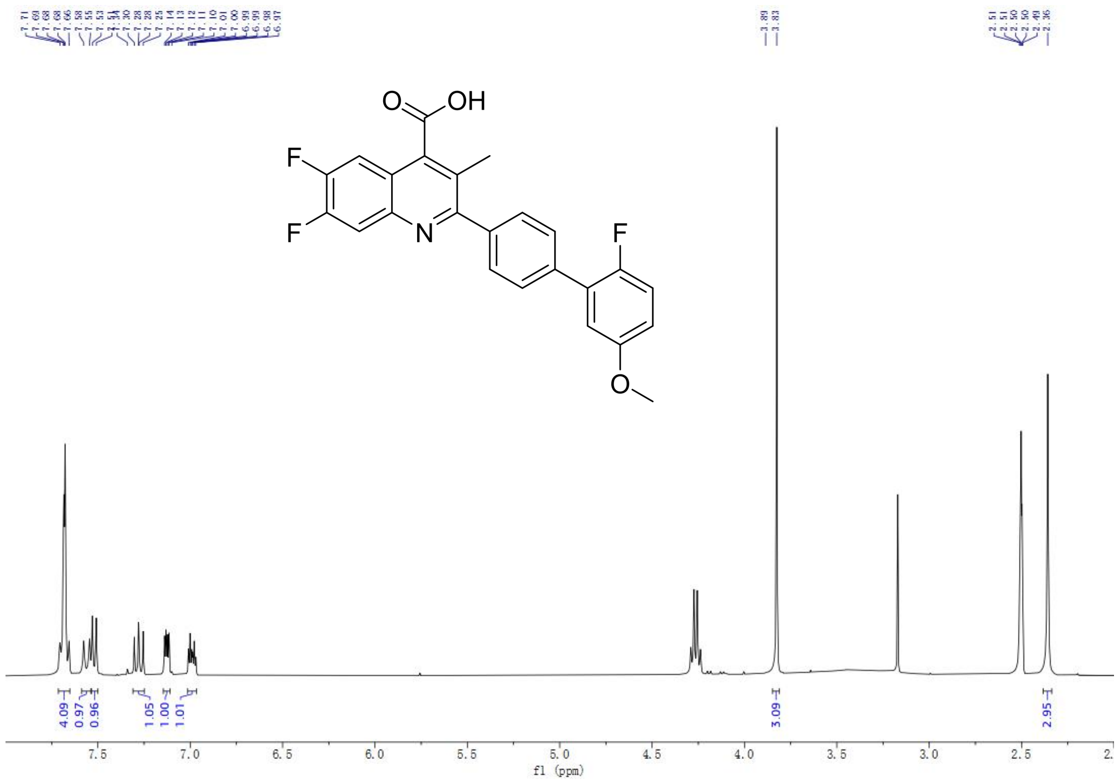

$^{1}$H-NMR spectrum of GA14

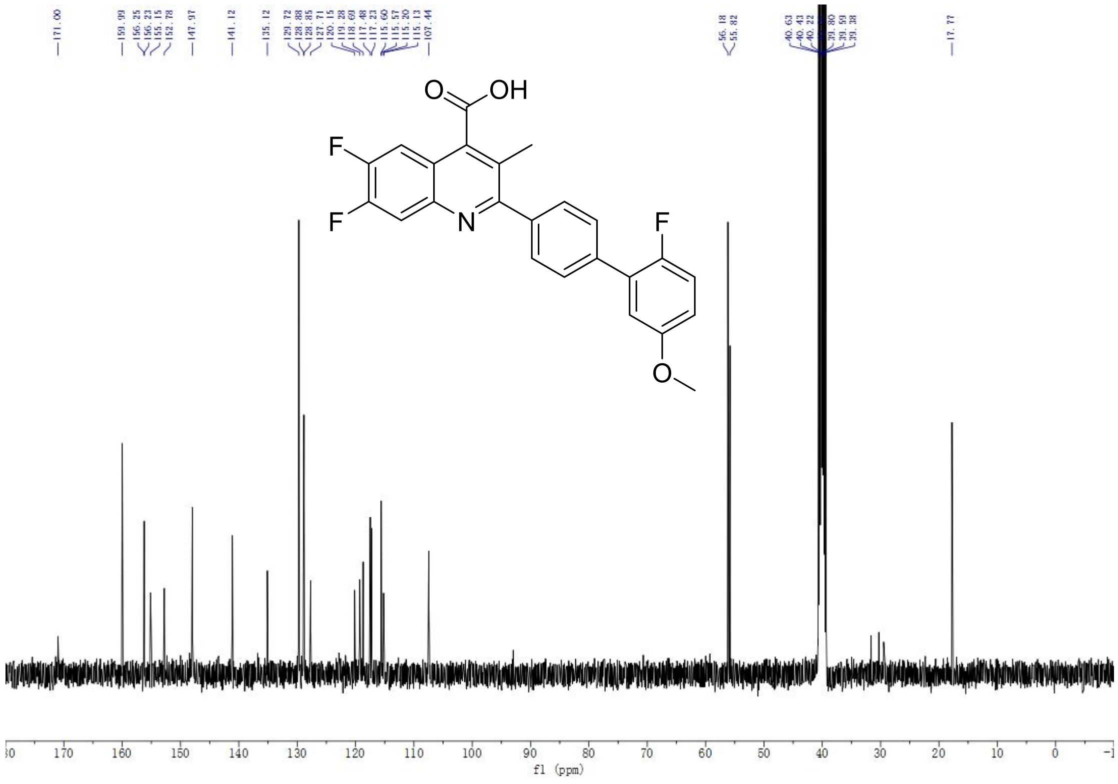

$^{13}$C-NMR spectrum of GA14

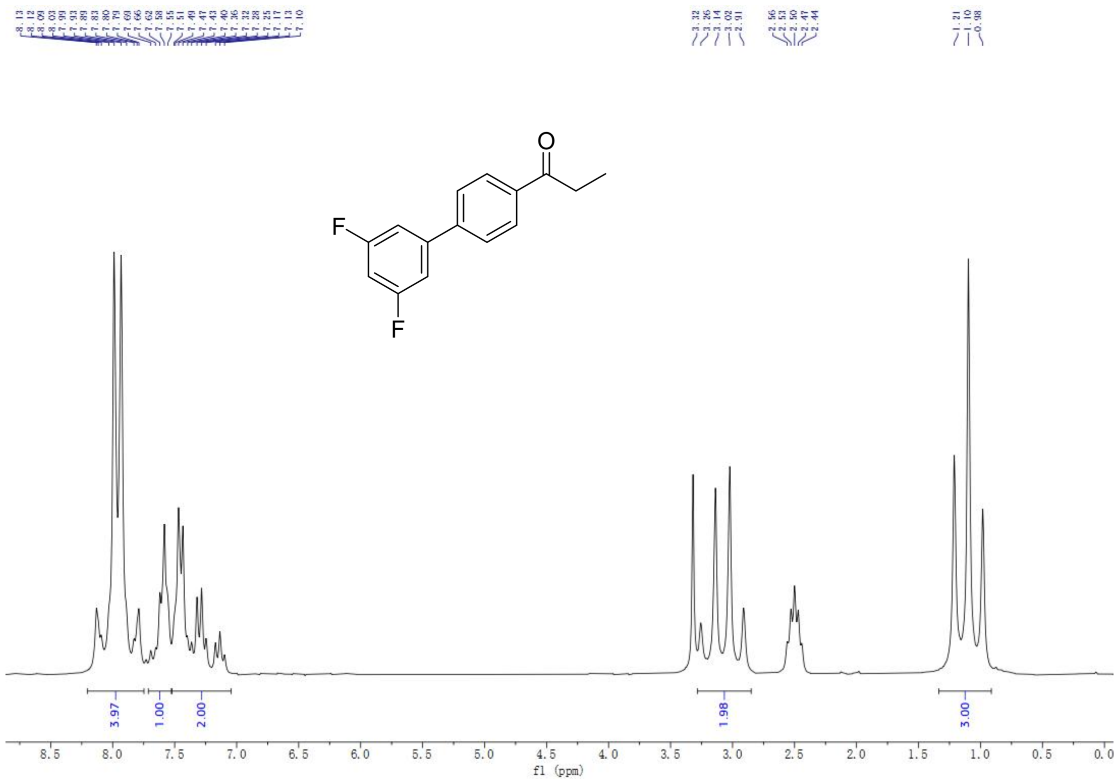

$^{1}$H-NMR spectrum of A2

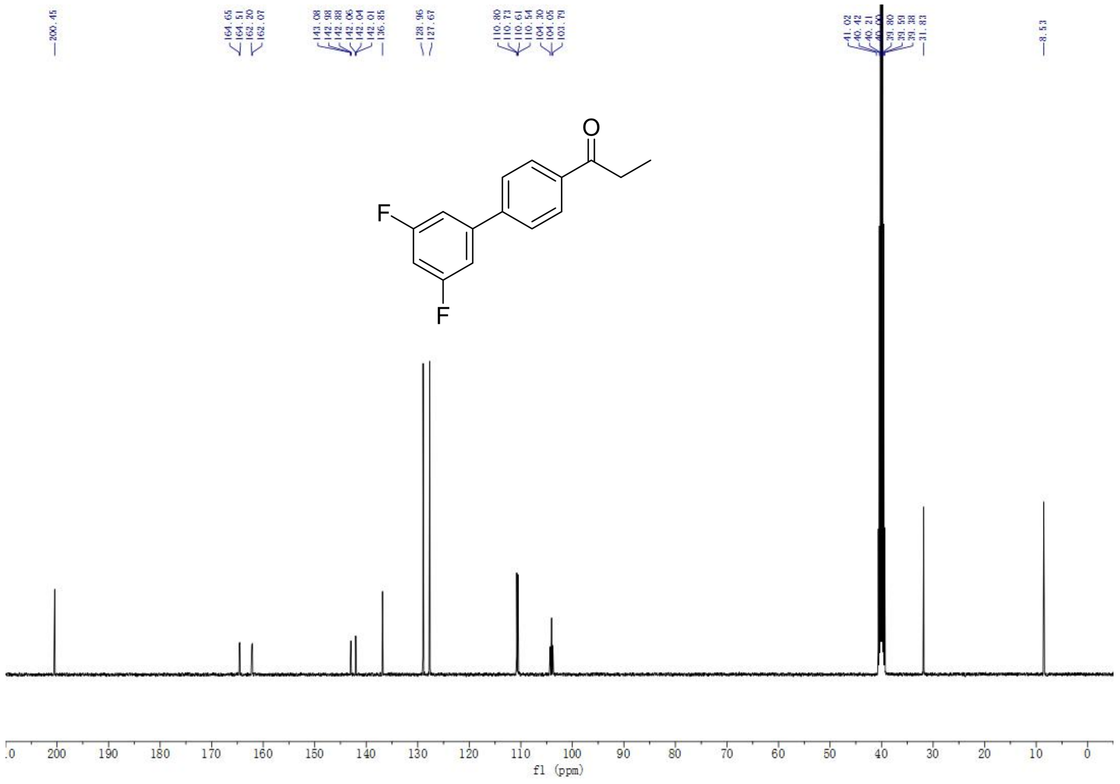

$^{13}$C-NMR spectrum of A2

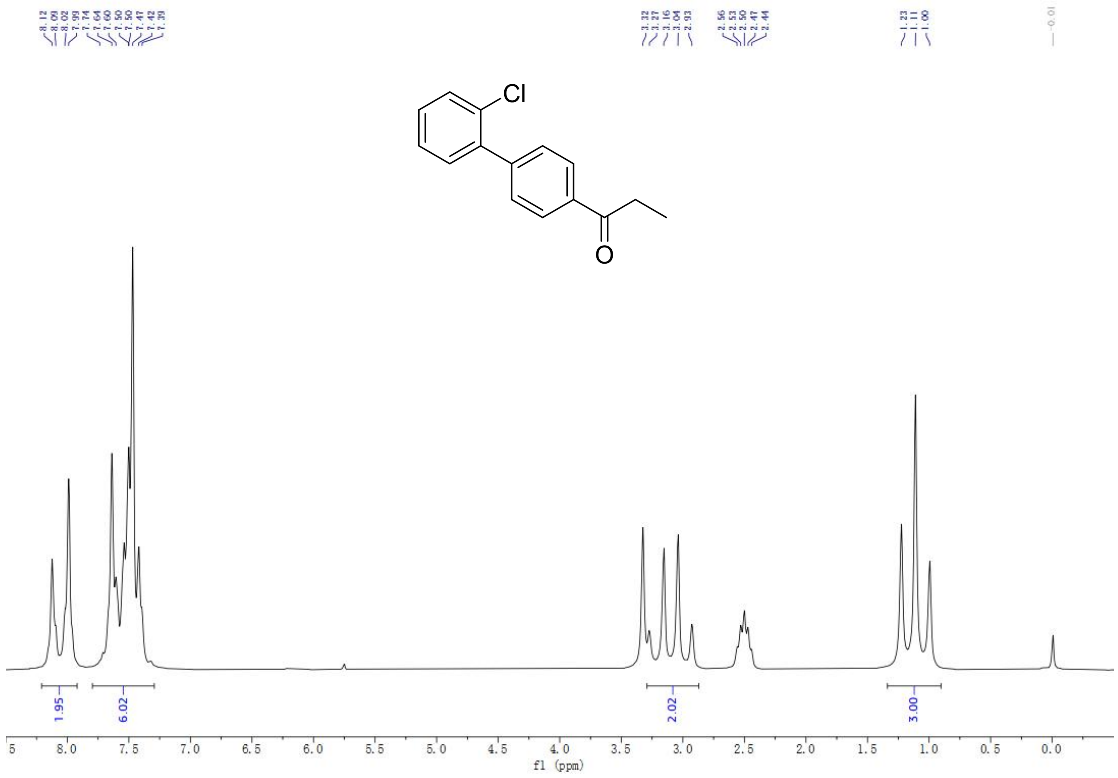

$^{1}$H-NMR spectrum of A3

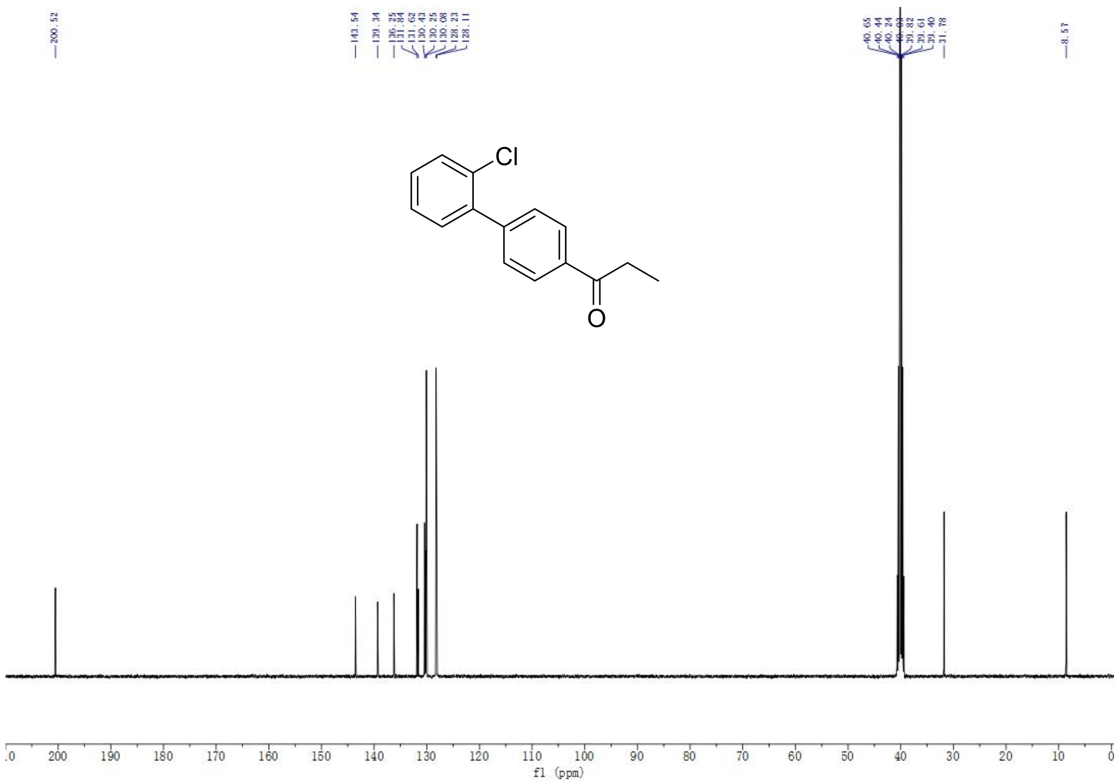

$^{13}$C-NMR spectrum of A3

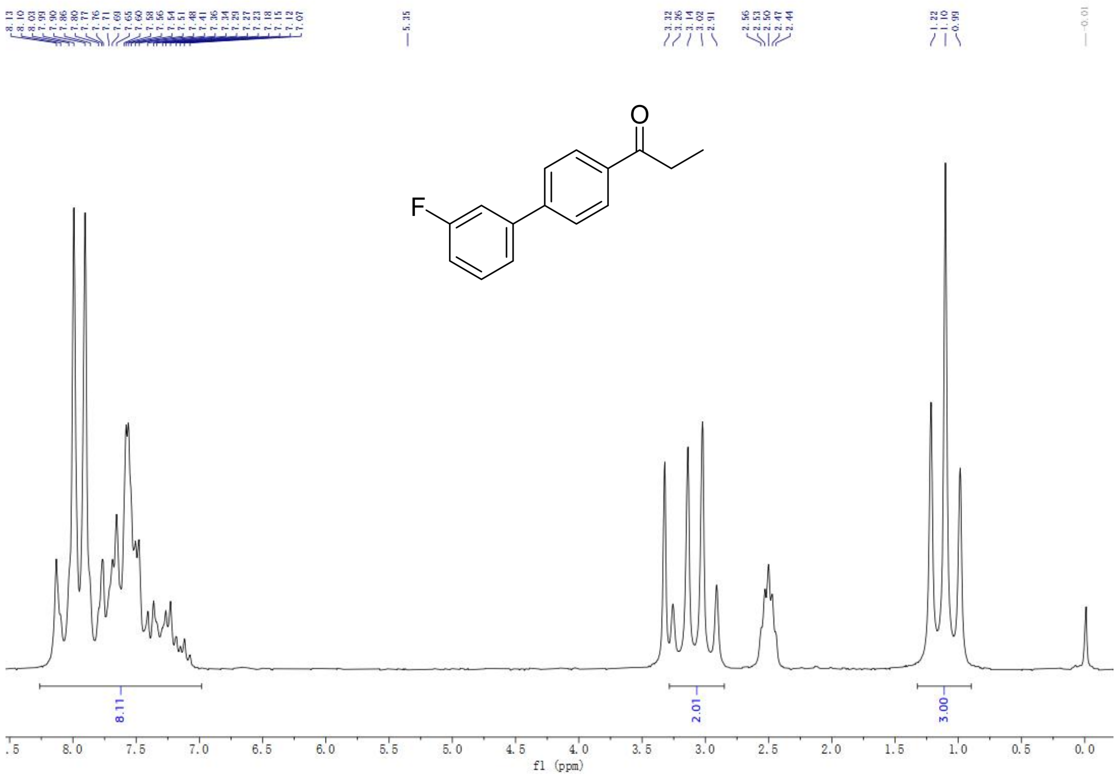

$^{1}$H-NMR spectrum of A4

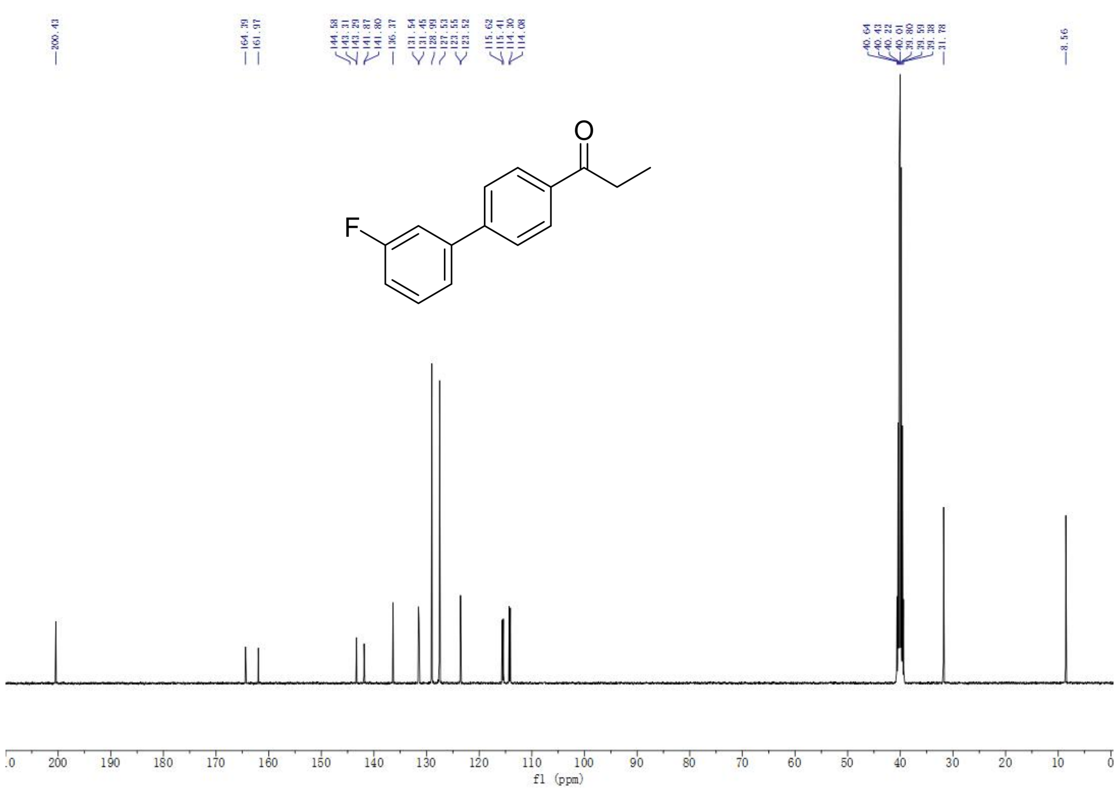

$^{13}$C-NMR spectrum of A4

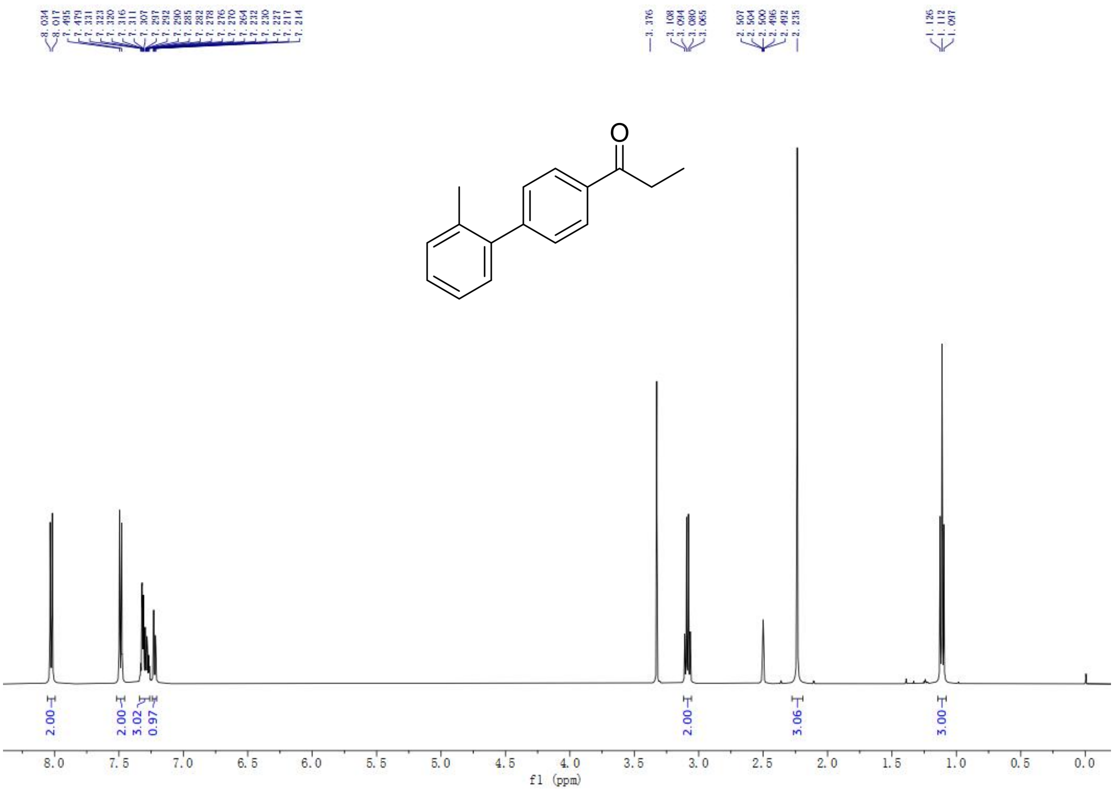

$^{1}$H-NMR spectrum of A6

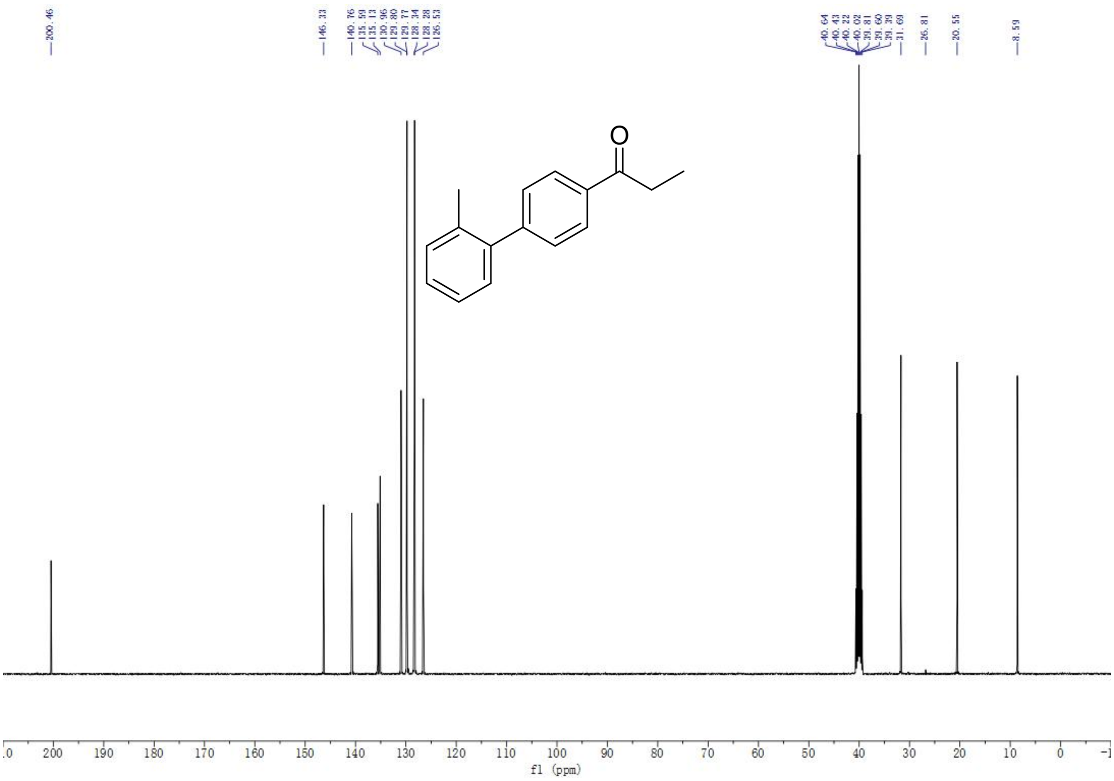

$^{13}$C-NMR spectrum of A6

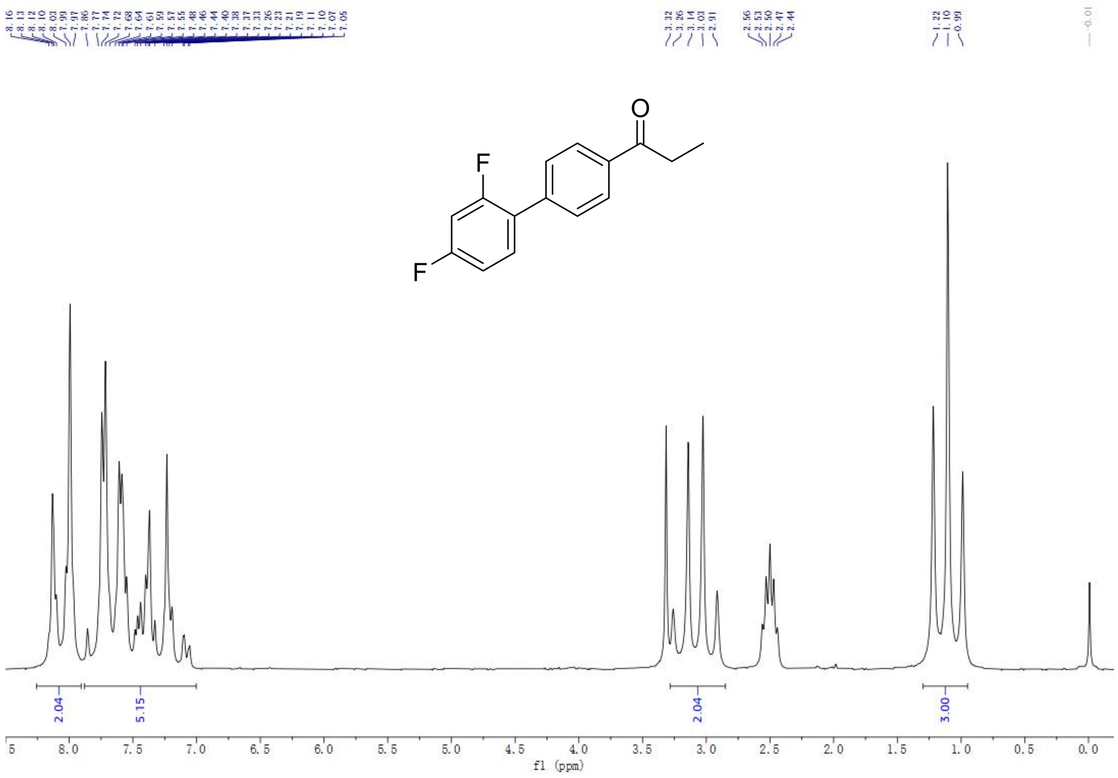

[1]H-NMR spectrum of A8

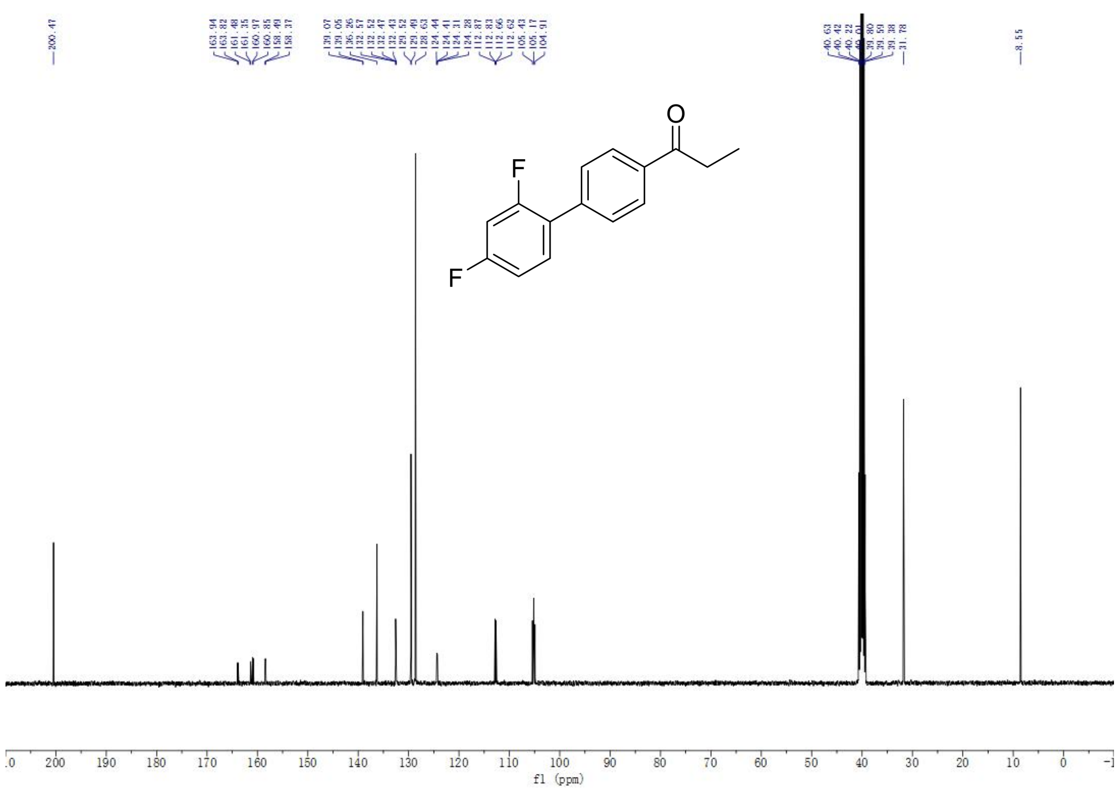

[13]C-NMR spectrum of A8

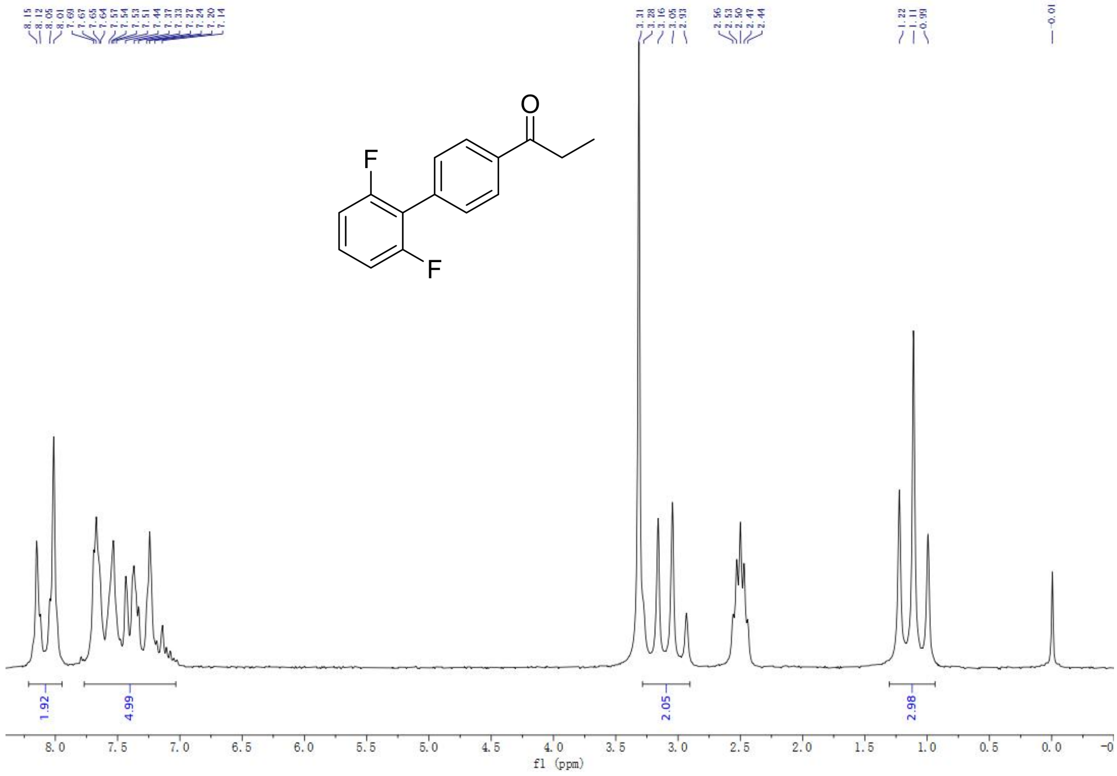

$^1$H-NMR spectrum of A11

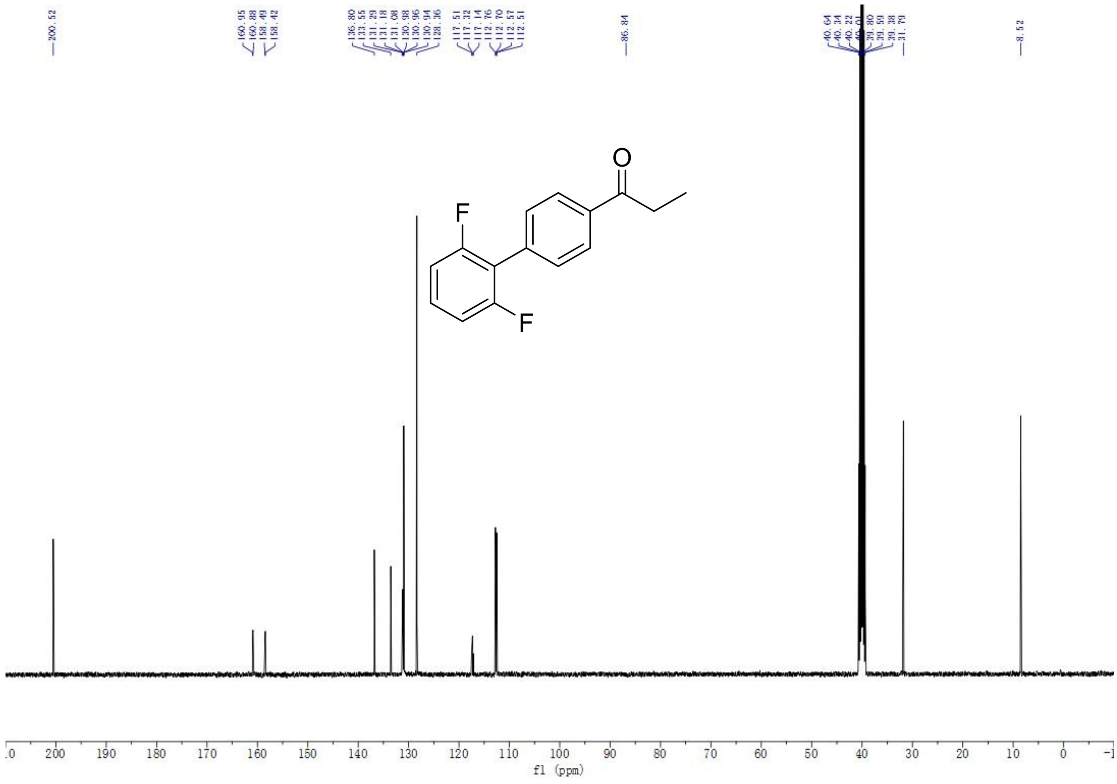

$^{13}$C-NMR spectrum of A11

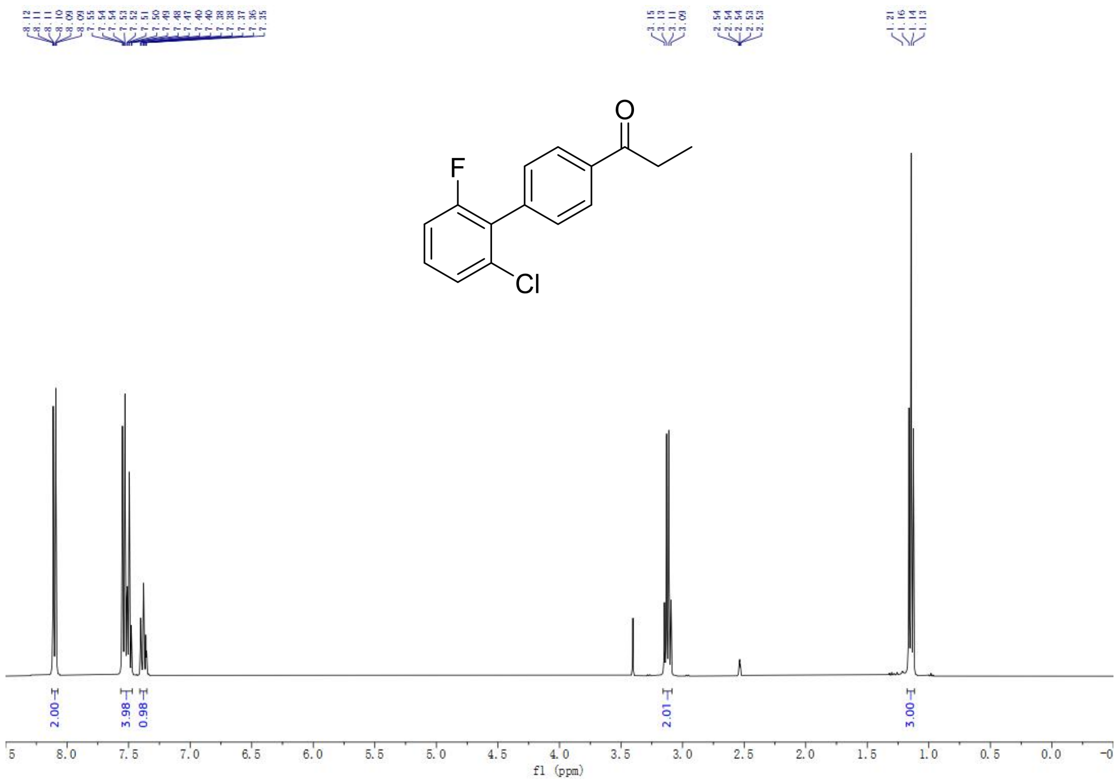

$^1$H-NMR spectrum of A12

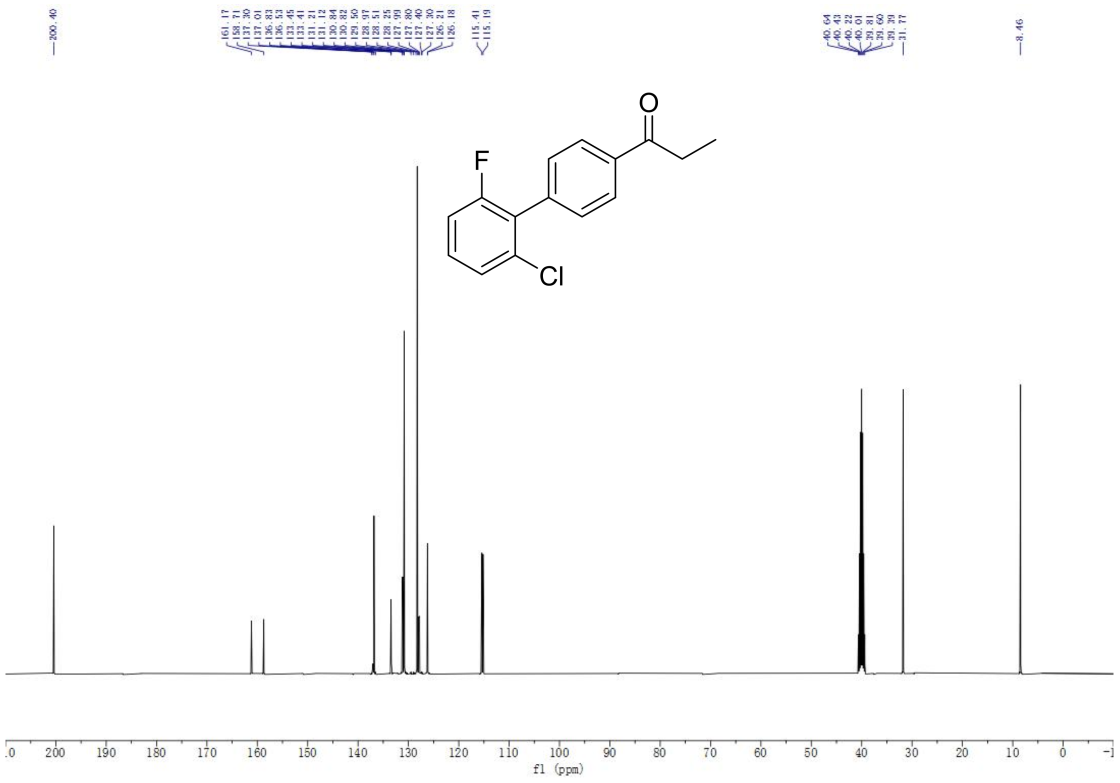

$^{13}$C-NMR spectrum of A12

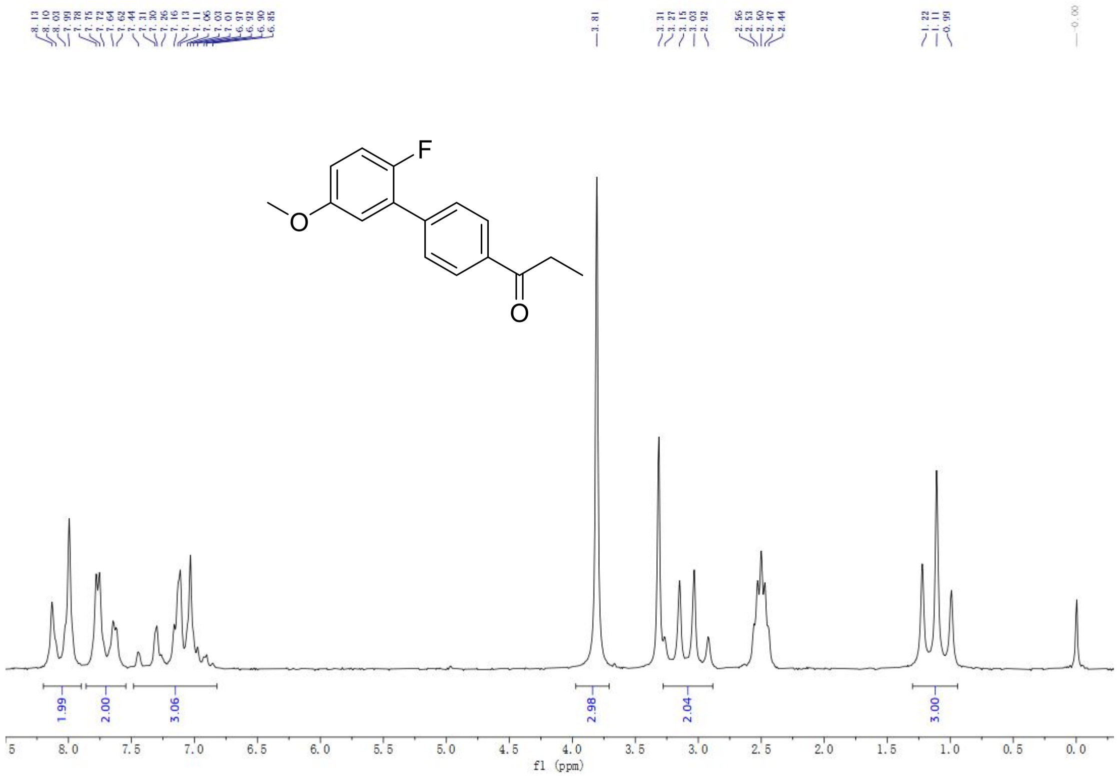

[1]H-NMR spectrum of A14

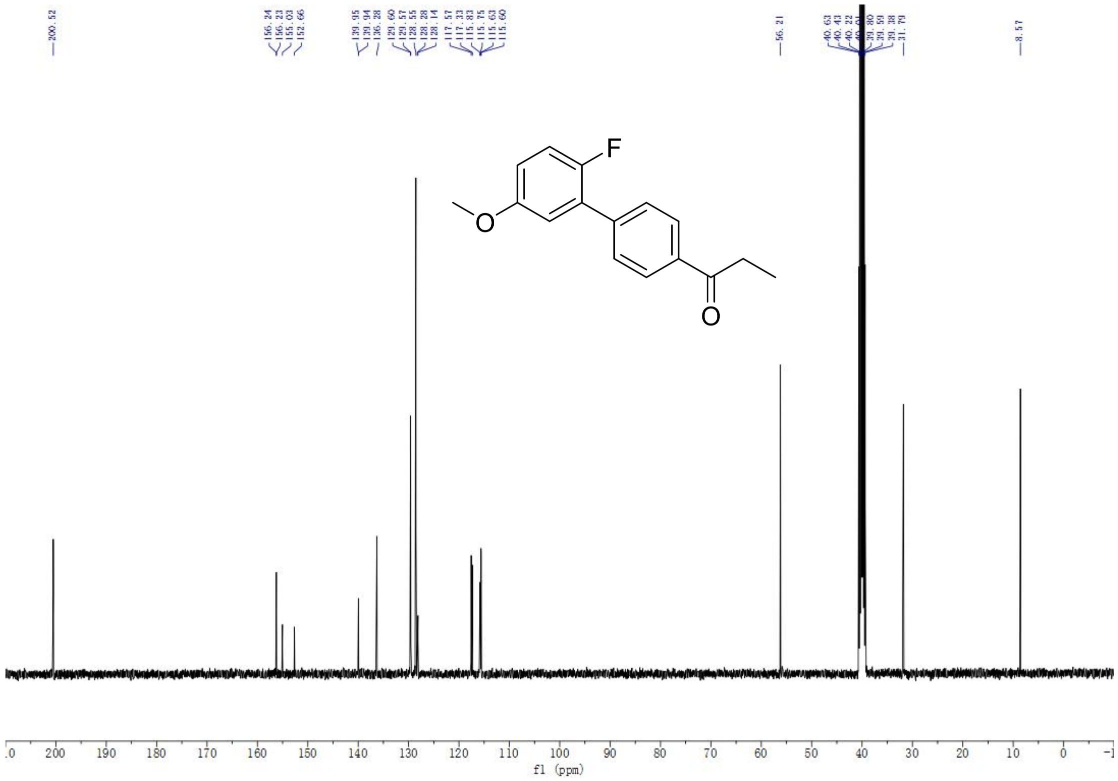

[13]C-NMR spectrum of A14

### HRMS Spectrum

Monoisotopic Mass, Even Electron Ions
1 formula(e) evaluated with 1 results within limits (up to 50 closest results for each mass)
Elements Used:
C: 23-23 H: 15-15 N: 1-1 O: 2-2 F: 3-3

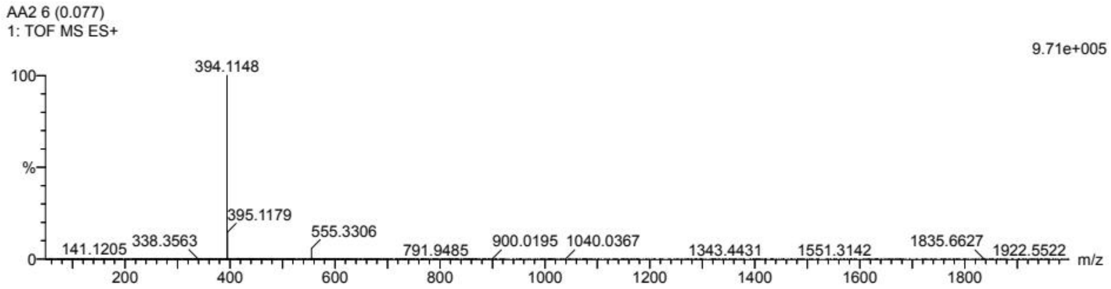

| | | | | | | | | |
|---|---|---|---|---|---|---|---|---|
| Minimum: | | | | -1.5 | | | | |
| Maximum: | | 5.0 | 50.0 | 50.0 | | | | |
| Mass | Calc. Mass | mDa | PPM | DBE | i-FIT | Norm | Conf(%) | Formula |
| 394.1148 | 394.1055 | 9.3 | 23.6 | 15.5 | 97.8 | n/a | n/a | C23 H15 N O2 F3 |

HRMS Spectrum of AA2

Monoisotopic Mass, Even Electron Ions
1 formula(e) evaluated with 1 results within limits (up to 50 closest results for each mass)
Elements Used:
C: 23-23 H: 16-16 N: 1-1 O: 2-2 F: 1-1 Cl: 1-1

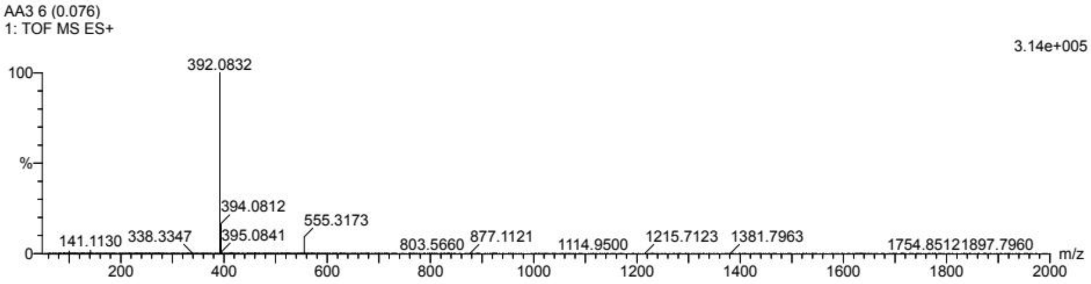

| | | | | | | | | |
|---|---|---|---|---|---|---|---|---|
| Minimum: | | | | -1.5 | | | | |
| Maximum: | | 5.0 | 50.0 | 50.0 | | | | |
| Mass | Calc. Mass | mDa | PPM | DBE | i-FIT | Norm | Conf(%) | Formula |
| 392.0832 | 392.0854 | -2.2 | -5.6 | 15.5 | 96.6 | n/a | n/a | C23 H16 N O2 F Cl |

HRMS Spectrum of AA3

Monoisotopic Mass, Even Electron Ions
1 formula(e) evaluated with 1 results within limits (up to 50 closest results for each mass)
Elements Used:
C: 24-24 H: 19-19 N: 1-1 O: 3-3 Cl: 1-1

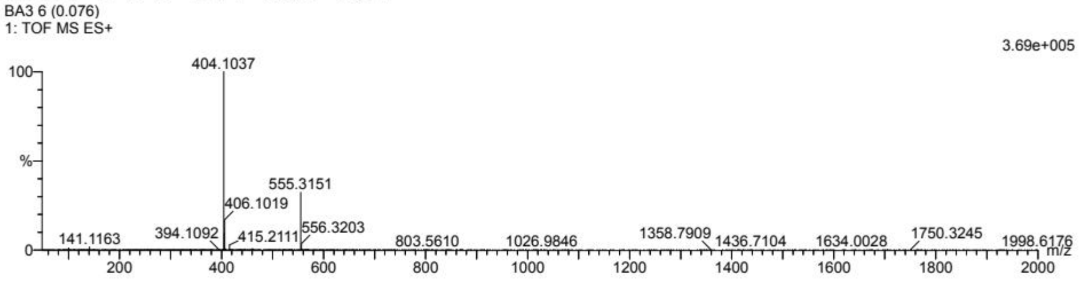

| | | | | | | | | |
|---|---|---|---|---|---|---|---|---|
| Minimum: | | | | -1.5 | | | | |
| Maximum: | | 5.0 | 50.0 | 50.0 | | | | |
| Mass | Calc. Mass | mDa | PPM | DBE | i-FIT | Norm | Conf(%) | Formula |
| 404.1037 | 404.1053 | -1.6 | -4.0 | 15.5 | 89.0 | n/a | n/a | C24 H19 N O3 Cl |

#### HRMS Spectrum of BA3

Monoisotopic Mass, Even Electron Ions
1 formula(e) evaluated with 1 results within limits (up to 50 closest results for each mass)
Elements Used:
C: 24-24 H: 19-19 N: 1-1 O: 3-3 F: 1-1

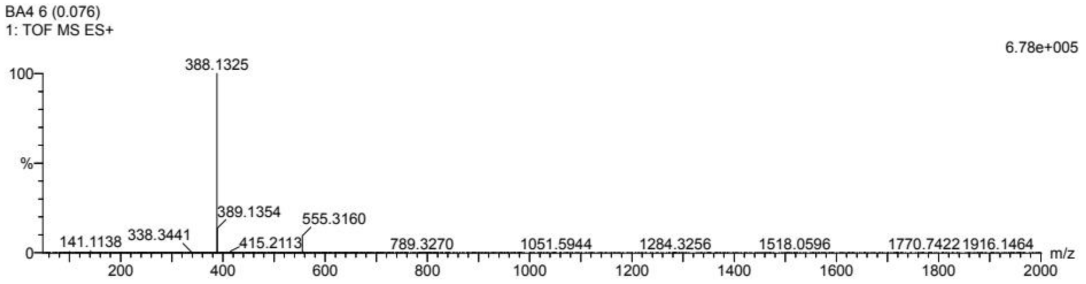

| | | | | |
|---|---|---|---|---|
| Minimum: | | | | -1.5 |
| Maximum: | | 5.0 | 50.0 | 50.0 |

| Mass | Calc. Mass | mDa | PPM | DBE | i-FIT | Norm | Conf(%) | Formula |
|---|---|---|---|---|---|---|---|---|
| 388.1325 | 388.1349 | -2.4 | -6.2 | 15.5 | 166.1 | n/a | n/a | C24 H19 N O3 F |

#### HRMS Spectrum of BA4

Monoisotopic Mass, Even Electron Ions
1 formula(e) evaluated with 1 results within limits (up to 50 closest results for each mass)
Elements Used:
C: 24-24 H: 18-18 N: 1-1 O: 2-2 F: 2-2

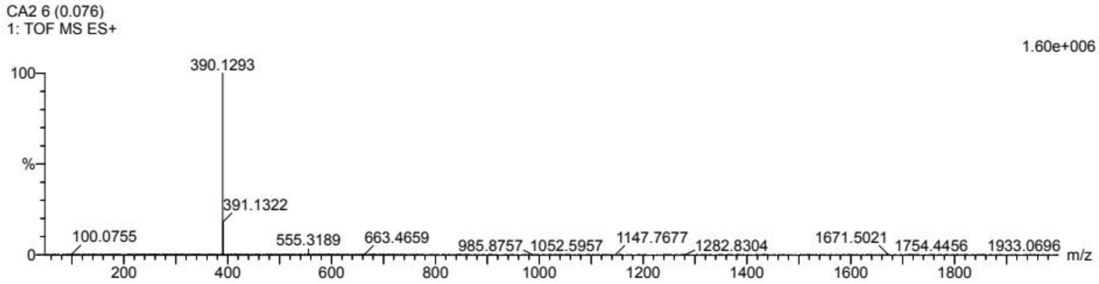

| | | | | |
|---|---|---|---|---|
| Minimum: | | | | -1.5 |
| Maximum: | | 5.0 | 50.0 | 50.0 |

| Mass | Calc. Mass | mDa | PPM | DBE | i-FIT | Norm | Conf(%) | Formula |
|---|---|---|---|---|---|---|---|---|
| 390.1293 | 390.1306 | -1.3 | -3.3 | 15.5 | 153.0 | n/a | n/a | C24 H18 N O2 F2 |

#### HRMS Spectrum of CA2

Monoisotopic Mass, Even Electron Ions
1 formula(e) evaluated with 1 results within limits (up to 50 closest results for each mass)
Elements Used:
C: 24-24 H: 19-19 N: 1-1 O: 2-2 F: 1-1

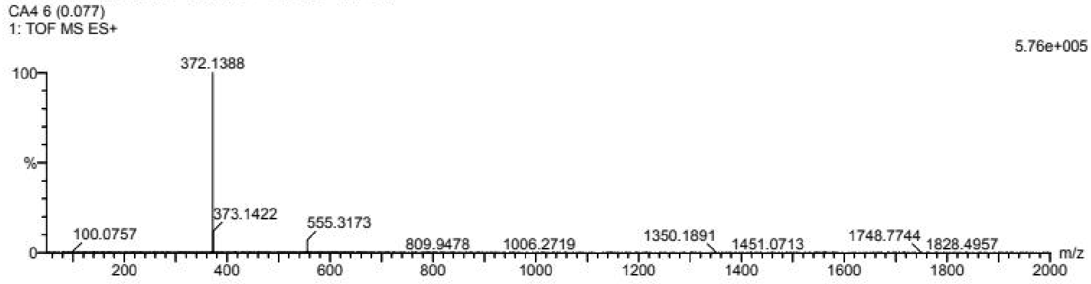

| | | | | |
|---|---|---|---|---|
| Minimum: | | | | -1.5 |
| Maximum: | | 5.0 | 50.0 | 50.0 |

| Mass | Calc. Mass | mDa | PPM | DBE | i-FIT | Norm | Conf(%) | Formula |
|---|---|---|---|---|---|---|---|---|
| 372.1388 | 372.1400 | -1.2 | -3.2 | 15.5 | 124.5 | n/a | n/a | C24 H19 N O2 F |

#### HRMS Spectrum of CA4

Monoisotopic Mass, Even Electron Ions
1 formula(e) evaluated with 1 results within limits (up to 50 closest results for each mass)
Elements Used:
C: 24-24 H: 18-18 N: 1-1 O: 2-2 F: 2-2
CA8 6 (0.076)
1: TOF MS ES+

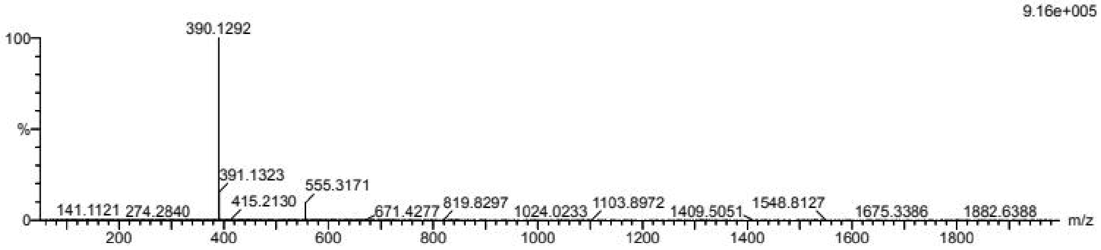

| | | | | | | | | |
|---|---|---|---|---|---|---|---|---|
| Minimum: | | | | -1.5 | | | | |
| Maximum: | | 5.0 | 50.0 | 50.0 | | | | |
| Mass | Calc. Mass | mDa | PPM | DBE | i-FIT | Norm | Conf(%) | Formula |
| 390.1292 | 390.1306 | -1.4 | -3.6 | 15.5 | 145.8 | n/a | n/a | C24 H18 N O2 F2 |

HRMS Spectrum of CA8

Monoisotopic Mass, Even Electron Ions
7 formula(e) evaluated with 1 results within limits (up to 50 best isotopic matches for each mass)
Elements Used:
C: 23-23 H: 13-13 N: 1-1 O: 2-2 F: 2-2 S: 0-6 Cl: 1-1
EA2-0914 12 (0.088)

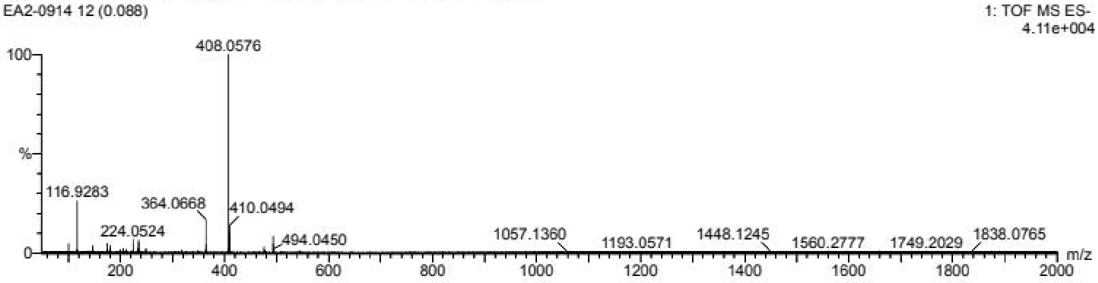

| | | | | | | | | |
|---|---|---|---|---|---|---|---|---|
| Minimum: | | | | -1.5 | | | | |
| Maximum: | | 5.0 | 10.0 | 50.0 | | | | |
| Mass | Calc. Mass | mDa | PPM | DBE | i-FIT | Norm | Conf(%) | Formula |
| 408.0576 | 408.0603 | -2.7 | -6.6 | 16.5 | 377.2 | n/a | n/a | C23 H13 N O2 F2 Cl |

HRMS Spectrum of EA2

Monoisotopic Mass, Even Electron Ions
1 formula(e) evaluated with 1 results within limits (up to 50 best isotopic matches for each mass)
Elements Used:
C: 23-23 H: 14-14 N: 1-1 O: 2-2 Cl: 2-2
EA3-0914 13 (0.093)

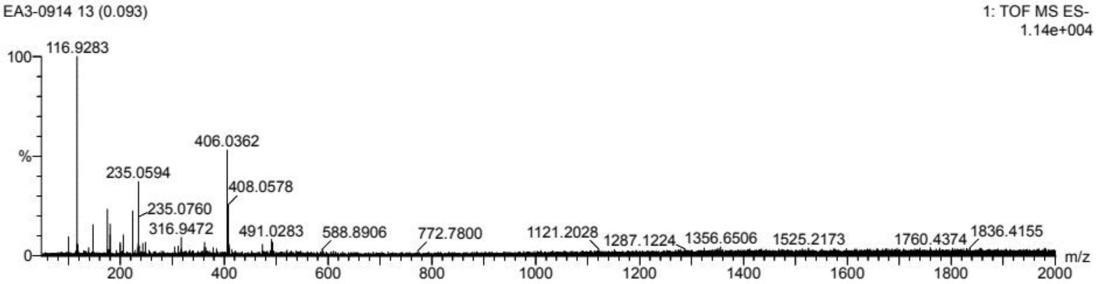

| | | | | | | | | |
|---|---|---|---|---|---|---|---|---|
| Minimum: | | | | -1.5 | | | | |
| Maximum: | | 5.0 | 10.0 | 50.0 | | | | |
| Mass | Calc. Mass | mDa | PPM | DBE | i-FIT | Norm | Conf(%) | Formula |
| 406.0362 | 406.0402 | -4.0 | -9.9 | 16.5 | 389.5 | n/a | n/a | C23 H14 N O2 Cl2 |

HRMS Spectrum of EA3

Monoisotopic Mass, Even Electron Ions
1 formula(e) evaluated with 1 results within limits (up to 50 closest results for each mass)
Elements Used:
C: 23-23 H: 16-16 N: 1-1 O: 2-2 F: 1-1 Cl: 1-1
EA4 7 (0.085)
1: TOF MS ES+

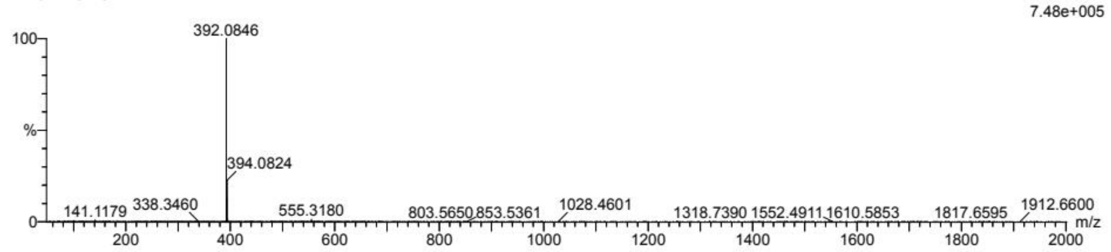

| | | | | |
|---|---|---|---|---|
| Minimum: | | | | -1.5 |
| Maximum: | | 5.0 | 50.0 | 50.0 |

| Mass | Calc. Mass | mDa | PPM | DBE | i-FIT | Norm | Conf(%) | Formula |
|---|---|---|---|---|---|---|---|---|
| 392.0846 | 392.0854 | -0.8 | -2.0 | 15.5 | 131.9 | n/a | n/a | C23 H16 N O2 F Cl |

HRMS Spectrum of EA4

Monoisotopic Mass, Even Electron Ions
1 formula(e) evaluated with 1 results within limits (up to 50 best isotopic matches for each mass)
Elements Used:
C: 24-24 H: 17-17 N: 1-1 O: 2-2 Cl: 1-1
EA6-0914 11 (0.083)
1: TOF MS ES-

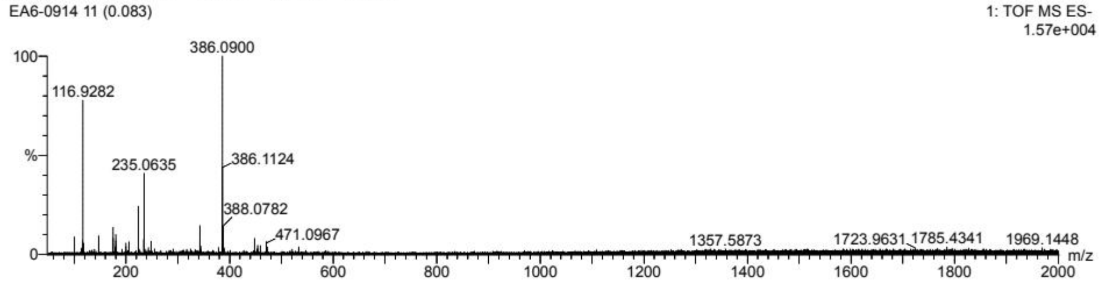

| | | | | |
|---|---|---|---|---|
| Minimum: | | | | -1.5 |
| Maximum: | | 5.0 | 10.0 | 50.0 |

| Mass | Calc. Mass | mDa | PPM | DBE | i-FIT | Norm | Conf(%) | Formula |
|---|---|---|---|---|---|---|---|---|
| 386.0900 | 386.0948 | -4.8 | -12.4 | 16.5 | 417.8 | n/a | n/a | C24 H17 N O2 Cl |

HRMS Spectrum of EA6

Monoisotopic Mass, Even Electron Ions
1 formula(e) evaluated with 1 results within limits (up to 50 closest results for each mass)
Elements Used:
C: 23-23 H: 15-15 N: 1-1 O: 2-2 F: 2-2 Cl: 1-1
EA8 7 (0.085)
1: TOF MS ES+

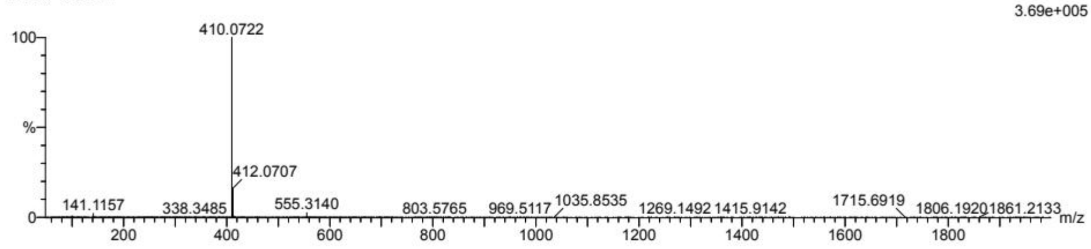

| | | | | |
|---|---|---|---|---|
| Minimum: | | | | -1.5 |
| Maximum: | | 5.0 | 50.0 | 50.0 |

| Mass | Calc. Mass | mDa | PPM | DBE | i-FIT | Norm | Conf(%) | Formula |
|---|---|---|---|---|---|---|---|---|
| 410.0722 | 410.0759 | -3.7 | -9.0 | 15.5 | 96.0 | n/a | n/a | C23 H15 N O2 F2 Cl |

HRMS Spectrum of EA8

Monoisotopic Mass, Even Electron Ions
1 formula(e) evaluated with 1 results within limits (up to 50 closest results for each mass)
Elements Used:
C: 24-24 H: 18-18 N: 1-1 O: 3-3 F: 2-2
FA11 5 (0.068)
1: TOF MS ES+

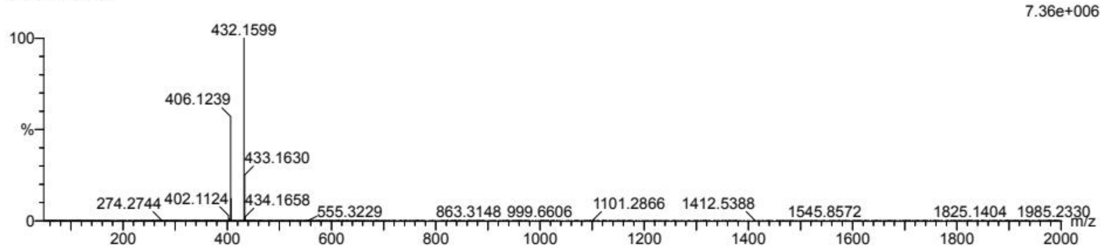

| | | | | | | | | |
|---|---|---|---|---|---|---|---|---|
| Minimum: | | | | -1.5 | | | | |
| Maximum: | | 5.0 | 50.0 | 50.0 | | | | |
| Mass | Calc. Mass | mDa | PPM | DBE | i-FIT | Norm | Conf(%) | Formula |
| 406.1239 | 406.1255 | -1.6 | -3.9 | 15.5 | 195.4 | n/a | n/a | C24 H18 N O3 F2 |

HRMS Spectrum of FA11

Monoisotopic Mass, Even Electron Ions
1 formula(e) evaluated with 1 results within limits (up to 50 closest results for each mass)
Elements Used:
C: 24-24 H: 18-18 N: 1-1 O: 3-3 F: 1-1 Cl: 1-1
FA12 5 (0.068)
1: TOF MS ES+

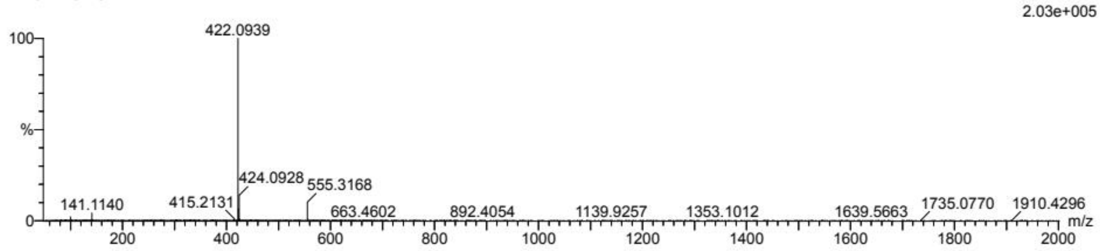

| | | | | | | | | |
|---|---|---|---|---|---|---|---|---|
| Minimum: | | | | -1.5 | | | | |
| Maximum: | | 5.0 | 50.0 | 50.0 | | | | |
| Mass | Calc. Mass | mDa | PPM | DBE | i-FIT | Norm | Conf(%) | Formula |
| 422.0939 | 422.0959 | -2.0 | -4.7 | 15.5 | 97.4 | n/a | n/a | C24 H18 N O3 F Cl |

HRMS Spectrum of FA12

Monoisotopic Mass, Even Electron Ions
1 formula(e) evaluated with 1 results within limits (up to 50 closest results for each mass)
Elements Used:
C: 25-25 H: 21-21 N: 1-1 O: 4-4 F: 1-1
FA14 5 (0.068)
1: TOF MS ES+

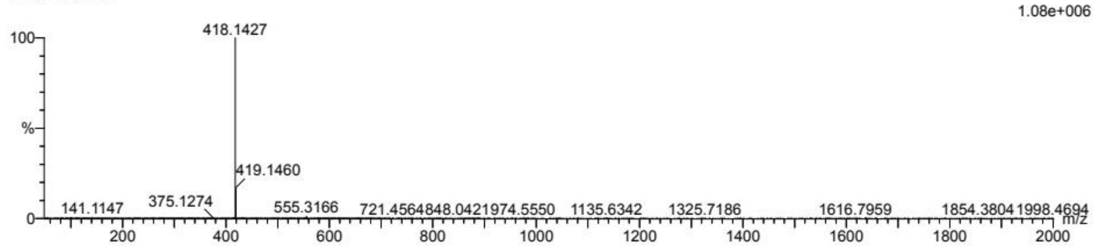

| | | | | | | | | |
|---|---|---|---|---|---|---|---|---|
| Minimum: | | | | -1.5 | | | | |
| Maximum: | | 5.0 | 50.0 | 50.0 | | | | |
| Mass | Calc. Mass | mDa | PPM | DBE | i-FIT | Norm | Conf(%) | Formula |
| 418.1427 | 418.1455 | -2.8 | -6.7 | 15.5 | 161.3 | n/a | n/a | C25 H21 N O4 F |

HRMS Spectrum of FA14

Monoisotopic Mass, Even Electron Ions
1 formula(e) evaluated with 1 results within limits (up to 50 closest results for each mass)
Elements Used:
C: 15-15   H: 13-13   O: 1-1   F: 2-2
H-A2 7 (0.085)
1: TOF MS ES+

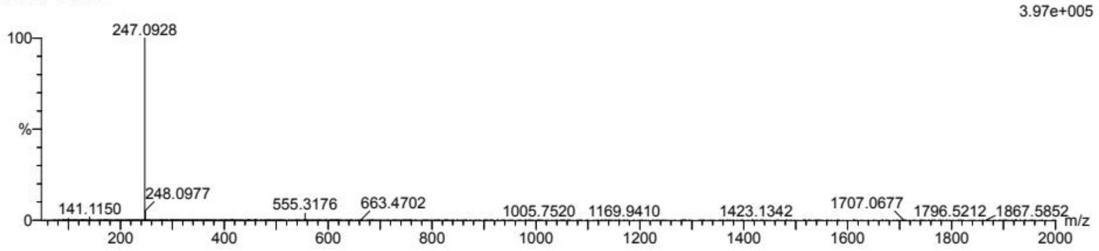

| | | | | | | | | |
|---|---|---|---|---|---|---|---|---|
| Minimum: | | | | -1.5 | | | | |
| Maximum: | | 5.0 | 50.0 | 50.0 | | | | |
| Mass | Calc. Mass | mDa | PPM | DBE | i-FIT | Norm | Conf(%) | Formula |
| 247.0928 | 247.0934 | -0.6 | -2.4 | 8.5 | 146.0 | n/a | n/a | C15 H13 O F2 |

HRMS Spectrum of A2

Monoisotopic Mass, Even Electron Ions
1 formula(e) evaluated with 1 results within limits (up to 50 closest results for each mass)
Elements Used:
C: 15-15   H: 14-14   O: 1-1   Cl: 1-1
H-A3 7 (0.085)
1: TOF MS ES+

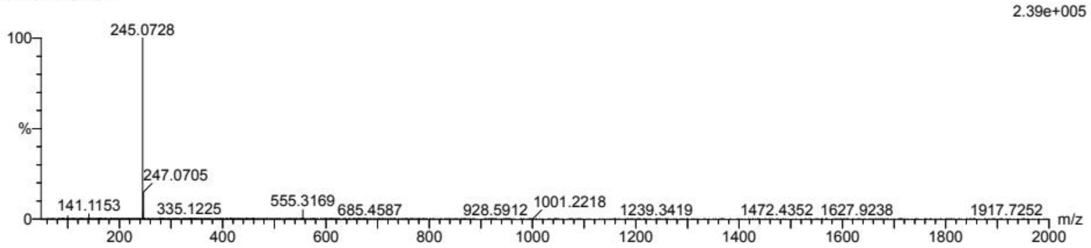

| | | | | | | | | |
|---|---|---|---|---|---|---|---|---|
| Minimum: | | | | -1.5 | | | | |
| Maximum: | | 5.0 | 50.0 | 50.0 | | | | |
| Mass | Calc. Mass | mDa | PPM | DBE | i-FIT | Norm | Conf(%) | Formula |
| 245.0728 | 245.0733 | -0.5 | -2.0 | 8.5 | 81.1 | n/a | n/a | C15 H14 O Cl |

HRMS Spectrum of A3

Monoisotopic Mass, Even Electron Ions
1 formula(e) evaluated with 1 results within limits (up to 50 best isotopic matches for each mass)
Elements Used:
C: 15-15   H: 14-14   O: 1-1   F: 1-1

H-A4 7 (0.085)

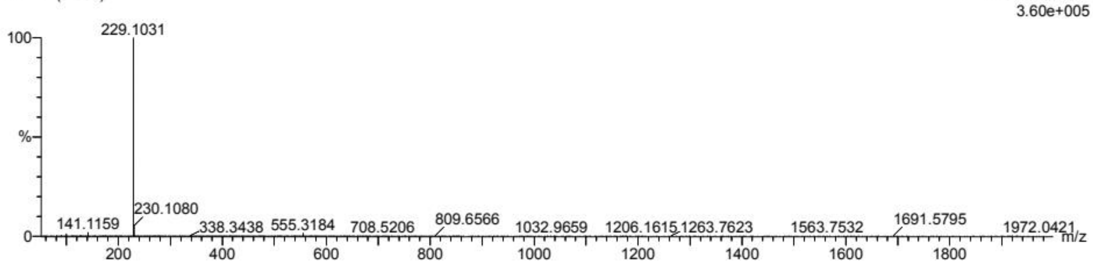

| | | | | | | | | |
|---|---|---|---|---|---|---|---|---|
| Minimum: | | | | -1.5 | | | | |
| Maximum: | | 5.0 | 10.0 | 50.0 | | | | |
| Mass | Calc. Mass | mDa | PPM | DBE | i-FIT | Norm | Conf(%) | Formula |
| 229.1031 | 229.1029 | 0.2 | 0.9 | 8.5 | 115.9 | n/a | n/a | C15 H14 O F |

HRMS Spectrum of A4

Monoisotopic Mass, Even Electron Ions
1 formula(e) evaluated with 1 results within limits (up to 50 closest results for each mass)
Elements Used:
C: 16-16 H: 17-17 O: 1-1
H-A6 7 (0.085)
1: TOF MS ES+

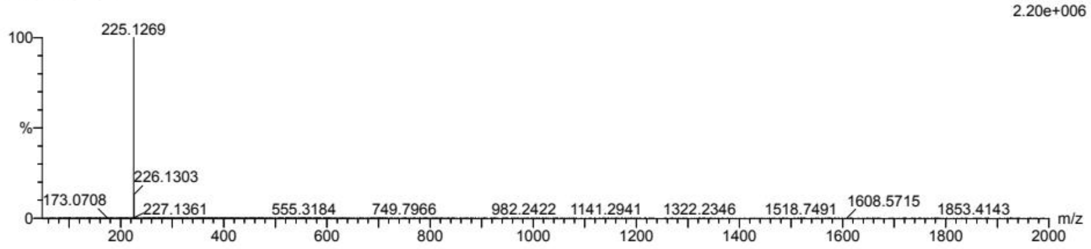

| | | | | | | | | |
|---|---|---|---|---|---|---|---|---|
| Minimum: | | | | -1.5 | | | | |
| Maximum: | | 5.0 | 50.0 | 50.0 | | | | |
| Mass | Calc. Mass | mDa | PPM | DBE | i-FIT | Norm | Conf(%) | Formula |
| 225.1269 | 225.1279 | -1.0 | -4.4 | 8.5 | 126.4 | n/a | n/a | C16 H17 O |

HRMS Spectrum of A6

Monoisotopic Mass, Even Electron Ions
1 formula(e) evaluated with 1 results within limits (up to 50 closest results for each mass)
Elements Used:
C: 15-15 H: 13-13 O: 1-1 F: 2-2

H-A8 7 (0.085)
1: TOF MS ES+

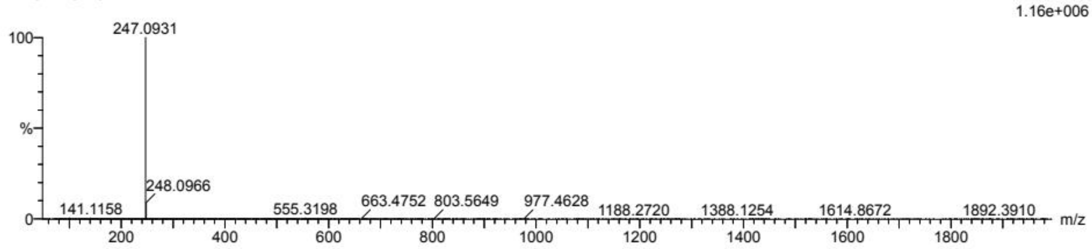

| | | | | | | | | |
|---|---|---|---|---|---|---|---|---|
| Minimum: | | | | -1.5 | | | | |
| Maximum: | | 5.0 | 50.0 | 50.0 | | | | |
| Mass | Calc. Mass | mDa | PPM | DBE | i-FIT | Norm | Conf(%) | Formula |
| 247.0931 | 247.0934 | -0.3 | -1.2 | 8.5 | 162.2 | n/a | n/a | C15 H13 O F2 |

HRMS Spectrum of A8

Monoisotopic Mass, Even Electron Ions
1 formula(e) evaluated with 1 results within limits (up to 50 closest results for each mass)
Elements Used:
C: 15-15 H: 13-13 O: 1-1 F: 2-2
H-A11 6 (0.077)
1: TOF MS ES+

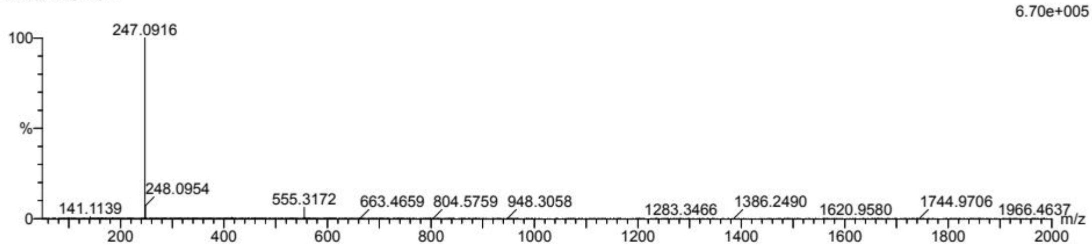

| | | | | | | | | |
|---|---|---|---|---|---|---|---|---|
| Minimum: | | | | -1.5 | | | | |
| Maximum: | | 5.0 | 50.0 | 50.0 | | | | |
| Mass | Calc. Mass | mDa | PPM | DBE | i-FIT | Norm | Conf(%) | Formula |
| 247.0916 | 247.0934 | -1.8 | -7.3 | 8.5 | 142.2 | n/a | n/a | C15 H13 O F2 |

HRMS Spectrum of A11

Monoisotopic Mass, Even Electron Ions
1 formula(e) evaluated with 1 results within limits (up to 50 closest results for each mass)
Elements Used:
C: 15-15 H: 13-13 O: 1-1 F: 1-1 Cl: 1-1
H-A12 7 (0.085)
1: TOF MS ES+

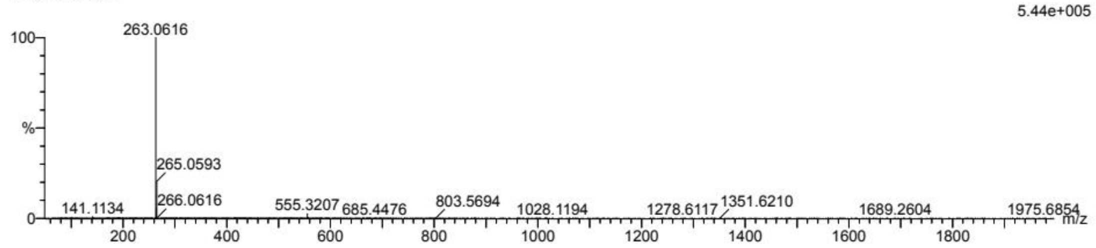

| | | | | | | | | |
|---|---|---|---|---|---|---|---|---|
| Minimum: | | | | -1.5 | | | | |
| Maximum: | | 5.0 | 50.0 | 50.0 | | | | |
| Mass | Calc. Mass | mDa | PPM | DBE | i-FIT | Norm | Conf(%) | Formula |
| 263.0616 | 263.0639 | -2.3 | -8.7 | 8.5 | 121.2 | n/a | n/a | C15 H13 O F Cl |

HRMS Spectrum of A12

Monoisotopic Mass, Even Electron Ions
1 formula(e) evaluated with 1 results within limits (up to 50 closest results for each mass)
Elements Used:
C: 16-16 H: 16-16 O: 2-2 F: 1-1
H-A14 6 (0.077)
1: TOF MS ES+

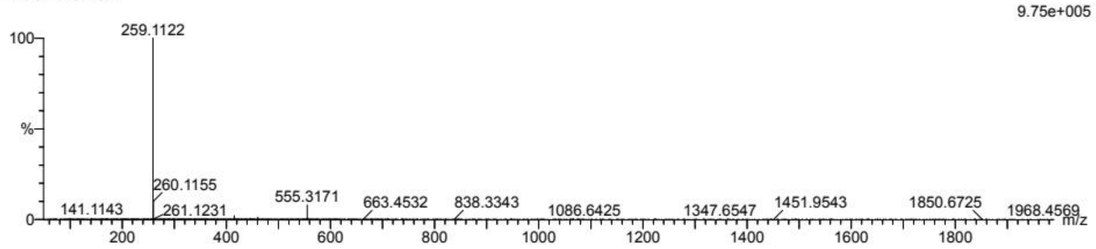

| | | | | | | | | |
|---|---|---|---|---|---|---|---|---|
| Minimum: | | | | -1.5 | | | | |
| Maximum: | | 5.0 | 50.0 | 50.0 | | | | |
| Mass | Calc. Mass | mDa | PPM | DBE | i-FIT | Norm | Conf(%) | Formula |
| 259.1122 | 259.1134 | -1.2 | -4.6 | 8.5 | 111.9 | n/a | n/a | C16 H16 O2 F |

HRMS Spectrum of A14
